# Signatures of adaptive evolution in platyrrhine primate genomes

**DOI:** 10.1101/2021.08.19.456368

**Authors:** Hazel Byrne, Timothy H. Webster, Sarah F. Brosnan, Patrícia Izar, Jessica W. Lynch

## Abstract

The family Cebidae (capuchin and squirrel monkeys) form a remarkable platyrrhine clade exhibiting among the largest primate encephalisation quotients. Each cebid lineage is characterised by notable lineage-specific traits, with capuchins showing striking similarities to Hominidae including high sensorimotor intelligence with tool use, advanced cognitive abilities, and behavioural flexibility. Here, we take a comparative genomics approach, analysing five cebid branches including successive lineages, to infer a stepwise timeline for cebid adaptive evolution. We uncover candidate targets of selection across various periods of cebid evolution that may underlie the emergence of lineage-specific traits. Our analyses highlight shifting and sustained selective pressures on genes related to brain development, longevity, reproduction, and morphology, including evidence for cumulative and diversifying neurobiological adaptations over cebid evolutionary history. In addition to generating a new, high-quality reference genome assembly for robust capuchins, our results lend to a better understanding of the adaptive diversification of this distinctive primate clade.

## Introduction

Platyrrhine primates (also known as Neotropical primates) present a striking example of the adaptive diversification of a primate clade into diverse ecological niches. Platyrrhines of South and Central America and catarrhines of Africa and Asia (and extinct forms from Europe) likely diverged via transatlantic dispersal of the platyrrhine ancestor from Africa to South America 40 to 44 million years ago (mya), with the earliest South American fossils resembling small Eocene African anthropoids (*1, 2*). The crown platyrrhine radiation began to diversify 20 to 25 mya, with most of the extant diversity contained in the rainforests of Amazonia and the Atlantic Forest biome (*1, 3*). It has been suggested that ecological opportunity across multidimensional niches in expanding rainforest environments may have driven the diversification of major platyrrhine lineages, leading to the evolution of a plethora of forms with over 20 extant genera and 170 extant species (*4, 5*). Platyrrhine primates show striking phenotypic diversity in body and brain size, skeletal morphology, pelage patterns, group size, social and mating systems, life history and longevity, behavioural plasticity, diet and dietary adaptations, among many other traits, with this diversity becoming increasingly well characterised in recent years. We know very little, however, about the genetic changes involved in the evolution of this incredible array of diversity and where in the platyrrhine clade those changes occurred.

While all major platyrrhine groups show lineage-specific traits, the family Cebidae (capuchin and squirrel monkeys, following (*6*) and current IUCN Red List Taxonomy) are compelling considering their large encephalisation quotient (EQ; relative brain to body size), with reconstructions showing one of the fastest increases in EQ across primates along the ancestral Cebidae branch (*7*). Capuchins (subfamily Cebinae) are a particularly remarkable platyrrhine clade with many striking similarities to Hominidae including social conventions and traditions, complex relationships, high dexterity, sensorimotor intelligence with tool use and extractive foraging, advanced derived cognitive abilities, diverse behavioural repertoire and flexibility, and slow maturation (*8*). These traits are uncommon or absent among other platyrrhines and it is of great anthropological interest to gain insight into the evolutionary mechanisms underlying the independent emergence of these convergent traits and their associated genomic changes. The existence of two capuchin lineages (gracile and robust) with both shared and derived traits (including differences in cranial and post-cranial skeletal morphology, tool use, social and sexual behaviours, etc.), which diverged within a similar timeframe to *Homo* and *Pan*, brings further interest to understand their distinct evolutionary trajectories. Squirrel monkeys (genus *Saimiri*), the sister group to capuchins, are also characterised by slow maturation and large EQ, but lack other parallels to apes and humans described above for capuchins. Squirrel monkeys are hyper-gregarious with the largest stable social groups among platyrrhines, and frequently engage in mixed-species associations, especially with capuchins (*9*). Squirrel monkeys are also a key primate biomedical model with foci on neuroendocrinology, ophthalmology, pharmacology, behaviour, viral persistence, infectious diseases, cancer treatment, and reproductive physiology, among others (*10*).

Here, we take a comparative genomics approach to uncover signatures of adaptive evolution in cebid genomes to better understand the adaptive diversification of this distinctive platyrrhine primate clade. We focus on the three extant cebid lineages—robust capuchins (genus *Sapajus*), gracile capuchins (genus *Cebus*), and squirrel monkeys (genus *Saimiri*)—as well as the ancestral capuchin (Cebinae) and ancestral Cebidae branches. Through the analysis of successive lineages in the platyrrhine phylogeny, we are able to infer a stepwise timeline for cebid adaptive evolution and identify candidate adaptive genes that may underlie the emergence of lineage-specific traits. Previous work assessing signatures of adaptive evolution in protein-coding regions for cebid lineages considered the entire capuchin subfamily (Cebinae) together as represented by a single species (*Cebus imitator*), uncovering broad signatures of positive selection on the brain and DNA repair (which was associated with longevity) (*11*), or focused on signatures of convergence among encephalised primate lineages including humans (*12*). This work greatly expands upon these existing studies by individually analysing five distinct cebid branches to infer the targets of selection during various time periods of cebid evolution.

## Results

### Robust capuchin reference genome

At the start of this study, annotated genome assemblies were publicly available for *Cebus imitator* and *Saimiri boliviensis*. We generated a new genome assembly for *Sapajus apella* using short-read data (∼148-fold coverage) scaffolded with Dovetail’s Chicago proximity ligation libraries (Table S1) using their HiRise pipeline (*13*). Total length of this genome assembly was 2,520 Mbp (in 6631 scaffolds) with an N50 of 27.1 Mbp (29 scaffolds) and N90 of 4.04 Mbp (116 scaffolds). We identified 91.5% (5,666) of BUSCO’s (*14*) Euarchontoglires-specific conserved single-copy orthologs in the assembly including 85% (5,264) complete (with 0.6% duplicated) and 6.5% (402) fragmented; and 90.3% (224) of CEGMA’s (*15*) core eukaryotic genes (CEGs). Together, assembly metrics and genome completeness based on gene content indicate a contiguous, high-quality reference genome assembly for robust capuchins (*Sapajus*). We estimated genome size with filtered short read data based on *k*-mer (31-mer) frequencies using the four approaches resulting in an estimated haploid genome length for our *S. apella* reference individual between 2,918 and 3,029 Mbp (Table S2). Previous estimates of genome size for other robust capuchin species, *Sapajus libidinosus* and *Sapajus nigritus*, estimated using Feulgen image analysis densitometry, ranged between 3,276-3,374 Mbp and 2,921-3,025 Mbp, respectively (*16*). These estimates, in particular for *S. nigritus*, are very similar to our estimates for *S. apella* calculated in this study.

We pooled raw RNAseq data (367 million read pairs) derived from total RNA from 17 tissues from the same reference individual and, post-filtering, retained 341 million read pairs (Table S3). We assessed quality metrics and completeness of the seven transcript assembles generated using cleaned RNAseq read pairs with rnaQUAST (*17*) and BUSCO, which revealed that upwards of 94% of the transcripts aligned to the genome with an average aligned percentage of greater than 92.7%, and indicated the final assemblies used in downstream analyses (TrinDNv2, PASAv1, and NRv1) were high-quality, near complete transcriptomes (∼96 to 97% complete) (Table S4). Repeat annotation of the genome assembly using libraries of both known and *de novo* elements estimated the total interspersed content of the genome as 43.02% (1.06 Gbp), and total annotated repeat content (including transposable elements as well as small RNA, satellites, simple repeats, and low complexity repeats) as 44.63% (1.12 Gbp) (Table S5). After three iterations of Maker (*18, 19*) to predict and annotate gene models in the robust capuchin genome assembly (Table S6) and subsequent filtering, we recovered 25,279 predicted genes for *S. apella* for downstream analyses.

### Ortholog alignment & branch/branch-site model tests

We initially identified 12,160 one-to-one orthologs recovered in at least two of the ten species we used in our comparative genomic analyses; these include nine primates, of which four are platyrrhines, and mouse (Figure 1; Table S7). After filtering for a minimum of five species and the presence of at least one capuchin lineage (gracile or robust), alignment using Guidance2 (*20*) with 100 bootstraps, and filtering for errors to reduce the likelihood of false positives, we retained a set of 9,216 conservative, manually-curated CDS alignments which were highly likely to represent one-to-one ortholog groups across their length. Detailed information on each of the final alignments, including group ID, assigned gene symbol, and Entrez ID can be found in Table S8. In total, there were 207 different combinations of species (species sets) represented in the final alignments (Table S9), with most alignments assigned to the set of all species (full) (N = 4,636) or sets with nine species (N = 2,819), and the rest to sets with between eight and five species (N = 1,761) (Tables 1, S10).

**Figure 1.**
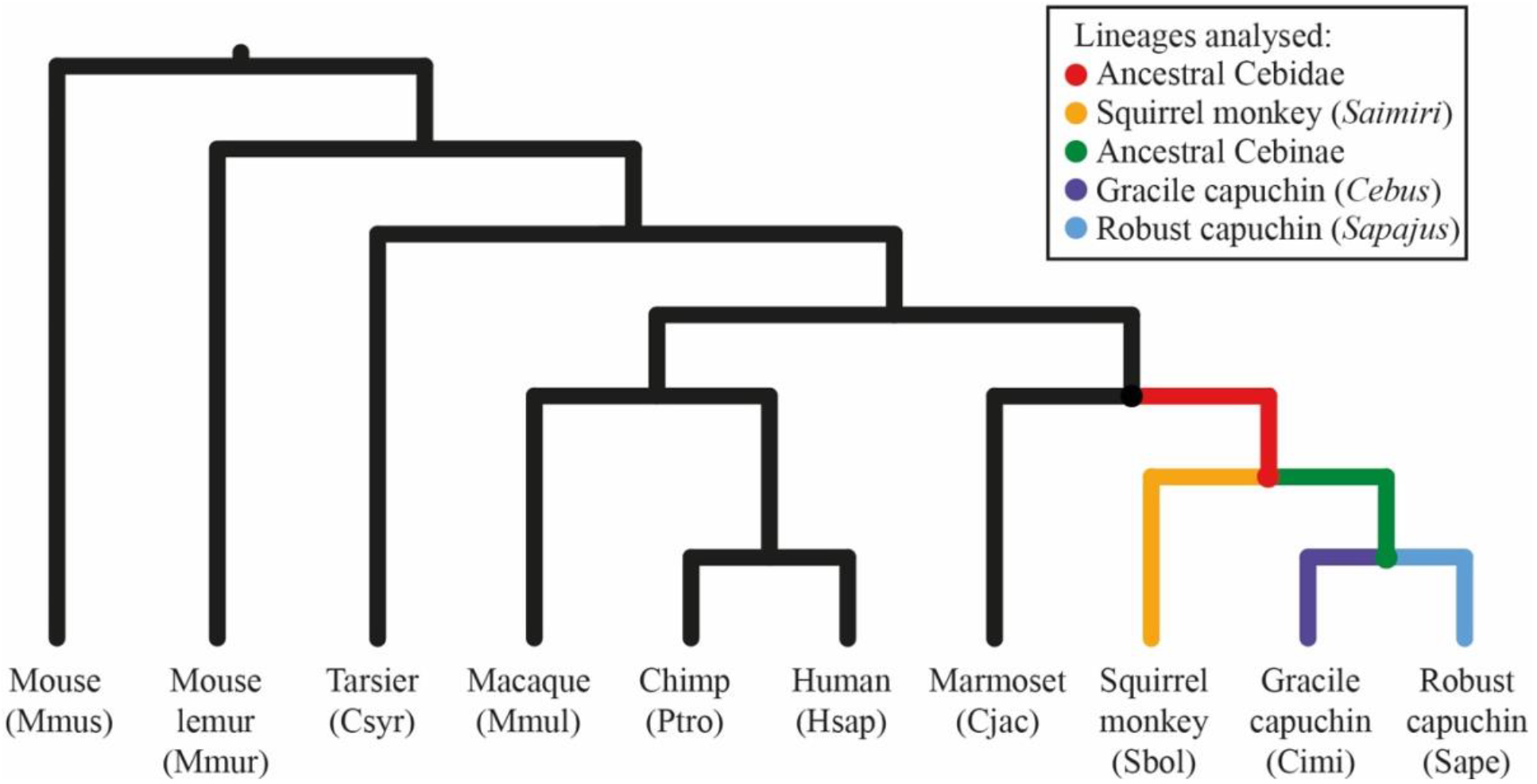
Phylogenetic reconstruction showing the consensus guide tree topology and the cebid branches assessed for signatures of positive selection. H3a is not shown, but includes branches for Cimi, Sape, and their ancestor.

**Table 1.**
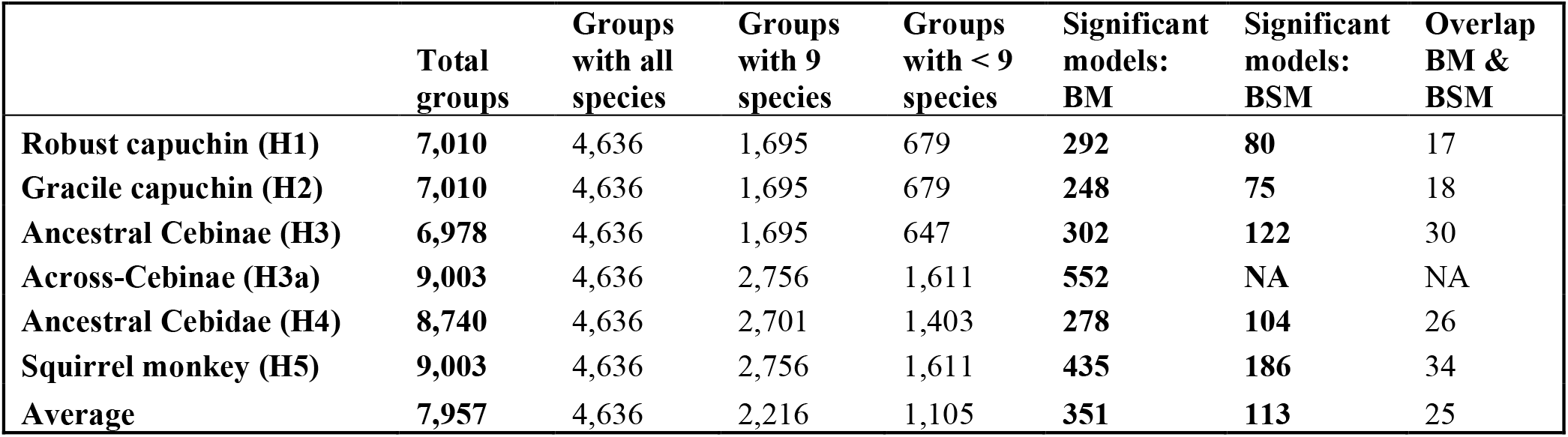
Counts of groups and significant results for BM and BSM tests with PAML.

|  | Total<br>groups | Groups<br>with all<br>species | Groups<br>with 9<br>species | Groups<br>with < 9<br>species | Significant<br>models:<br>BM | Significant<br>models:<br>BSM | Overlap<br>BM &<br>BSM |
| --- | --- | --- | --- | --- | --- | --- | --- |
| **Robust capuchin (H1)** | **7,010** | 4,636 | 1,695 | 679 | **292** | **80** | 17 |
| **Gracile capuchin (H2)** | **7,010** | 4,636 | 1,695 | 679 | **248** | **75** | 18 |
| **Ancestral Cebinae (H3)** | **6,978** | 4,636 | 1,695 | 647 | **302** | **122** | 30 |
| **Across-Cebinae (H3a)** | **9,003** | 4,636 | 2,756 | 1,611 | **552** | **NA** | NA |
| **Ancestral Cebidae (H4)** | **8,740** | 4,636 | 2,701 | 1,403 | **278** | **104** | 26 |
| **Squirrel monkey (H5)** | **9,003** | 4,636 | 2,756 | 1,611 | **435** | **186** | 34 |
| **Average** | **7,957** | 4,636 | 2,216 | 1,105 | **351** | **113** | 25 |

These 9,216 alignments were used as input to our codon-based models of evolution based on non-synonymous versus synonymous substitutions (ω or dN/dS ratio) to identify candidate genes under selection in six cebid lineages of interest (Figure 1); (H1) robust capuchin (*Sapajus*); (H2) gracile capuchin (*Cebus*); (H3) ancestral Cebinae (capuchins); (H3a) across-capuchins (all Cebinae; branches H1, H2, and H3 combined); (H4) ancestral Cebidae (i.e., ancestor to capuchins and squirrel monkeys); and (H5) squirrel monkey (*Saimiri*).

In total we tested 86,485 models for 11 lineage and test combinations using codeml from PAML (*21*); 47,744 branch models (BM) across all six lineages, which tests for elevated dN/dS ratios along the target branch indicating accelerated evolution; and 38,741 branch-site models (BSM) across five lineages (excluding H3a), which tests for episodic selection by searching for positively selected sites in the target lineage. Groups (alignments) analysed per lineage varied between 6,978 and 9,003 of 9,216 total (Tables 1, S10), with averages of 7,957 BM and 7,748 BSM tests. Across the six lineages analysed for BM, we recovered 248 to 552 (avg. 351) models with significant signatures of accelerated evolution. In contrast, across the five lineages analysed for BSM, we found 75 to 186 (avg. 113) models with significant signatures of episodic positive selection, much fewer than for BM tests particularly for shorter branches for the capuchin lineages. Between 17 and 34 (avg. 25) groups are significant for both BM and BSM tests for the same lineage (Tables 1, S10). Lists of all groups (genes) analysed for each of the six lineages, along with significance for BM and/or BSM tests, can be found in Tables S11–S16. More detailed information for the groups with significant evidence of accelerated evolution or episodic selection from the BM and/or BSM tests including p-value, LRT statistic, and likelihood scores is located in Tables S17– S22.

### Gene set enrichment

Gene set enrichment analyses using DAVID v.6.8 (*22*) for each set of significant genes from each combination of lineage and test (six BM, five BSM) aided interpretation of the biological significance of the results. We assessed lists of enriched BP (biological process), CC (cellular component), and MF (molecular function) gene ontology (GO) terms, UP keywords, KEGG pathways, Reactome pathways, and disease annotations, as well as functional annotation clustering across the three GO terms together under the high classification stringency criteria, with an EASE score of < 0.05 required for all enriched annotated terms. Across all lineages for the BM and BSM gene sets, we recovered between 2 to 13 (avg. 6) and 0 to 9 (avg. 3) GO clusters, and 68 and 189 (avg. 103) and 10 to 123 (avg. 60) enriched terms (all annotation categories), respectively (Table S23). Information on each of the enriched annotated terms and GO clusters including description, gene counts and hits, and statistical results such as EASE score and fold enrichment, for each gene set enrichment analysis are found in Tables S24–S44. We briefly summarise the gene set enrichment results in Table 2. A more detailed written summary of the gene set enrichment results for each lineage is presented in the supplementary materials.

**Table 2.**
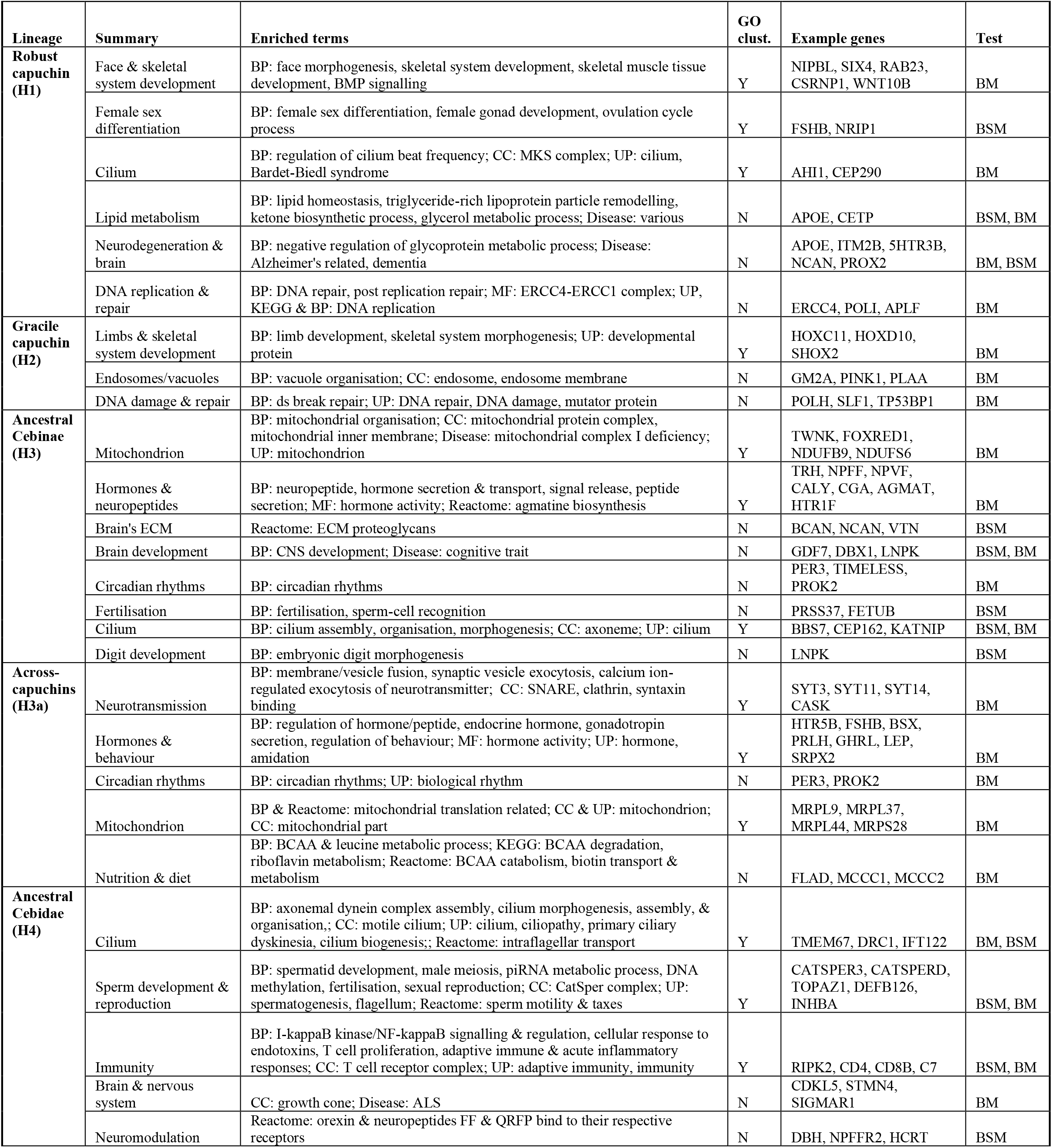
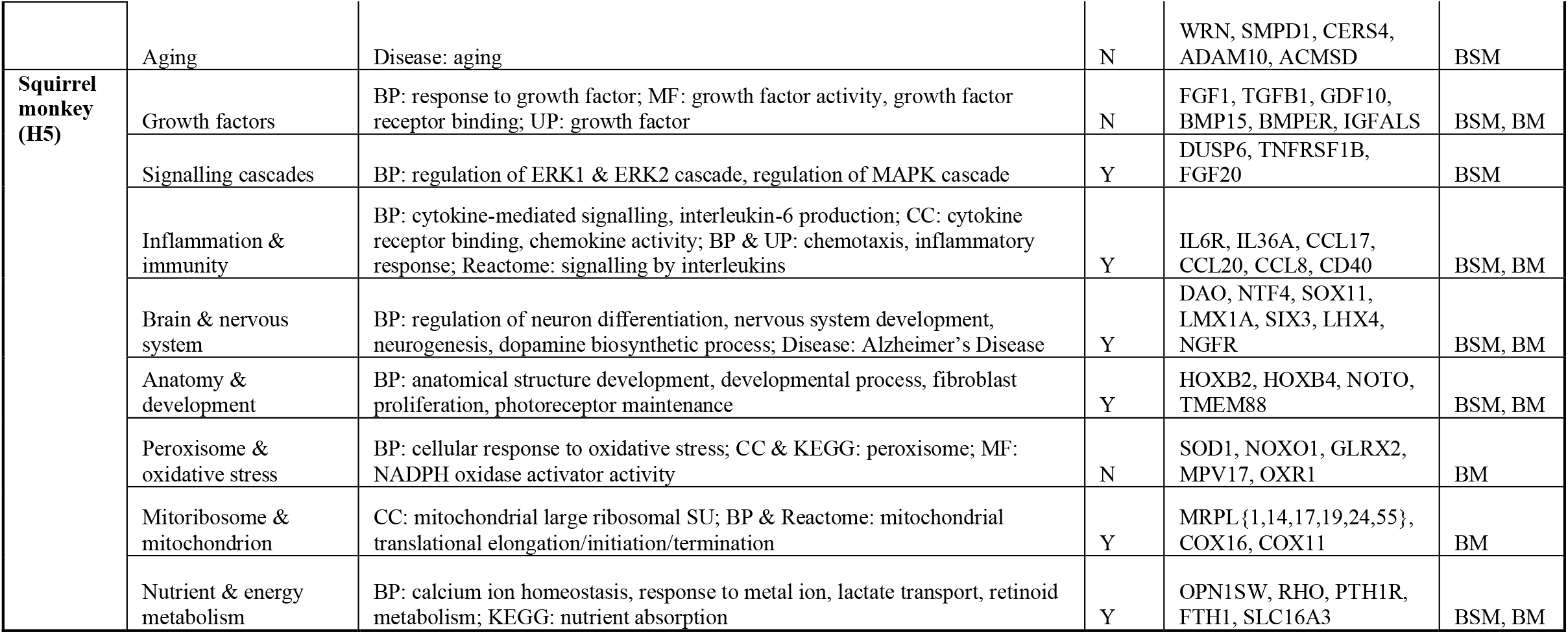
Summary of gene set enrichment results for each cebid lineage including enriched terms, example genes, and test.

## Discussion

Our analyses reveal signatures of positive selection on many lineage-specific traits across Cebidae and highlight branches with strong selective pressure on genes related to brain development and function, longevity, behaviour, reproduction, and morphology (Figure 2). Perhaps most striking are the sustained signatures of positive selection on brain evolution across Cebidae, which appear early in cebid history with subsequent selection on different aspects of central nervous system (CNS) development at various time intervals for different lineages. While we recover an evolutionary trajectory of encephalisation beginning in ancestral Cebidae and continuing independently in squirrel monkeys and capuchins, the strongest evidence for selection on neuroplasticity, behavioural flexibility, and manual dexterity is found for ancestral Cebinae or the entire capuchin clade when considered together (across-capuchins). The most striking signatures of selection recovered independently for the capuchin genera relate to their body shape and skeletal morphology, including the distinctive robust cranial and skeletal morphology in robust capuchins (*Sapajus*), and, conversely, the gracile limb morphology associated with more rapid, agile movement in gracile capuchins (*Cebus*). All three extant cebid genera are long-lived for their body size, and each shows independent signatures of selection on genes related to aging, longevity, and/or neurodegeneration. In addition, in contrast to other closely related taxa, all three cebid genera live in relatively large groups with polygynandry and complex sexual interactions, and we recover signatures of sustained positive selection related to sperm production/morphology and reproductive behaviour. Our comparative approach to uncovering candidate targets of positive selection within Cebidae highlights shifting and sustained selective pressures within this clade, including evidence for cumulative and diversifying neurobiological adaptations over cebid evolutionary history. In the following sections, we discuss our results describing adaptive evolutionary change in these lineages across various biological categories.

**Figure 2.**
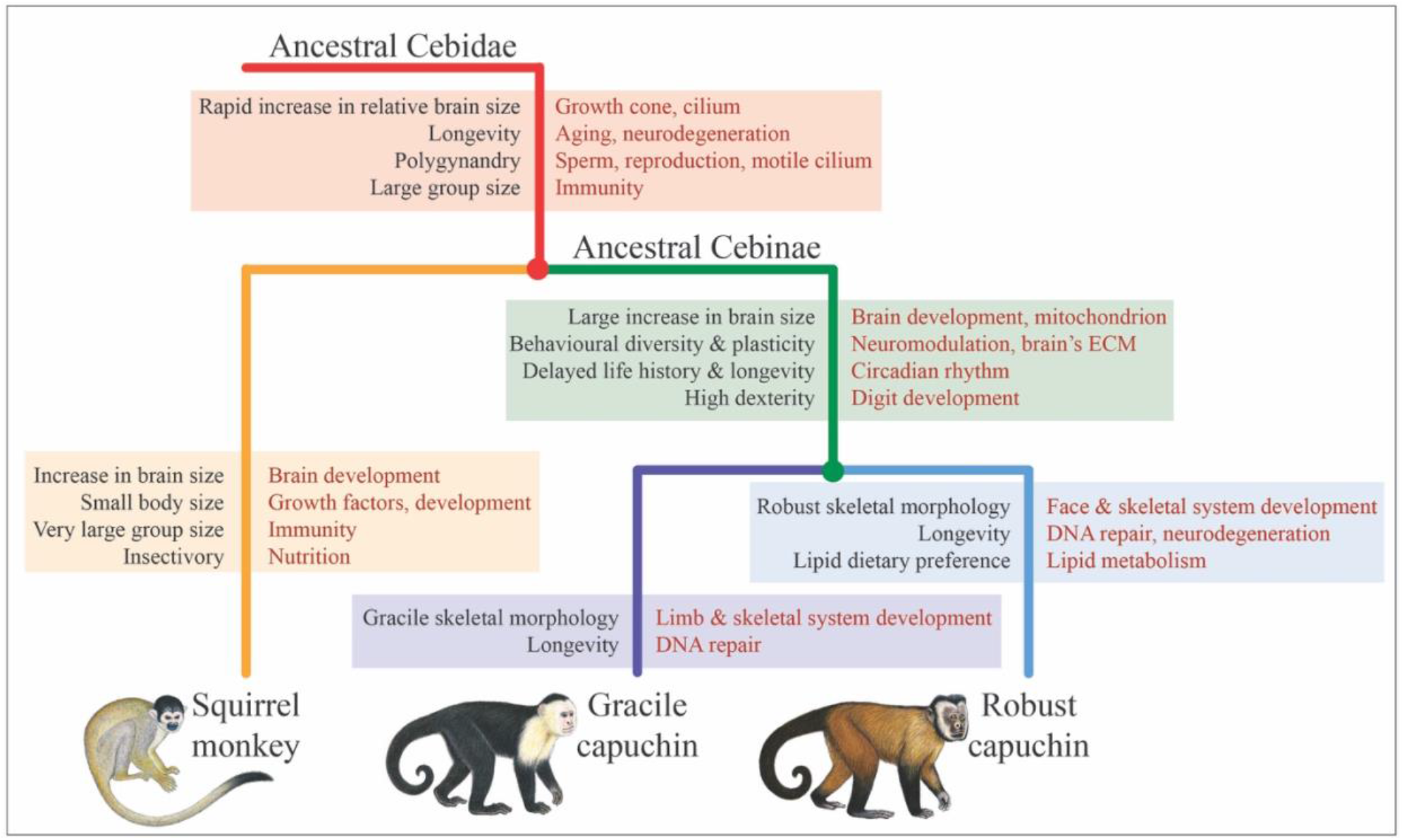
Graphic summary of select signatures of adaptive evolution per cebid lineage. On the left are traits associated with each branch, which for the ancestral lineages are inferred based on traits shared by all daughter lineages. Brain size changes are based on (*7*). On the right (in red text) are the associated signatures of adaptive evolution. H3a is not shown. Illustrations by Stephen Nash ©.

### Neurodevelopment and plasticity

A major hallmark of primate evolution is expansion of the brain, with numerous independent shifts to larger brain mass relative to body size also occurring among different primate lineages. Larger brains have long been associated with increased cognitive capabilities, higher social complexity, and increased ability to respond to environmental and socioecological challenges (*23*). The most encephalised primates after humans are platyrrhines of the family Cebidae— capuchins and squirrel monkeys (*11, 24*)—and ancestral state reconstructions have indicated that the second fastest increase in the rate of encephalisation across primates occurred along the ancestral Cebidae branch (*7*). Overall, our results are consistent with an evolutionary trajectory of encephalisation and adaptive brain evolution beginning in ancestral Cebidae and continuing independently in both squirrel monkeys and capuchins after their divergence around 13.8 million years ago (*25*).

We recovered signatures of selection and accelerated evolution on the CNS that may be associated with this encephalisation shift—in particular, related to brain development and patterning. For the ancestral Cebidae branch, we recovered the enriched CC term “growth cone”, a motile, sensory structure that plays a critical role in precisely specified brain wiring patterns, guiding axons to their targets during neural development, and is also essential in the mature brain for plasticity-dependent synaptogenesis (*26*). Growth cone dynamics and axonal tract development are regulated by ciliary signalling (*27*), and, notably, some of the strongest selective signatures for ancestral Cebidae are related to the cilium (both primary and motile, as well as microtubules). Many of the most enriched terms are cilium-specific and including a suite of genes with essential roles in ciliogenesis and implicated in the ciliopathy Joubert’s syndrome. Primary cilia are found in almost all mammalian cells and the range of symptoms characterising ciliopathies highlights the difficulty in associating the signal of selection on cilium with a single adaptive function; it is notable, however, that many of these disorders are characterised by pronounced neurodevelopmental abnormalities. Primary cilia are critical to the development of the CNS, playing essential roles in early patterning, neurogenesis, and neuronal migration and connectivity, at least in part owing to their essential role in mediating signal transduction in key signalling pathways (*28*). Taken together, these results suggest adaptive evolution of the CNS and brain patterning in ancestral Cebidae, which may be linked to the increase in brain size found along this branch.

After the Cebinae/Saimirinae divergence, relative brain size of both squirrel monkeys and capuchins is modelled to have increased independently at a similar rate. For squirrel monkeys, it is explained by a reduction in body size and moderate increase in brain size, while for capuchins it is driven by a large increase in brain size along with a smaller increase in body size (*7*). In agreement with this, we find continued brain-related signatures of selection in both lineages. For squirrel monkeys, we recovered various enriched brain-related GO terms for the BSM gene set including “regulation of neuron differentiation”, “nervous system development”, and “neurogenesis”, among others. A gene in the squirrel monkey gene set is *ADCYAP1*, which is accelerated in humans and has been associated with human brain size evolution (*29*). Some of these signatures for squirrel monkeys may also relate to the adaptive maintenance of a large brain size while reducing body size.

Capuchins are particularly notable for their large brains and high EQs, the latter second only to humans among primates (*30*), and other hallmarks of their evolution include their derived cognitive abilities, sensorimotor intelligence, diverse behavioural repertoire, and extensive behavioural plasticity (*8*). Capuchins show striking convergence with great apes (particularly humans) across these traits, which are uncommon among other platyrrhines. Related to these traits, we recover the enriched BP GO term “CNS development” (BSM gene set) for ancestral Cebinae with important developmental genes such as *GDF7*, which contributes to neuronal cell identity in the developing embryonic nervous system. As with ancestral Cebidae, we also find signatures of enrichment related to cilia for ancestral Cebinae (in both gene sets) with several genes involved in primary cilium function that are also found in the “CNS development” GO term (such as *CEP162* and *BBS7*) and implicated in ciliopathies including Seckel and Bardet-Biedl syndrome. Orkin et al. (*11*) also found signatures of adaptive evolution related to brain development and neurogenesis for *Cebus imitator*. Importantly, however, our study places these positive selection pressures for brain development as most strongly affecting the ancestral Cebidae and Cebinae lineages; this suggests that brain organisation and function may have become relatively stable with only minor divergence across the two capuchin genera despite their subsequent divergent ecological and morphological adaptations (but see below for some brain-related genes of interest).

Behavioural repertoires are manifestations of neural activity and changes in behaviour are ultimately followed by alterations in neuronal connectivity i.e., neuroplasticity (*31*). We found further brain-related signatures for capuchins putatively associated with this trait. Two highly ranked genes in the “CNS development” GO term for ancestral Cebinae encode chondroitin sulphate proteoglycans (CSPGs) of the lectican family that are specifically expressed in the CNS: *NCAN* (neurocan), the 4^th^ ranked gene in the BSM gene set, and *BCAN* (brevican), the 2^nd^ ranked gene in the BM gene set and also in the BSM gene set. These CSPGs serve as guidance cues during brain development, as well as play important roles in neuroplasticity by modulating synaptic connections in the adult brain. They are abundant components of the brain’s extracellular matrix, forming condensed lattice-like structures known as perineuronal nets (PNNs) that form as one of the ultimate acts coinciding with the closure of critical periods for experience-dependent plasticity. The relationship between neurons and PNNs is a central mechanism controlling neuroplasticity, with PNNs playing many important roles in CNS functions including regulating synaptic plasticity, stabilising synapses, and neuroprotection. They are involved in cognition through encoding, maintaining, and updating memories, as well as recovery after nervous system damage, psychiatric disease, and neurodegeneration (*32, 33*). It is therefore significant that two of the most central and abundant components of PNNs, the CSPGs brevican (*BCAN*) and neurocan (*NCAN*), show strong signatures of selection in ancestral Cebinae, with *BCAN* also selected in *Cebus* and *NCAN* also selected in *Sapajus*. Indeed, signatures of selection potentially related to synaptic plasticity appeared even earlier along the ancestral Cebidae branch given the importance of the growth cone for plasticity-dependent synaptogenesis, as discussed.

For the across-capuchin gene set, we recover strong signatures related to neurotransmission and vesicle fusion including six genes encoding synaptotagmin and synaptotagmin-like proteins which are known to play important roles in regulated neurotransmitter release and hormone secretion. Among these genes is *SYT11*, which forms an essential component of a neuronal vesicular trafficking pathway crucial for development and synaptic plasticity, and plays an important role in dopamine transmission (*34*). Other related genes for the across-capuchin gene set are involved in the regulation of synaptic AMPA receptors, which play a key role in synaptic plasticity being involved in long-term potentiation and depression of synaptic transmission in the hippocampus; and encoding or interacting with neurexins, neuronal cell surface proteins involved in synaptic contacts and transmission.

Although the brain-related signatures are strongest for the ancestral cebid lineages, we also find distinct significantly accelerated genes related to neurodevelopment in each of the capuchin genera suggesting some, perhaps minor, continuation of adaptive brain evolution independently in robust and gracile capuchins after their divergence around 5 to 6 million years ago. This signature is more notable for robust capuchins; we recover enriched terms related to cilia including the UP keywords “cilium” and “Bardet-Biedl syndrome”, and the CC GO term “MKS complex,” covering three genes involved in ciliogenesis and required for the formation of primary non-motile cilium. One of these is *AHI1*, which is required for both cerebellar and cortical development in humans, and may play a crucial role in ciliary signalling during cerebellum embryonic development as a positive modulator of classical Wnt signalling (*35*). *AHI1* also shows an accelerated rate of evolution along the human lineage since the split from chimpanzees and bonobos (*36*). For gracile capuchins, the enriched UP keyword “developmental protein” contains multiple genes with important roles in CNS development.

### Mitochondria and energy metabolism

The brain is one of the most metabolically expensive organs in the vertebrate body and large brains are, therefore, an evolutionarily costly adaptation (*37*). Tissues with high energy requirements, such as the brain, are highly dependent on mitochondria with hundreds to thousands within a single neuron (*38*). Signatures of adaptive evolution in nuclear-encoded mitochondrial genes have been found in large brained/encephalised mammals including the elephant and anthropoid primates generally (*39*–*41*), as well as in bats which have a high energy demand owing to flight (*42*). Mitochondria also play many important roles in the nervous system including in neurotransmitter metabolism, neurogenesis, neuroplasticity, and nervous system development, and are strongly implicated in aging (*43, 44*). We find recurrent signatures of selection on the mitochondrion in multiple cebid lineages. This signature is the strongest in ancestral Cebinae with recurrent, sweeping signatures across many annotation categories in the BM gene set including specific enriched terms related to the mitochondrial inner membrane and protein complexes which underlie the role of the mitochondrion as the cell’s powerhouse. The ancestral capuchin branch is where absolute brain volume shows both the greatest total increase and the fastest rate of increase among Cebidae branches (*7*), supporting the putative relationship between this signature, encephalisation, and the high energy requirements of large brains.

We also recover enriched broad mitochondrial terms for across-capuchins and ancestral Cebidae shared with the ancestral Cebinae branch, as well as additional signatures shared between across-capuchins and squirrel monkeys specific to nuclear-encoded mitochondrial ribosomal proteins (which form mitoribosomes) and the translation of essential mitochondrial mRNAs. Although signatures are sometimes shared across lineages, the genes involved usually differ; for example, there are four and six nuclear-encoded mitochondrial ribosomal genes for the across-capuchins and squirrel monkey branches, respectively, but with no overlapping genes. These results further support an evolutionary trajectory of encephalisation initiating in ancestral Cebidae, and continuing independently in capuchins and squirrel monkeys.

### Longevity, aging, and neurodegeneration

The cognitive advantages of a large brain should have an adaptive impact by reducing mortality thereby allowing selection to favour a longer life (*45, 46*). In this regard, the parallels between humans and capuchins are striking: capuchin monkeys have slow maturation and extended juvenescence reaching maturity at around 8 to 10 years old. Capuchin monkeys are among the most long-lived primates, reaching over 50 years in captivity, though life expectancy is thought to be much lower in the wild (*47, 48*) (Perry, pers. comm.). Consistent with this, we recovered broad signatures of selection on aging and related processes across various capuchin branches.

The maintenance of genomic stability is considered a major factor underlying human longevity with the accumulation of macromolecular damage, such as DNA damage, one of the most significant factors contributing to aging (*49*). We recover signatures of selection on DNA damage and repair related genes for both the robust and gracile capuchin branches independently, including enriched terms such as “double strand break repair”, “cellular response to DNA damage stimulus”, “DNA repair”, and “DNA damage”. These results support those of Orkin et al. (*11*), who also found signatures of selection related to DNA repair and damage in *Cebus imitator*. Aging is also associated with a decline in mitochondrial function with strong links between mitochondria and a wide range of processes associated with aging including senescence and inflammation (*44*). As discussed in a previous section, there are strong signatures of selection on mitochondria across cebid branches.

Squirrel monkeys are also long-lived primates when considering their small body size— around 30 years in captivity (*47*)—and thus selective pressure on longevity may have arisen along the ancestral Cebidae branch. Indeed, for ancestral Cebidae, we found the enriched disease annotation “aging” with several genes implicated in age-related neurodegeneration. A particularly notable gene in this annotation is *WRN* which plays a major role in genome stability with mutations in *WRN* associated with defective telomere maintenance and causing Werner syndrome, which is characterised by rapid onset of cellular senescence, early cancer onset, and premature aging (*50*). In addition, there are several important genes with signatures of selection related to sphingolipid and ceramide metabolism for ancestral Cebidae including a ceramide synthase (*CERS4*), and *SMPD1*, which encodes a lysosomal acid sphingomyelinase (ASM). Recent studies have highlighted the importance of ASM as a critical mediator for pathologies in aging and age-related neurodegenerative diseases, with ASM viewed as a promising drug target for anti-aging and the treatment of age-related neurodegenerative diseases (*51*). We also find another important ceramide synthase gene, *CERS*, in the across-capuchin gene set, which catalyses the synthesis of C18-ceramide in brain neurons, with elevated expression of this gene associated with increased longevity in humans (*52*).

Humans are particularly susceptible to age-related neurodegenerative disorders such as Alzheimer’s disease (AD). While non-human primates show some age-related neurodegeneration, pathological neurodegeneration such as seen in AD is rare (*53*). Interestingly, we recover many genes across the capuchin branches associated with age-related neurodegenerative disorders in humans. This is particularly evident for robust capuchins with various enriched disease annotations related to AD (also found for squirrel monkeys) and dementia, as well as several genes directly associated with AD including *APOE*, a major genetic risk factor locus in humans. For ancestral Cebidae, we recovered the important related gene *ADAM10*, an alpha secretase involved in the cleavage of APP thereby preventing the generation of amyloid beta peptides associated with the development of AD (*54*).

Notably, there is a strong relationship between circadian rhythms and aging. Emerging in early infancy, the circadian system undergoes significant changes through an organism’s lifespan affecting rhythms of behaviours, temperature regulation, and hormone release, among others, and is implicated in human longevity (*55*). We found signatures of selection on circadian rhythms in both ancestral Cebinae and across-capuchins represented by enriched BP GO terms and UP keywords and including genes encoding core components of the circadian clock such as *PER3*. Precisely timed rhythmic activities that are tuned to periodic biotic and abiotic cycles of an organism’s environment are likely to confer adaptive advantage (*56*) and these signatures may relate to a variety of factors including, for example, the high activity levels of capuchins, as well as capuchin longevity, slow maturation, and/or delayed life history.

### Behaviour and cognition

The behavioural diversity characterising capuchin monkeys includes social conventions and local traditions, complex and intimate social relationships, ecological and dietary flexibility, tool use, and extractive foraging including an astounding degree of planning, with capuchin behaviour varying by age, sex, and geographically across populations of the same species (*57*–*60*). Crucial neuromodulators influencing the brain and likely shaping this behavioural variation include the major neurotransmitter systems, as well as neuropeptides and hormones. Neuromodulators play a prominent role in nervous system function, from simple reflexes to influencing synaptic plasticity and neurogenesis, and mediating higher cognitive processes such as sensory processing, memory encoding, learning, mood, and decision-making, with essential roles in modulating behaviour (*61, 62*).

We recover broad and sweeping signatures of selection on hormones, neuropeptides, and behaviour for the cebine branches, particularly for ancestral Cebinae and across-capuchins, with many related enriched terms. Interesting genes found across these annotations include those encoding neuropeptides and receptors that play important roles in many physiologic processes including cognition, memory, sensory/pain processing, stress, hormone and insulin secretion, appetite regulation, metabolism, and energy homeostasis. Several of these genes encode peptides that play central roles in feeding behaviour including ghrelin and obestatin (*GHRL*), and leptin (*LEP*), as well as other important neuropeptide related genes (e.g., *NPFF, NPFFR1, GALR3, BSX, GPR39*). Signatures of selection related to other crucial neuromodulators include on three genes encoding serotonin receptors for the various capuchin branches, and on at least three genes involved in dopamine signalling for ancestral Cebinae. The ancestral Cebinae gene set is also enriched for terms related to agmatine biosynthesis (including the gene *AGMAT*); agmatine is widely and unevenly distributed in the mammalian brain, acting as a neuromodulator that may directly participate in learning and memory processes, and is sometimes taken orally to treat depression in humans (*63*).

There are multiple genes in the hormone annotations in the ancestral Cebinae and/or across-capuchin gene sets related to the thyroid hormone (TH) and thyroid-stimulating hormone (e.g., *TRH, CGA, PAX8*). TH is a key metabolic hormone with many physiologic functions including critical roles in differentiation, growth, and metabolism. TH dramatically impacts mammalian brain development, with its importance highlighted by the deleterious and irreversible effects of TH deficiency/dysfunction during foetal and neonatal periods. TH also plays important roles in normal adult brain function and has a profound influence on behaviour throughout life, with adult-onset TH dysfunction associated with a range of CNS-related pathologies, neurological and behavioural abnormalities, and alterations in mood and cognition (*64*).

While capuchins are known for their cognitive and social behaviours, squirrel monkeys live in extremely large groups and also show differentiated social relationships and prosocial behaviours that they share with capuchins such as predator mobbing and alarm calls (*65, 66*). For ancestral Cebidae, signatures of selection related to neuromodulation are recovered on genes such as *DBH*, dopamine beta-hydroxylase, which catalyses the conversion of dopamine to noradrenaline, playing a role in the bioavailability of both crucial neuromodulators, as well as another serotonin receptor, several neuropeptide receptors, and *RPH3A. RPH3A* plays an important role in neurotransmitter release and is involved in the exocytosis of arginine vasopressin (AVP), which is notable given AVP mediates complex mammalian social behaviours. For squirrel monkeys, we recover the enriched BP GO terms “dopamine metabolic process” and “dopamine biosynthetic process”.

### Reproduction and mating systems

Platyrrhine primates show a diverse range of mating systems and sexual/reproductive characteristics and behaviours. Both capuchin and squirrel monkeys are characterised by multi-male multi-female mating systems (polygynandry), unlike many of their closely related lineages, for example, the flexible polyandry-monogamy seen in callitrichids and the social monogamy of owl monkeys. Polygynandrous mating systems are associated with post-copulatory sexual selection through sperm competition (*67*). Sperm competition can be directed at the quantity and quality of sperm, for example, effectuated via changes to rates of spermatogenesis, sperm cell size, morphology, and mobility, copulation frequency, testes size, and the morphology of the penis, accessory glands, and ducts (*67*). In line with the reproductive shift to polygynandry in ancestral Cebidae, we find strong and sweeping signatures of selection putatively related to sperm competition in both gene sets with various enriched terms describing motile cilium and flagella, spermatogenesis, sperm development, male meiosis, fertilisation, and reproduction. These terms cover a suite of interesting genes including two important members of the CatSper complex, a sperm-specific ion channel involved in several important steps of fertilisation including sperm hyperactivation and capacitation, which allow sperm to reach and interact with an oocyte. Similar signatures can also be found for ancestral Cebinae with “fertilisation” forming the top ranked individual BP GO term, as well as more broadly on motile cilia. In line with the shift to multi-male multi-female mating systems, cebid lineages are also characterised by larger group sizes, which is particularly notable for squirrel monkeys (*68*), and may underlie some of the enriched immune system related results recovered for ancestral Cebidae, squirrel monkeys, and ancestral Cebinae.

The behavioural repertoire of capuchins includes new reproductive/courtship behaviours and complex intimate individual relationships, which may also relate to their mating system (*69*). In agreement, we recovered genes related to sex steroids and reproductive hormones/peptides in the ancestral Cebinae and across-capuchins gene sets, many of which are contained in aforementioned enriched hormone-related terms discussed in the previous section. For across-capuchins, we also recovered more specific enriched GO terms describing the secretion of gonadotropin, luteinising hormone, and endocrine hormones. Several genes in these terms are involved in sex steroid metabolism, while others are associated with the pituitary glycoprotein hormones and prolactin such as *PRLH*, prolactin releasing hormone, which stimulates prolactin release and regulates prolactin expression, as well as lactation, behaviour, and the reproductive system. Among the most notable genes recovered for capuchins is *NPVF*, found in both the ancestral Cebinae and across-capuchins branches, encoding the neuropeptides NPSF and NPVF (also referred to as the RFamide-related peptides, RFRP-1 and RFRP-3), which are mammalian homologs of the avian neuropeptide gonadotropin-inhibitory hormone. These neuropeptides act as potent negative regulators of gonadotropin synthesis and secretion, with a range of functions in the modulation of reproduction including the regulation of sexual behaviour, sexual maturation, ovulatory cycle, gonadal function, reproductive seasonality, and stress-induced reproductive suppression, among others (*70*).

### Body size and morphology

While the two capuchin lineages, robust and gracile, share many traits as discussed throughout, significant derived characters arose since their divergence 5 to 6 mya, hence their division into two genera, *Sapajus* and *Cebus* (*59, 71*). The most notable differences relate to their body shape and skeletal morphology, and this is reflected in the strongest signatures of selection recovered individually for both lineages. In line with their name, robust capuchins (*Sapajus*) are generally stockier and more skeletally robust, with shorter, thicker limbs, as well as striking differences in cranio-dental morphology particularly relating to the robust masticatory architecture of the skull, a specialisation to process tougher foods (durophagy) such as encased nuts and palm fruits (*59, 72*). Further, *Sapajus libidinosus* is known to habitually use stone tools to access a variety of encased foods including otherwise inaccessible foods, a skill that takes many years to perfect, and this ability likely relates to their more robust skeletal morphology (*71, 73*). This derived morphology is reflected in the range of enriched GO terms and candidate genes we found in this lineage related to facial, skeletal system, and skeletal muscle tissue morphogenesis and development, as well as BMP signalling pathway. Several genes in these terms are explicitly associated with skull bone fusion and morphology such as *SIX4*, which plays an important role in cranial morphogenesis and synchondrosis formation during embryonic development (*74*). Another is *RAB23*, which functions in limb patterning, coordinating early osteogenesis, and controlling the growth and fusion of developing skull bones, and is implicated in the premature fusion of craniofacial sutures seen in Carpenter syndrome (*75*). A third interesting gene in these annotations encodes the protein delangin (*NIPBL*) that plays a role in the development of the limbs and skull/face bones; defects in this gene are the primary cause of Cornelia de Lange syndrome, which is characterised by distinctive facial features, limb/skeletal dysmorphology, and slow postnatal growth (*76*).

Similarly, for gracile capuchins (*Cebus*)—characterised by long slender limbs and a slighter body plan (*59, 77*)—we recovered various enriched annotated terms related to limb and skeletal system development, including several homeobox transcription factors of the Hox (5 of 21 analysed) and Shox (1 of 2 analysed) families that play fundamental roles in embryonic pattern formation, axis control, and are required for normal limb development (*78*). Many other genes in these terms are also associated with various skeletal dysmorphologies and congenital limb defects in humans. More broadly, there are signatures of selection on embryonic development for gracile capuchins including enriched terms such as the GO term “chordate embryonic development” and UP keyword “developmental protein”. Taken together, the results for robust and gracile capuchins are suggestive of adaptive pressure on developmental pathways related to the skull/face, limbs, and skeletal system that may underlie the morphological differences between these capuchin lineages.

Also related to morphology, we recover enriched the GO term “embryonic digit morphogenesis” for ancestral Cebinae. Capuchins have a high degree of manual dexterity, possessing pseudo-opposable thumbs augmenting their precision grip ability, which plays a role in their sensorimotor intelligence, and show increased dexterity compared to squirrel monkeys (*79, 80*).

Among the most unusual aspects of squirrel monkey biology is their large brain size in the context of their overall small body size, which distinguishes them from the other most encephalised primate lineages. Reconstructions have indicated that squirrel monkey body size decreased and their brain size increased further after squirrel monkeys and capuchins diverged (*7*). Our results for squirrel monkeys reveal broad signatures related to growth factors across our analyses with enriched growth factor related GO terms and genes encoding or associated with members of the fibroblast growth factor and transforming growth factor beta families, and many implicated in human stature and dwarfism including the short stature homeobox gene (*SHOX*). The most significant selective signatures for squirrel monkeys relate to cellular signalling cascades with various enriched annotations describing the mitogen-activated protein kinase (MAPK) and ERK1/2 signalling pathways involved in basic cellular processes including cell proliferation and differentiation. Together, the signatures of selection on ERK/MAPK cascades and growth factors may be related to the reduced body size of squirrel monkeys, and/or the adaptive maintenance of a large brain size while reducing body size.

### Diet and nutrition

Capuchins inhabit a complex omnivorous dietary niche characterised by dietary flexibility, high nutrient density, and easy digestibility for their small gut (*81*), with high sensorimotor intelligence related to their extractive foraging capabilities. For across-capuchins, we recovered various diet/metabolism related signatures including for branched chain amino acids (BCAAs), essential amino acids required in the diet that are major constituents of muscle protein; riboflavin, a B vitamin involved in many physiologic processes, necessary for normal cell growth and function; and biotin, another essential B vitamin involved in the conversion of food to energy, and important for embryonic growth. It is notable that all of these nutrients are found in lipid- and protein-rich food sources such as meats, eggs, and nuts. Among the capuchin lineages, robust capuchins show a preference for food with a high lipid content such as nuts and insects (*82*), and we recover various enriched GO terms related to lipid metabolism, which may be linked to their increased ability (versus gracile capuchins) to access fat-rich nuts as a result of both their robust skeletal morphology and, in some species, their stone tool use. Robust capuchins also show various signatures potentially related to water homeostasis including enriched GO terms for kidney/renal system development and sodium ion transport. Selective pressure on water homeostasis may relate to range expansion into drier habitats such as the Cerrado for some *Sapajus* lineages in the Pleistocene (*83*).

Similarly, in the highly insectivorous squirrel monkeys (*9*), we recovered various enriched terms related to nutrition including mineral absorption, response to metal ions, retinoid metabolism, and calcium homeostasis, which is notable given many insects are considered a poor source of calcium.

### Limitations and future directions

While a single genome per species or lineage can give insight into evolutionary processes deep in time, the sequencing of more individuals in each of these lineages will be critical for studying patterns of demography and selection in more recent history. The inclusion of additional individuals is also required to determine if variants discovered in this study are fixed or vary within species. Moreover, without functional genomic experiments, some of the significant genes described in this study might reflect the relaxation of selection rather than adaptive evolution, and these genes thus remain candidates until they are validated. Furthermore, given protein function, which is generally derived from humans, mice, and other model organisms, is little understood in the context of the biology of these cebid lineages, the functional significance of selection on these candidate genes and the association of these signals with specific adaptive functions is correlative. Codon-based models of evolution are also unable to consider variation in regulatory controls and gene expression, which can both also have important adaptive implications.

The new draft reference assembly, short read data, and RNAseq data from 17 tissues for the same robust capuchin individual provide a useful resource for future genomics studies of capuchins and primates more broadly. Future directions might include long read sequencing, a candidate technology to fill gaps in the assembly and increase the contiguity to chromosome scale. The results from this study will be useful in downstream applications for the study of genes of interest in both captive and field studies of platyrrhines, as well as opening new avenues of research for the study of primate brain evolution and comparative brain biology.

## Materials and Methods

### Genome sequencing, assembly & size estimation

Whole blood was collected during a routine physical from Mango, a female captive brown robust capuchin (*Sapajus apella*) housed at the Language Research Center, Georgia State University (IACUC number: A16031). Mango was aged and thought to have been wild-caught in the 1970s, she was the last remaining individual from the colony’s original source population. Dovetail Genomics extracted high molecular weight DNA from the blood sample to construct one shotgun library and three “Chicago” proximity ligation libraries with chimeric pairs spanning up to 50 Kbp in physical distance. The shotgun library was sequenced across four HiSeq 4000 lanes producing 1.33 billion 150 bp paired end (PE) read pairs (399 Gbp), an estimated 148-fold sequencing coverage (based on a genome size of 2.7 Gbp). The three Chicago libraries were pooled and sequenced across two HiSeq 4000 lanes generating 800 million 100 bp PE read pairs with ∼220-fold physical coverage. All sequencing was performed at the DNA Technologies Core, UC Davis. All raw reads were deposited on NCBI’s sequence read archive (SRA) (Table S1). A preliminary *de novo* assembly was generated by Dovetail Genomics from quality-filtered short read shotgun data using the Meraculous assembler (*84*). The final draft assembly was generated by scaffolding the preliminary assembly with the Chicago libraries using Dovetail’s HiRise pipeline (*13*). Total length of this genome assembly was 2,520.3 Mbp (in 6631 scaffolds) with an N50 of 27.1 Mbp (29 scaffolds) and N90 of 4.04 Mbp (116 scaffolds). The longest scaffold was 90.4 Mbp.

We evaluated completeness of the genome assembly by its estimated gene content using CEGMA v2.5 (*15*) and BUSCO v3.0.2 (*14*) to calculate the proportion of 248 CEGs or 6,192 Euarchontoglires-specific conserved single copy orthologs, respectively, that were either complete, fragmented, or missing. Using quality-filtered, nuclear only, endogenous short reads, we also performed *k*-mer counting with Jellyfish v.2.2.6 (*85*) to generate a *k*-mer frequency distribution of 31-mers and then estimated genome size using four approaches. We generated an initial mitochondrial genome assembly for Mango by mapping a set of putative mitochondrial short read pairs to a complete *S. apella* mitochondrial genome using MIRA v.4.0.2 (*86*), and then performing baiting and iterative mapping with a MITObim v.1.9.1 (*87*) wrapper script to generate the final mitochondrial genome assembly.

### RNA sequencing & transcript assemblies

The reference individual, Mango, was euthanised in the months after genome sequencing when a cancerous tumour was discovered, allowing the ethical collection of fresh tissue for RNA sequencing from the same individual. Tissue collection was performed during necropsy at Yerkes National Primate Research Center within hours of her death. Seventeen tissues samples were harvested and placed in RNAlater (Invitrogen), and subsequently, total RNA was isolated from each sample followed by poly-A tail selection library preparation. The libraries were pooled and sequenced on a single HiSeq 3000 lane generating ∼367 million 150 bp PE read pairs (102.5 Gbp) with between 16.8 and 27.4 million reads pairs per tissue. These steps were performed by the Technology Center for Genomics & Bioinformatics (TCGB) at UCLA. All raw reads were deposited on NCBI’s SRA (Table S1). After *k*-mer correction, filtering, trimming, and rRNA removal steps, we generated seven transcript assemblies with the cleaned RNAseq read pairs, as follows: *de novo* (TrinDNv2); abundance filtered *de novo* (TrinDNv2); reference-based (Cuffv1); PASA with TrinDNv2 and Cuffv1 as input (PASAv1); genome-guided (TrinGGv1); PASA with TrinDNv2, Cuffv1, and TrinGGv1 as input (PASAv2); and non-redundant with PASAv2 as input (NRv1). This ultimately resulted in three assemblies (TrinDNv2, PASAv1, and NRv1) for use as direct evidence in various iterations of the genome annotation pipeline. We checked quality metrics and completeness of the seven assemblies using rnaQUAST v1.5.0 (*17*) with BUSCO v3.0.2 in transcriptome mode using the Euarchontoglires-specific BUSCOs gene set.

### Repeat & genome annotation

To assess the repeat content of the robust capuchin genome, we first performed a homology-based repeat annotation of our genome assembly using known elements with RepeatMasker v4.0.7 (*88*), followed by *de novo* repeat identification using the library of unknown repeats generated with RepeatModeler v1.0.11 (*89*), and finally, we used ProcessRepeats from RepeatMasker to summarise all annotated repeats in the genome assembly. We annotated the robust capuchin genome assembly in three iterations of Maker v3.01.02 (*18, 19*) to predict gene models, incorporating direct evidence from transcript assemblies, homology to the predicted proteomes of platyrrhine primates and humans, and *ab initio* predictions from Augustus v3.3 (*90*) with a robust capuchin-specific HMM that was trained initially in BUSCO and twice subsequently using high-quality gene models from each of the first two passes of Maker (Table S6). Predicted gene models from the third pass of Maker were functionally annotated using Blast2GO v5.2.5 (*91*) and filtered based on supporting evidence and presence of annotations.

### Identification of orthologs, alignment & filtering

In order to assess signatures of positive selection in other platyrrhine primate genomes, we first identified orthologs using the OrthoMCL pipeline (*92*) across ten species; four platyrrhine primates (*Sapajus*; *Cebus*; *Saimiri*; *Callithrix*), five other primates (*Macaca, Pan, Homo, Carlito, Microcebus*), and mouse (*Mus*). As input to the pipeline, we used predicted CDS and protein sequence files from Ensembl (or for *Sapajus* from our genome annotation) for all species that were filtered for the longest isoform per gene. We generated a set of 9,216 conservative, manually-curated CDS alignments which were highly likely to represent one-to-one orthologs across their length by: (i) filtering the OrthoMCL output for one-to-one orthologs, a minimum of five species, and the presence of at least one capuchin lineage; (ii) aligning CDS sequences for these filtered orthologs groups by codon using Guidance2 v.2.02 (*20*) with the MAFFT aligner v.7.419 (*93*) with 100 guidance bootstraps; and (iii) visually inspecting all alignments for errors and editing as required to reduce the likelihood of false positives.

### Branch model & branch-site model tests

We specified six lineages (foreground branches) for the positive selection tests, as follows: (H1) robust capuchin (*Sapajus*); (H2) gracile capuchin (*Cebus*); (H3) ancestral Cebinae (capuchins); (H3a) across-capuchins more generally (all Cebinae); (H4) ancestral Cebidae; and (H5) squirrel monkey (*Saimiri*). We assigned species set IDs to each combination of species (207 species sets) found in the final alignments and generated unrooted tree files that specified the various foreground branches analysed for each species set (759 tree files). We ran two different tests for positive selection with codeml from the PAML package v.4.9 (*21*) which are based on rates of non-synonymous versus synonymous substitutions (ω or dN/dS ratio): the branch-site model (BSM), which tests for episodic selection by searching for positively selected sites in the foreground branch; and (B) the branch model (BM), which tests for elevated dN/dS ratios along the foreground branch indicating accelerated evolution. We did not run the BSM test for H3a, thus a total of 11 lineage and test combinations were conducted with codeml. For each BM test, we assessed two models as follows; the alternative branch model which separates the tree into foreground and background branches that have distinct ω parameters allowing them to evolve with separate dN/dS ratios, and the null model which uses a single ω parameter across the tree. For each BSM test, we assessed an alternative branch-site model allowing for positive selection on the foreground branch and a null model allowing only for purifying and neutral selection on the foreground and background lineages. After estimating parameters and calculating the likelihood with codeml, we performed likelihood ratio tests (LRTs) by comparing the likelihood of the alignment under the alternative versus under the null model, and calculated p-values from the chi-square distribution with one degree of freedom.

We conducted 11 gene set enrichment analyses, one for the set of significant genes from each combination of lineage and test (BM or BSM) using DAVID v.6.8 (*22*) with the entire human gene set as the background population of genes. In DAVID, we assessed lists/charts of enriched (i) BP, CC, and MF GO terms (the “all” option), (ii) UP keywords, (iii) KEGG pathways, (iv) Reactome pathways, and (v) disease annotations, as well as functional annotation clustering across the three GO terms together under the high classification stringency criteria, with an EASE score of < 0.05 required for all enriched annotated terms for both approaches.

## Acknowledgments

We thank Jonathon Rodgers and LSSA Support at UCLA for computational assistance; staff at Dovetail Genomics, DNA Technologies Core (UC Davis), and TCGB (UCLA) for sequencing assistance; Amelia Wilkes and the staff at the Language Research Center for Mango’s care and the collection of her blood sample; staff at Yerkes National Primate Research Center for tissue collection for RNAseq during Mango’s necropsy; the Broad Institute, Amanda Melin, and Joe Orkin for generating the squirrel monkey and gracile capuchin genomes used in this study; Stephen Nash for the use of his illustrations; Colin Brand for his helpful comments; and the Institute for Society and Genetics (UCLA) and Anthropology Dept. (University of Utah) for postdoctoral support for HB.

## Author contributions

Conceptualisation: HB, JWL, PI

Sample & data collection: HB, JWL, SFB

Reference genome assembly analyses: HB

Positive selection & enrichment analyses: HB

Interpretation: HB, JWL, THW, PI

Supervision: JWL, THW

Writing—original draft: HB

Writing—review & editing: HB, THW, JWL, PI, SFB

## Data and materials availability

The reference genome, WGS and Chicago library sequencing reads, and RNAseq reads for 17 tissues for our reference *Sapajus apella* individual are available at NCBI BioProjects under the accession no. PRJNA717806 (www.ncbi.nlm.nih.gov/bioproject/717806) [**to be released upon acceptance of the manuscript for publication**]. The version of the reference genome assembly used in this study, as well as the mitochondrial genome assembly and annotation, are available on a Zenodo repository (https://doi.org/10.5281/zenodo.5225106).

## Supplementary Materials

One supplementary document with extended methods and results, supplementary figures S1, S2 and S3, and supplementary tables S1, S2, S3, S5, S6, S7, S10, and S23. Supplementary tables S4, S8, S9, S11 to S22, and S24 to S44 are found as separate excel files.

## References

1. P. Perelman, W. E. Johnson, C. Roos, H. N. Seuánez, J. E. Horvath, M. a M. Moreira, B. Kessing, J. Pontius, M. E. Roelke, Y. Rumpler, M. P. C. Schneider, A. Silva, S. J. O’Brien, J. Pecon-Slattery, A molecular phylogeny of living primates. PLoS Genet. 7, e1001342 (2011).

2. M. Bond, M. F. Tejedor, K. E. Campbell, L. Chornogubsky, N. Novo, F. Goin, Eocene primates of South America and the African origins of New World monkeys. Nature. 520, 538–541 (2015).

3. J. Cracraft, C. C. Ribas, F. M. D’Horta, J. Bates, R. P. Almeida, A. Aleixo, J. P. Boubli, K. E. Campbell, F. W. Cruz, M. Ferreira, S. C. Fritz, C. H. Grohmann, E. M. Latrubesse, L. G. Lohmann, L. J. Musher, A. Nogueira, A. O. Sawakuchi, P. Baker, “The origin and evolution of Amazonian species diversity” in Neotropical Diversification: Patterns and Processes, V. Rull, A. Carnaval, Eds. (Springer, Heidelberg, 2020), pp. 225–244.

4. N. M. Jameson Kiesling, S. V Yi, K. Xu, F. Gianluca Sperone, D. E. Wildman, The tempo and mode of New World monkey evolution and biogeography in the context of phylogenomic analysis. Mol. Phylogenet. Evol. 82 Pt B, 386–99 (2015).

5. A. Estrada, P. A. Garber, A. B. Rylands, C. Roos, E. Fernandez-Duque, A. Di Fiore, K. Anne-Isola Nekaris, V. Nijman, E. W. Heymann, J. E. Lambert, F. Rovero, C. Barelli, J. M. Setchell, T. R. Gillespie, R. A. Mittermeier, L. V. Arregoitia, M. de Guinea, S. Gouveia, R. Dobrovolski, S. Shanee, N. Shanee, S. A. Boyle, A. Fuentes, K. C. MacKinnon, K. R. Amato, A. L. S. Meyer, S. Wich, R. W. Sussman, R. Pan, I. Kone, B. Li, Impending extinction crisis of the world’s primates: Why primates matter. Sci. Adv. 3 (2017), doi:10.1126/sciadv.1600946.

6. A. Rylands, R. A. Mittermeier, “Family Cebidae (capuchins and squirrel monkeys)” in Handbook of the mammals of the world. Volume 3, R. Mittermeier, A. B. Rylands, D. E. Wilson, Eds. (Lynx Edicions, Barcelona, 2013), pp. 348–389.

7. S. H. Montgomery, I. Capellini, R. A. Barton, N. I. Mundy, Reconstructing the ups and downs of primate brain evolution: implications for adaptive hypotheses and Homo floresiensis. BMC Biol. 8, 1–19 (2010).

8. D. M. Fragaszy, E. Visalberghi, L. M. Fedigan, Eds., The complete capuchin: the biology of the genus Cebus (Cambridge University Press, Cambridge, 2004).

9. H. S. Zimbler-Delorenzo, A. I. Stone, Integration of field and captive studies for understanding the behavioral ecology of the squirrel monkey (Saimiri sp.). Am. J. Primatol. 73, 607–622 (2011).

10. S. D. Tardif, C. R. Abee, K. G. Mansfield, Workshop summary: Neotropical primates in biomedical research. ILAR J. 52, 386–392 (2011).

11. J. D. Orkin, M. J. Montague, D. Tejada-Martinez, M. de Manuel, J. del Campo, S. C. Hernandez, A. Di Fiore, C. Fontsere, J. A. Hodgson, M. C. Janiak, L. F. K. Kuderna, E. Lizano, M. P. Martin, Y. Niimura, G. H. Perry, C. S. Valverde, J. Tang, W. C. Warren, J. P. de Magalhães, S. Kawamura, T. Marquès-Bonet, R. Krawetz, A. D. Melin, The genomics of ecological flexibility, large brains, and long lives in capuchin monkeys revealed with fecalFACS. Proc. Natl. Acad. Sci. U. S. A. 118 (2021), doi:10.1073/pnas.2010632118.

12. A. M. Boddy, P. W. Harrison, S. H. Montgomery, J. A. Caravas, M. A. Raghanti, K. A. Phillips, N. I. Mundy, D. E. Wildman, Evidence of a conserved molecular response to selection for increased brain size in primates. Genome Biol. Evol. 9, 700–713 (2017).

13. N. H. Putnam, B. O. Connell, J. C. Stites, B. J. Rice, P. D. Hartley, C. W. Sugnet, D. Haussler, D. S. Rokhsar, Chromosome-scale shotgun assembly using an in vitro method for long-range linkage. Genome Res. 26, 342–350 (2016).

14. F. A. Simão, R. M. Waterhouse, P. Ioannidis, E. V. Kriventseva, E. M. Zdobnov, BUSCO: Assessing genome assembly and annotation completeness with single-copy orthologs. Bioinformatics. 31, 3210–3212 (2015).

15. G. Parra, K. Bradnam, I. Korf, CEGMA: A pipeline to accurately annotate core genes in eukaryotic genomes. Bioinformatics. 23, 1061–1067 (2007).

16. L. Fantini, M. D. Mudry, M. Nieves, Genome size of two Cebus species (Primates: Platyrrhini) with a fertile hybrid and their quantitative genomic differences. Cytogenet. Genome Res. 135, 33–41 (2011).

17. E. Bushmanova, D. Antipov, A. Lapidus, V. Suvorov, A. D. Prjibelski, RnaQUAST: A quality assessment tool for de novo transcriptome assemblies. Bioinformatics. 32, 2210–2212 (2016).

18. C. Holt, M. Yandell, MAKER2: An annotation pipeline and genome-database management tool for second-generation genome projects. BMC Bioinformatics. 12 (2011), doi:10.1186/1471-2105-12-491.

19. M. S. Campbell, M. Y. Law, C. Holt, J. C. Stein, G. D. Moghe, D. E. Hufnagel, J. Lei, R. Achawanantakun, D. Jiao, C. J. Lawrence, D. Ware, S. H. Shiu, K. L. Childs, Y. Sun, N. Jiang, M. Yandell, MAKER-P: A Tool kit for the rapid creation, management, and quality control of plant genome annotations. Plant Physiol. 164, 513–524 (2014).

20. I. Sela, H. Ashkenazy, K. Katoh, T. Pupko, GUIDANCE2: Accurate detection of unreliable alignment regions accounting for the uncertainty of multiple parameters. Nucleic Acids Res. 43, W7–W14 (2015).

21. Z. Yang, PAML 4: Phylogenetic analysis by maximum likelihood. Mol. Biol. Evol. 24, 1586–1591 (2007).

22. D. W. Huang, B. T. Sherman, R. A. Lempicki, Systematic and integrative analysis of large gene lists using DAVID bioinformatics resources. Nat. Protoc. 4, 44–57 (2009).

23. H. J. Jerison, Animal intelligence as encephalization. Philos. Trans. R. Soc. Lond. B. Biol. Sci. 308, 21–35 (1985).

24. A. M. Boddy, M. R. Mcgowen, C. C. Sherwood, L. I. Grossman, M. Goodman, D. E. Wildman, Comparative analysis of encephalization in mammals reveals relaxed constraints on anthropoid primate and cetacean brain scaling. J. Evol. Biol. 25, 981–994 (2012).

25. K. L. Chiou, L. Pozzi, J. W. Lynch Alfaro, A. Di Fiore, Pleistocene diversification of living squirrel monkeys (Saimiri spp.) inferred from complete mitochondrial genome sequences. Mol. Phylogenet. Evol. 59, 736–745 (2011).

26. M. Igarashi, Molecular basis of the functions of the mammalian neuronal growth cone revealed using new methods. Proc. Japan Acad. Ser. B Phys. Biol. Sci. 95, 358–377 (2019).

27. J. Guo, J. M. Otis, S. K. Suciu, C. Catalano, L. Xing, S. Constable, D. Wachten, S. Gupton, J. Lee, A. Lee, K. H. Blackley, T. Ptacek, J. M. Simon, S. Schurmans, G. D. Stuber, T. Caspary, E. S. Anton, Primary cilia signaling promotes axonal tract development and is disrupted in Joubert syndrome-related disorders models. Dev. Cell. 51, 759-774.e5 (2019).

28. S. M. Park, H. J. Jang, J. H. Lee, Roles of primary cilia in the developing brain. Front. Cell. Neurosci. 13, 1–10 (2019).

29. Y. Q. Wang, Y. P. Qian, S. Yang, H. Shi, C. H. Liao, H. K. Zheng, J. Wang, A. A. Lin, L. L. Cavalli-Sforza, P. A. Underhill, R. Chakraborty, L. Jin, B. Su, Accelerated evolution of the pituitary adenylate cyclase-activating polypeptide precursor gene during human origin. Genetics. 170, 801–806 (2005).

30. H. Stephan, G. Baron, H. D. Frahm, Comparative size of brains and brain structures. Comp. primate Biol. 4, 1–38 (1988).

31. J. D. Sweatt, Neural plasticity and behavior – sixty years of conceptual advances. J. Neurochem. 139, 179–199 (2016).

32. J. W. Fawcett, T. Oohashi, T. Pizzorusso, The roles of perineuronal nets and the perinodal extracellular matrix in neuronal function. Nat. Rev. Neurosci. 20, 451–465 (2019).

33. A. C. Reichelt, D. J. Hare, T. J. Bussey, L. M. Saksida, Perineuronal Nets: Plasticity, Protection, and Therapeutic Potential. Trends Neurosci. 42, 458–470 (2019).

34. M. Shimojo, J. Madara, S. Pankow, X. Liu, J. Yates, T. C. Südhof, A. Maximov, Synaptotagmin-11 mediates a vesicle trafficking pathway that is essential for development and synaptic plasticity. Genes Dev. 33, 365–376 (2019).

35. M. A. Lancaster, D. J. Gopal, J. Kim, S. N. Saleem, J. L. Silhavy, C. M. Louie, B. E. Thacker, Y. Williams, M. S. Zaki, J. G. Gleeson, Defective Wnt-dependent cerebellar midline fusion in a mouse model of Joubert syndrome. Nat. Med. 17, 726–731 (2011).

36. R. J. Ferland, W. Eyaid, R. V. Collura, L. D. Tully, R. S. Hill, D. Al-Nouri, A. Al-Rumayyan, M. Topcu, G. Gascon, A. Bodell, Y. Y. Shugart, M. Ruvolo, C. A. Walsh, Abnormal cerebellar development and axonal decussation due to mutations in AHI1 in Joubert syndrome. Nat. Genet. 36, 1008–1013 (2004).

37. J. W. Mink, R. J. Blumenschine, D. B. Adams, Ratio of central nervous system to body metabolism in vertebrates: Its constancy and functional basis. Am. J. Physiol. - Regul. Integr. Comp. Physiol. 10, 203–212 (1981).

38. M. Rango, N. Bresolin, Brain mitochondria, aging, and Parkinson’s disease. Genes (Basel). 9 (2018), doi:10.3390/genes9050250.

39. J. W. Doan, T. R. Schmidt, D. E. Wildman, M. Uddin, A. Goldberg, M. Hüttemann, M. Goodman, M. L. Weiss, L. I. Grossman, Coadaptive evolution in cytochrome c oxidase: 9 of 13 subunits show accelerated rates of nonsynonymous substitution in anthropoid primates. Mol. Phylogenet. Evol. 33, 944–950 (2004).

40. L. I. Grossman, D. E. Wildman, T. R. Schmidt, M. Goodman, Accelerated evolution of the electron transport chain in anthropoid primates. Trends Genet. 20, 578–585 (2004).

41. M. Goodman, K. N. Sterner, M. Islam, M. Uddin, C. C. Sherwood, P. R. Hof, Z. C. Hou, L. Lipovich, H. Jia, L. I. Grossman, D. E. Wildman, Phylogenomic analyses reveal convergent patterns of adaptive evolution in elephant and human ancestries. Proc. Natl. Acad. Sci. U. S. A. 106, 20824–20829 (2009).

42. Y. Y. Shen, L. Liang, Z. H. Zhu, W. P. Zhou, D. M. Irwin, Y. P. Zhang, Adaptive evolution of energy metabolism genes and the origin of flight in bats. Proc. Natl. Acad. Sci. U. S. A. 107, 8666–8671 (2010).

43. A. Cheng, Y. Hou, M. P. Mattson, Mitochondria and neuroplasticity. ASN Neuro. 2, 243–256 (2010).

44. N. Sun, R. J. Youle, T. Finkel, The mitochondrial basis of aging. Mol. Cell. 61, 654–666 (2016).

45. J. M. Allman, T. McLaughlin, A. Hakeem, Brain structures and life-span in primate species. Proc. Natl. Acad. Sci. U. S. A. 90, 3559–3563 (1993).

46. C. González-Lagos, D. Sol, S. M. Reader, Large-brained mammals live longer. J. Evol. Biol. 23, 1064–1074 (2010).

47. C. P. van Schaik, K. Isler, “Life history evolution in primates” in The evolution of primate societies, J. C. Mitani, J. Call, P. M. Kappeler, R. A. Palombit, J. B. Silk, Eds. (University of Chicago Press, Chicago, 2012), pp. 220–244.

48. A. M. Bronikowski, M. Cords, S. C. Alberts, J. Altmann, D. K. Brockman, L. M. Fedigan, Pusey, T. Stoinski, K. B. Strier, W. F. Morris, Female and male life tables for seven wild primate species. Sci. Data. 3, 1–8 (2016).

49. M. Yousefzadeh, C. Henpita, R. Vyas, C. Soto-Palma, P. Robbins, L. Niedernhofer, Dna damage—how and why we age? Elife. 10, 1–17 (2021).

50. A. S. Multani, S. Chang, WRN at telomeres: Implications for aging and cancer. J. Cell Sci. 120, 713–721 (2007).

51. M. H. Park, H. K. Jin, J. Bae, Potential therapeutic target for aging and age-related neurodegenerative diseases: the role of acid sphingomyelinase. Exp. Mol. Med. 52, 380–389 (2020).

52. S. M. Jazwinski, S. Kim, J. Dai, L. Li, X. Bi, J. C. Jiang, J. Arnold, M. A. Batzer, J. A. Walker, D. A. Welsh, C. M. Lefante, J. Volaufova, L. Myers, L. J. Su, D. B. Hausman, M. V. Miceli, E. Ravussin, L. W. Poon, K. E. Cherry, M. A. Welsch, HRAS1 and LASS1 with APOE are associated with human longevity and healthy aging. Aging Cell. 9, 698–708 (2010).

53. A. Von Gunten, M. therese Clerc, R. Tomar, P. S. John Smith, Evolutionary considerations on aging and Alzheimer’s disease. J. Alzheimer’s Dis. Park. 08, 1–12 (2018).

54. S. Lammich, E. Kojro, R. Postina, S. Gilbert, R. Pfeiffer, M. Jasionowski, C. Haass, F. Fahrenholz, Constitutive and regulated α-secretase cleavage of Alzheimer’s amyloid precursor protein by a disintegrin metalloprotease. Proc. Natl. Acad. Sci. U. S. A. 96, 3922–3927 (1999).

55. O. Froy, Circadian rhythms, aging, and life span in mammals. Physiology. 26, 225–235 (2011).

56. D. A. Paranjpe, V. K. Sharma, Evolution of temporal order in living organisms. J. Circadian Rhythms. 3, 1–13 (2005).

57. E. B. Ottoni, P. Izar, Capuchin monkey tool use: Overview and implications. Evol. Anthropol. 17, 171–178 (2008).

58. S. Perry, Social traditions and social learning in capuchin monkeys (Cebus). Philos. Trans. R. Soc. B Biol. Sci. 366, 988–996 (2011).

59. J. W. Lynch Alfaro, J. de S. E. Silva-Jr, A. B. Rylands, How different are robust and gracile capuchin monkeys? An argument for the use of Sapajus and Cebus. Am. J. Primatol. 74, 273–286 (2012).

60. S. Perry, Behavioural variation and learning across the lifespan in wild white-faced capuchin monkeys. Philos. Trans. R. Soc. B Biol. Sci. 375 (2020), doi:10.1098/rstb.2019.0494.

61. B. Beck, G. Pourié, Ghrelin, neuropeptide Y, and other feeding-regulatory peptides active in the hippocampus: Role in learning and memory. Nutr. Rev. 71, 541–561 (2013).

62. A. H. Bazzari, H. R. Parri, Neuromodulators and long-term synaptic plasticity in learning and memory: A steered-glutamatergic perspective. Brain Sci. 9 (2019), doi:10.3390/brainsci9110300.

63. D. H. Bergin, Y. Jing, G. Williams, B. G. Mockett, H. Zhang, W. C. Abraham, P. Liu, Safety and neurochemical profiles of acute and sub-chronic oral treatment with agmatine sulfate. Sci. Rep. 9, 1–13 (2019).

64. J. W. Smith, A. T. Evans, B. Costall, J. W. Smythe, Thyroid hormones, brain function and cognition: A brief review. Neurosci. Biobehav. Rev. 26, 45–60 (2002).

65. S. Boinski, K. Sughrue, L. Selvaggi, R. Quatrone, M. Henry, S. Cropp, An expanded test of the ecological model of primate social evolution: competitive regimes and female bonding in three species of squirrel monkeys (Saimiri oerstedii, S. boliviensis, and S. sciureus). Behaviour. 139, 227–261 (2002).

66. J. L. Frechette, K. E. Sieving, S. Boinski, Social and personal information use by squirrel monkeys in assessing predation risk. Am. J. Primatol. 76, 956–966 (2014).

67. A. F. Dixson, Copulatory and postcopulatory sexual selection in primates. Folia Primatol. 89, 258–286 (2018).

68. T. Pinheiro, S. F. Ferrari, M. A. Lopes, Activity budget, diet, and use of space by two groups of squirrel monkeys (Saimiri sciureus) in eastern Amazonia. Primates. 54, 301–308 (2013).

69. P. Izar, A. Stone, S. Carnegie, E. S. Nakai, “Sexual selection, female choice and mating systems” in South American Primates: Comparative Perspectives in the Study of Behavior, Ecology, and Conservation, A. P. Garber, A. Estrada, J. C. Bicca-Marques, E. W. Heymann, K. B. Strier, Eds. (Springer, New York, 2009), pp. 157–189.

70. T. Ubuka, K. Tsutsui, Reproductive neuroendocrinology of mammalian gonadotropin-inhibitory hormone. Reprod. Med. Biol. 18, 225–233 (2019).

71. K. A. Wright, B. W. Wright, S. M. Ford, D. Fragaszy, P. Izar, M. Norconk, T. Masterson, D. G. Hobbs, M. E. Alfaro, J. W. Lynch Alfaro, The effects of ecology and evolutionary history on robust capuchin morphological diversity. Mol. Phylogenet. Evol. 82, 455–466 (2015).

72. T. J. Masterson, Sexual dimorphism and interspecific cranial form in two capuchin species: Cebus albifrons and C. apella. Am. J. Phys. Anthropol. 104, 487–511 (1997).

73. Y. Eshchar, P. Izar, E. Visalberghi, B. Resende, D. Fragaszy, When and where to practice: social influences on the development of nut-cracking in bearded capuchins (Sapajus libidinosus). Anim. Cogn. 19, 605–618 (2016).

74. N. Funato, New insights into cranial synchondrosis development: a mini review. Front. Cell Dev. Biol. 8, 1–9 (2020).

75. M. R. Hasan, M. Takatalo, H. Ma, R. Rice, T. Mustonen, D. P. C. Rice, Rab23 coordinates early osteogenesis by repressing FGF10-PERK1/2 and GLI1. Elife. 9, 1–35 (2020).

76. M. Ansari, G. Poke, Q. Ferry, K. Williamson, R. Aldridge, A. M. Meynert, H. Bengani, C. Y. Chan, H. Kayserili, Ş. Avci, R. C. M. Hennekam, A. K. Lampe, E. Redeker, T. Homfray, A. Ross, M. F. Smeland, S. Mansour, M. J. Parker, J. A. Cook, M. Splitt, R. B. Fisher, A. Fryer, A. C. Magee, A. Wilkie, A. Barnicoat, A. F. Brady, N. S. Cooper, C. Mercer, C. Deshpande, C. P. Bennett, D. T. Pilz, D. Ruddy, D. Cilliers, D. S. Johnson, D. Josifova, E. Rosser, E. M. Thompson, E. Wakeling, E. Kinning, F. Stewart, F. Flinter, K. M. Girisha, H. Cox, H. V. Firth, H. Kingston, J. S. Wee, J. A. Hurst, J. Clayton-Smith, J. Tolmie, J. Vogt, K. Tatton-Brown, K. Chandler, K. Prescott, L. Wilson, M. Behnam, M. McEntagart, R. Davidson, S. A. Lynch, S. Sisodiya, S. G. Mehta, S. A. McKee, S. Mohammed, S. Holden, S. M. Park, S. E. Holder, V. Harrison, V. McConnell, W. K. Lam, A. J. Green, D. Donnai, M. Bitner-Glindzicz, D. E. Donnelly, C. Nellåker, M. S. Taylor, D. R. FitzPatrick, Genetic heterogeneity in Cornelia de Lange syndrome (CdLS) and CdLS-like phenotypes with observed and predicted levels of mosaicism. J. Med. Genet. 51, 659–668 (2014).

77. W. L. Jungers, J. G. Fleagle, Postnatal growth allometry of the extremities in Cebus albifrons and Cebus apella: A longitudinal and comparative study. Am. J. Phys. Anthropol. 53, 471–478 (1980).

78. C. McQueen, M. Towers, Establishing the pattern of the vertebrate limb. Dev. 147 (2020), doi:10.1242/dev.177956.

79. M. B. Costello, D. M. Fragaszy, Prehension in Cebus and Saimiri: I. Grip type and hand preference. Am. J. Primatol. 15, 235–245 (1988).

80. V. Truppa, P. Carducci, G. Sabbatini, Object grasping and manipulation in capuchin monkeys (genera Cebus and Sapajus). Biol. J. Linn. Soc. 127, 563–582 (2019).

81. K. L. Allen, R. F. Kay, Dietary quality and encephalization in platyrrhine primates. Proc. Biol. Sci. 279, 715–21 (2012).

82. L. P. C. dos Santos, “Parâmetros nutricionais da dieta de duas populações de macacos-prego: Sapajus libidinosus no ecótono Cerrado/Caatinga e Sapajus nigritus na Mata Atlântica”, thesis, Universidade de São Paulo, São Paulo (2015).

83. M. G. M. Lima, J. C. Buckner, J. de S. e. Silva-Júnior, A. Aleixo, A. B. Martins, J. P. Boubli, A. Link, I. P. Farias, M. N. da Silva, F. Röhe, H. Queiroz, K. L. Chiou, A. Di Fiore, M. E. Alfaro, J. W. Lynch Alfaro, Capuchin monkey biogeography: understanding Sapajus Pleistocene range expansion and the current sympatry between Cebus and Sapajus. J. Biogeogr. 44, 810–820 (2017).

84. J. A. Chapman, I. Ho, S. Sunkara, S. Luo, G. P. Schroth, D. S. Rokhsar, Meraculous: de novo genome assembly with short paired-end reads. PLoS One. 6, e23501 (2011).

85. G. Marçais, C. Kingsford, A fast, lock-free approach for efficient parallel counting of occurrences of k-mers. Bioinformatics. 27, 764–770 (2011).

86. B. Chevreux, T. Pfisterer, B. Drescher, A. J. Driesel, W. E. G. Müller, T. Wetter, S. Suhai, Using the miraEST assembler for reliable and automated mRNA transcript assembly and SNP detection in sequenced ESTs. Genome Res. 14, 1147–1159 (2004).

87. C. Hahn, L. Bachmann, B. Chevreux, Reconstructing mitochondrial genomes directly from genomic next-generation sequencing reads—a baiting and iterative mapping approach. Nucleic Acids Res. 41, e129–e129 (2013).

88. A. F. A. Smit, R. Hubley, P. Green, RepeatMasker Open-4.0 2013-2015. Available at http://www.repeatmasker.org.

89. A. F. A. Smit, R. Hubley, RepeatModeler Open-1.0. 2008-2015. Available at http://www.repeatmasker.org.

90. M. Stanke, O. Keller, I. Gunduz, A. Hayes, S. Waack, B. Morgenstern, AUGUSTUS: ab initio prediction of alternative transcripts. Nucleic Acids Res. 34, W435–W439 (2006).

91. S. Götz, J. M. García-Gómez, J. Terol, T. D. Williams, S. H. Nagaraj, M. J. Nueda, M. Robles, M. Talón, J. Dopazo, A. Conesa, High-throughput functional annotation and data mining with the Blast2GO suite. Nucleic Acids Res. 36, 3420–3435 (2008).

92. L. Li, C. J. Stoeckert, D. S. Roos, OrthoMCL: identification of ortholog groups for eukaryotic genomes. Genome Res. 13, 2178–2189 (2003).

93. K. Katoh, D. M. Standley, MAFFT multiple sequence alignment software version 7: improvements in performance and usability. Mol. Biol. Evol. 30, 772–780 (2013).

94. R. M. Waterhouse, M. Seppey, F. A. Simão, M. Manni, P. Ioannidis, G. Klioutchnikov, E. V. Kriventseva, E. M. Zdobnov, BUSCO applications from quality assessments to gene prediction and phylogenomics. Mol. Biol. Evol. 35, 543–548 (2018).

95. A. M. Bolger, M. Lohse, B. Usadel, Trimmomatic: a flexible trimmer for Illumina sequence data. Bioinformatics. 30, 2114–2120 (2014).

96. G. W. Vurture, F. J. Sedlazeck, M. Nattestad, C. J. Underwood, H. Fang, J. Gurtowski, M. C. Schatz, GenomeScope: Fast reference-free genome profiling from short reads. Bioinformatics. 33, 2202–2204 (2017).

97. H. Sun, J. Ding, M. Piednoël, K. Schneeberger, FindGSE: Estimating genome size variation within human and Arabidopsis using k-mer frequencies. Bioinformatics. 34, 550–557 (2018).

98. S. Liu, Y. Liu, X. Yang, C. Tong, D. Edwards, I. A. P. Parkin, M. Zhao, J. Ma, J. Yu, S. Huang, X. Wang, J. Wang, K. Lu, Z. Fang, I. Bancroft, T.-J. Yang, Q. Hu, X. Wang, Z. Yue, L. Li, Haojie Yang, J. Wu, Q. Zhou, W. Wang, G. J. King, J. C. Pires, C. Lu, Z. Wu, P. Sampath, Z. Wang, H. Guo, S. Pan, L. Yang, J. Min, D. Zhang, D. Jin, W. Li, H. Belcram, J. Tu, M. Guan, C. Qi, D. Du, J. Li, L. Jiang, J. Batley, A. G. Sharpe, B.-S. Park, P. Ruperao, F. Cheng, N. E. Waminal, Y. Huang, C. Dong, L. Wang, J. Li, Z. Hu, M. Zhuang, Y. Huang, J. Huang, J. Shi, D. Mei, J. Liu, T.-H. Lee, J. Wang, H. Jin, Z. Li, X. Li, J. Zhang, L. Xiao, Y. Zhou, Z. Liu, X. Liu, R. Qin, X. Tang, W. Liu, Y. Wang, Y. Zhang, J. Lee, H. H. Kim, F. Denoeud, X. Xu, X. Liang, W. Hua, X. Wang, J. Wang, B. Chalhoub, A. H. Paterson, The Brassica oleracea genome reveals the asymmetrical evolution of polyploid genomes. Nat. Commun. 5, 1–11 (2014).

99. L. Song, L. Florea, Rcorrector: efficient and accurate error correction for Illumina RNA-seq reads. Gigascience. 4, s13742–015 (2015).

100. C. Quast, E. Pruesse, P. Yilmaz, J. Gerken, T. Schweer, P. Yarza, J. Peplies, F. O. Glöckner, The SILVA ribosomal RNA gene database project: improved data processing and web-based tools. Nucleic Acids Res. 41, D590–D596 (2012).

101. B. Langmead, S. L. Salzberg, Fast gapped-read alignment with Bowtie 2. Nat. Methods. 9, 357–359 (2012).

102. M. Johnson, I. Zaretskaya, Y. Raytselis, Y. Merezhuk, S. McGinnis, T. L. Madden, NCBI BLAST: a better web interface. Nucleic Acids Res. 36, W5–W9 (2008).

103. M. G. Grabherr, B. J. Haas, M. Yassour, J. Z. Levin, D. A. Thompson, I. Amit, X. Adiconis, L. Fan, R. Raychowdhury, Q. Zeng, Z. Chen, E. Mauceli, N. Hacohen, A. Gnirke, N. Rhind, F. di Palma, B. W. Birren, C. Nusbaum, K. Lindblad-Toh, N. Friedman, Regev, Trinity: reconstructing a full-length transcriptome without a genome from RNA-Seq data. Nat. Biotechnol. 29, 644–652 (2011).

104. A. Dobin, C. A. Davis, F. Schlesinger, J. Drenkow, C. Zaleski, S. Jha, P. Batut, M. Chaisson, T. R. Gingeras, STAR: ultrafast universal RNA-seq aligner. Bioinformatics. 29, 15–21 (2013).

105. C. Trapnell, A. Roberts, L. Goff, G. Pertea, D. Kim, D. R. Kelley, H. Pimentel, S. L. Salzberg, J. L. Rinn, L. Pachter, Differential gene and transcript expression analysis of RNA-seq experiments with TopHat and Cufflinks. Nat. Protoc. 7, 562–578 (2012).

106. M. A. Campbell, B. J. Haas, J. P. Hamilton, S. M. Mount, C. R. Buell, Comprehensive analysis of alternative splicing in rice and comparative analyses with Arabidopsis. BMC Genomics. 7, 1–17 (2006).

107. M. Carruthers, A. A. Yurchenko, J. J. Augley, C. E. Adams, P. Herzyk, K. R. Elmer, De novo transcriptome assembly, annotation and comparison of four ecological and evolutionary model salmonid fish species. BMC Genomics. 19, 1–17 (2018).

108. W. Li, A. Godzik, CD-hit: a fast program for clustering and comparing large sets of protein or nucleotide sequences. Bioinformatics. 22, 1658–1659 (2006).

109. B. Buchfink, C. Xie, D. H. Huson, Fast and sensitive protein alignment using DIAMOND. Nat. Methods. 12, 59–60 (2015).

110. W. J. Kent, BLAT—the BLAST-like alignment tool. Genome Res. 12, 656–664 (2002).

111. W. Bao, K. K. Kojima, O. Kohany, Repbase Update, a database of repetitive elements in eukaryotic genomes. Mob. DNA. 6, 1–6 (2015).

112. A. Piovesan, M. C. Pelleri, F. Antonaros, P. Strippoli, M. Caracausi, L. Vitale, On the length, weight and GC content of the human genome. BMC Res. Notes. 12, 1–7 (2019).

113. P. Jones, D. Binns, H.-Y. Chang, M. Fraser, W. Li, C. McAnulla, H. McWilliam, J. Maslen, A. Mitchell, G. Nuka, S. Pesseat, A. F. Quinn, A. Sangrador-Vegas, M. Scheremetjew, S.-Y. Yong, R. Lopez, S. Hunter, InterProScan 5: genome-scale protein function classification. Bioinformatics. 30, 1236–1240 (2014).

114. A. C. Beichman, K. P. Koepfli, G. Li, W. Murphy, P. Dobrynin, S. Kliver, M. T. Tinker, M. J. Murray, J. Johnson, K. Lindblad-Toh, E. K. Karlsson, K. E. Lohmueller, R. K. Wayne, Aquatic adaptation and depleted diversity: a deep dive into the genomes of the sea otter and giant otter. Mol. Biol. Evol. 36, 2631–2655 (2019).

115. S. Mallick, S. Gnerre, P. Muller, D. Reich, The difficulty of avoiding false positives in genome scans for natural selection. Genome Res. 19, 922–933 (2009).

116. W. Fletcher, Z. Yang, The effect of insertions, deletions, and alignment errors on the branch-site test of positive selection. Mol. Biol. Evol. 27, 2257–2267 (2010).

117. P. Markova-Raina, D. Petrov, High sensitivity to aligner and high rate of false positives in the estimates of positive selection in the 12 Drosophila genomes. Genome Res. 21, 863–874 (2011).

118. Z. Yang, M. Dos Reis, Statistical properties of the branch-site test of positive selection. Mol. Biol. Evol. 28, 1217–1228 (2011).

119. G. Jordan, N. Goldman, The effects of alignment error and alignment filtering on the sitewise detection of positive selection. Mol. Biol. Evol. 29, 1125–1139 (2012).

120. T. B. Sackton, Studying natural selection in the era of ubiquitous genomes. Trends Genet. 36, 792–803 (2020).

121. M. S. Springer, R. W. Meredith, J. Gatesy, C. A. Emerling, J. Park, C. A. Fisher, W. J. Murphy, Macroevolutionary dynamics and historical biogeography of primate diversification inferred from a species supermatrix. PLoS One. 7, e49521 (2012).

122. E. Maldonado, D. Almeida, T. Escalona, I. Khan, V. Vasconcelos, A. Antunes, LMAP: Lightweight Multigene Analyses in PAML. BMC Bioinformatics. 17, 1–46 (2016).

123. R. Nielsen, C. Bustamante, A. G. Clark, S. Glanowski, T. B. Sackton, M. J. Hubisz, A. Fledel-Alon, D. M. Tanenbaum, D. Civello, T. J. White, J. J. Sninsky, M. D. Adams, M. Cargill, A scan for positively selected genes in the genomes of humans and chimpanzees. PLoS Biol. 3, 0976–0985 (2005).

124. T. S. Mikkelsen, L. W. Hillier, E. E. Eichler, M. C. Zody, D. B. Jaffe, S. P. Yang, W. Enard, I. Hellmann, K. Lindblad-Toh, T. K. Altheide, N. Archidiacono, P. Bork, J. Butler, J. L. Chang, Z. Cheng, A. T. Chinwalla, P. Dejong, K. D. Delehaunty, C. C. Fronick, L. L. Fulton, Y. Gilad, G. Glusman, S. Gnerre, T. A. Graves, T. Hayakawa, K. E. Hayden, X. Huang, H. Ji, W. J. Kent, M. C. King, E. J. Kulbokas, M. K. Lee, G. Liu, C. Lopez-Otin, K. D. Makova, O. Man, E. R. Mardis, E. Mauceli, T. L. Miner, W. E. Nash, J. O. Nelson, S. Pääbo, N. J. Patterson, C. S. Pohl, K. S. Pollard, K. Prüfer, X. S. Puente, D. Reich, M. Rocchi, K. Rosenbloom, M. Ruvolo, D. J. Richter, S. F. Schaffner, A. F. A. Smit, S. M. Smith, M. Suyama, J. Taylor, D. Torrents, E. Tuzun, A. Varki, G. Velasco, M. Ventura, J. W. Wallis, M. C. Wendl, R. K. Wilson, E. S. Lander, R. H. Waterston, Initial sequence of the chimpanzee genome and comparison with the human genome. Nature. 437, 69–87 (2005).

125. G. C. Nickel, D. L. Tefft, K. Goglin, M. D. Adams, An empirical test for branch-specific positive selection. Genetics. 179, 2183–2193 (2008).

126. G. Jordan, “Analysis of alignment error and sitewise constraint in mammalian comparative genomics”, thesis, University of Cambridge, Cambridge (2011).

127. Y. Benjamini, Y. Hochberg, Controlling the false discovery rate: a practical and powerful approach to multiple testing. J. R. Stat. Soc. Ser. B. 57, 289–300 (1995).

128. G. Stelzer, N. Rosen, I. Plaschkes, S. Zimmerman, M. Twik, S. Fishilevich, T. I. Stein, R. Nudel, I. Lieder, Y. Mazor, S. Kaplan, D. Dahary, D. Warshawsky, Y. Guan-Golan, A. Kohn, N. Rappaport, M. Safran, D. Lancet, The GeneCards suite: from gene data mining to disease genome sequence analyses. Curr. Protoc. Bioinforma. 54, 1–30 (2016).

129. M. Simões-Costa, M. Stone, M. E. Bronner, Axud1 integrates Wnt signaling and transcriptional inputs to drive neural crest formation. Dev. Cell. 34, 544–554 (2015).

130. E. Flex, M. Jaiswal, F. Pantaleoni, S. Martinelli, M. Strullu, E. K. Fansa, A. Caye, A. De Luca, F. Lepri, R. Dvorsky, L. Pannone, S. Paolacci, S. C. Zhang, V. Fodale, G. Bocchinfuso, C. Rossi, E. M. M. Burkitt-Wright, A. Farrotti, E. Stellacci, S. Cecchetti, R. Ferese, L. Bottero, S. Castro, O. Fenneteau, B. Brethon, M. Sanchez, A. E. Roberts, H. G. Yntema, I. Van Der Burgt, P. Cianci, M. L. Bondeson, M. C. Digilio, G. Zampino, B. Kerr, Y. K. Aoki, M. L. Loh, A. Palleschi, E. Di Schiavi, A. Caré, A. Selicorni, B. Dallapiccola, I. C. Cirstea, L. Stella, M. Zenker, B. D. Gelb, H. Cavé, M. R. Ahmadian, M. Tartaglia, Activating mutations in RRAS underlie a phenotype within the RASopathy spectrum and contribute to leukaemogenesis. Hum. Mol. Genet. 23, 4315–4327 (2014).

131. G. Grimaldi, B. Vagaska, O. Ievglevskyi, E. Kondratskaya, J. Glover, J. Matthews, Loss of Tiparp results in aberrant layering of the cerebral cortex. eNeuro. 6, 1–15 (2019).

132. M. Kono, M. L. Allende, R. L. Proia, Sphingosine-1-phosphate regulation of mammalian development. Biochim. Biophys. Acta. 1781, 435–441 (2008).

133. G. He, S. Tavella, K. P. Hanley, M. Self, G. Oliver, R. Grifone, N. Hanley, C. Ward, N. Bobola, Inactivation of Six2 in mouse identifies a novel genetic mechanism controlling development and growth of the cranial base. Dev. Biol. 344, 720–730 (2010).

134. P. Wend, K. Wend, S. A. Krum, G. A. Miranda-Carboni, The role of WNT10B in physiology and disease. Acta Physiol. 204, 34–51 (2012).

135. K. Kania, F. Colella, A. H. K. Riemen, H. Wang, K. A. Howard, T. Aigner, F. Dell’Accio, T. D. Capellini, A. J. Roelofs, C. De Bari, Regulation of Gdf5 expression in joint remodelling, repair and osteoarthritis. Sci. Rep. 10, 1–11 (2020).

136. P. Buxton, C. Edwards, C. W. Archer, P. Francis-West, Growth/differentiation factor-5 (GDF-5) and skeletal development. J. Bone Jt. Surg. 83, S23–S30 (2001).

137. S. Beck-Cormier, M. Escande, C. Souilhol, S. Vandormael-Pournin, S. Sourice, P. Pilet, C. Babinet, M. Cohen-Tannoudji, Notchless is required for axial skeleton formation in mice. PLoS One. 9, 1–10 (2014).

138. C. L. Smith, M. D. Tallquist, PDGF function in diverse neural crest cell populations. Cell Adhes. Migr. 4, 561–566 (2010).

139. C. Niehrs, Function and biological roles of the Dickkopf family of Wnt modulators. Oncogene. 25, 7469–7481 (2006).

140. J. E. Lee, J. G. Gleeson, Cilia in the nervous system: Linking cilia function and neurodevelopmental disorders. Curr. Opin. Neurol. 24, 98–105 (2011).

141. E. N. Firat-Karalar, The ciliopathy gene product Cep290 is required for primary cilium formation and microtubule network organization. Turkish J. Biol. 42, 371–381 (2018).

142. S. J. Neufeld, F. Wang, J. Cobb, Genetic interactions between Shox2 and Hox genes during the regional growth and development of the mouse limb. Genetics. 198, 1117–1126 (2014).

143. C. Tickle, M. Towers, Sonic hedgehog signaling in limb development. Front. Cell Dev. Biol. 5, 1–19 (2017).

144. J. Zákány, M. Kmita, D. Duboule, A dual role for Hox genes in limb anterior-posterior asymmetry. Science (80). 304, 1669–1672 (2004).

145. L. A. Wyngaarden, S. Hopyan, Plasticity of proximal-distal cell fate in the mammalian limb bud. Dev. Biol. 313, 225–233 (2008).

146. Z. Iqbal, P. Cejudo-Martin, A. de Brouwer, B. van der Zwaag, P. Ruiz-Lozano, M. C. Scimia, J. D. Lindsey, R. Weinreb, B. Albrecht, A. Megarbane, Y. Alanay, Z. Ben-Neriah, M. Amenduni, R. Artuso, J. A. Veltman, E. van Beusekom, A. Oudakker, J. L. Millán, R. Hennekam, B. Hamel, S. A. Courtneidge, H. van Bokhoven, Disruption of the podosome adaptor protein TKS4 (SH3PXD2B) causes the skeletal dysplasia, eye, and cardiac abnormalities of Frank-Ter Haar syndrome. Am. J. Hum. Genet. 86, 254–261 (2010).

147. N. Funato, Y. Taga, L. E. Laurie, C. Tometsuka, M. Kusubata, K. Ogawa-Goto, The transcription factor HAND1 is involved in cortical bone mass through the regulation of collagen expression. Int. J. Mol. Sci. 21, 1–15 (2020).

148. V. Ouellet, P. M. Siegel, CCN3 modulates bone turnover and is a novel regulator of skeletal metastasis. J. Cell Commun. Signal. 6, 73–85 (2012).

149. R. Haraguchi, R. Kitazawa, K. Mori, R. Tachibana, H. Kiyonari, Y. Imai, T. Abe, S. Kitazawa, SFRP4-dependent Wnt signal modulation is critical for bone remodeling during postnatal development and age-related bone loss. Sci. Rep. 6, 1–14 (2016).

150. P. O. Simsek Kiper, H. Saito, F. Gori, S. Unger, E. Hesse, K. Yamana, R. Kiviranta, N. Solban, J. Liu, R. Brommage, K. Boduroglu, L. Bonafé, B. Campos-Xavier, E. Dikoglu, R. Eastell, F. Gossiel, K. Harshman, G. Nishimura, K. M. Girisha, B. J. Stevenson, H. Takita, C. Rivolta, A. Superti-Furga, R. Baron, Cortical-bone fragility — insights from sFRP4 deficiency in Pyle’s disease. N. Engl. J. Med. 374, 2553–2562 (2016).

151. S. Sipione, J. Monyror, D. Galleguillos, N. Steinberg, V. Kadam, Gangliosides in the brain: physiology, pathophysiology and therapeutic applications. Front. Neurosci. 14, 1–24 (2020).

152. E. C. Pandolfi, K. J. Tonsfeldt, H. M. Hoffmann, P. L. Mellon, Deletion of the homeodomain protein Six6 from GnRH neurons decreases GnRH gene expression, resulting in infertility. Endocrinology. 160, 2151–2164 (2019).

153. A. Vera, K. Stanic, H. Montecinos, M. Torrejón, S. Marcellini, T. Caprile, SCO-spondin from embryonic cerebrospinal fluid is required for neurogenesis during early brain development. Front. Cell. Neurosci. 7, 1–14 (2013).

154. K. H. Flippo, S. Strack, Mitochondrial dynamics in neuronal injury, development and plasticity. J. Cell Sci. 130, 671–681 (2017).

155. J. H. Jhamandas, V. Goncharuk, Role of neuropeptide FF in central cardiovascular and neuroendocrine regulation. Front. Endocrinol. 4, 1–6 (2013).

156. E. Angelopoulou, C. Quignon, L. J. Kriegsfeld, V. Simonneaux, Functional implications of RFRP-3 in the central control of daily and seasonal rhythms in reproduction. Front. Endocrinol. 10, 1–15 (2019).

157. L. J. Kriegsfeld, K. J. Jennings, G. E. Bentley, K. Tsutsui, Gonadotrophin-inhibitory hormone and its mammalian orthologue RFamide-related peptide-3: Discovery and functional implications for reproduction and stress. J. Neuroendocrinol. 30, 1–13 (2018).

158. Y. Zhao, J. Wu, X. Wang, H. Jia, D.-N. Chen, J.-D. Li, Prokineticins and their G proteincoupled receptors in health and disease. Prog. Mol. Biol. Transl. Sci. 161, 149–179 (2019).

159. C. Jia, M. P. Keasey, H. M. Malone, C. Lovins, R. R. Sante, V. Razskazovskiy, T. Hagg, Vitronectin from brain pericytes promotes adult forebrain neurogenesis by stimulating CNTF. Exp. Neurol. 312, 20–32 (2019).

160. P. O. Frappart, Y. Lee, J. Lamont, P. J. McKinnon, BRCA2 is required for neurogenesis and suppression of medulloblastoma. EMBO J. 26, 2732–2742 (2007).

161. L. Lo, E. L. Dormand, D. J. Anderson, Late-emigrating neural crest cells in the roof plate are restricted to a sensory fate by GDF7. Proc. Natl. Acad. Sci. U. S. A. 102, 7192–7197 (2005).

162. P. Joshi, C. Bodnya, M. L. Rasmussen, A. I. Romero-Morales, A. Bright, V. Gama, Modeling the function of BAX and BAK in early human brain development using iPSC-derived systems. Cell Death Dis. 11 (2020), doi:10.1038/s41419-020-03002-x.

163. J. Xie, Y. Jin, G. Wang, The role of SCF ubiquitin-ligase complex at the beginning of life. Reprod. Biol. Endocrinol. 17, 1–9 (2019).

164. C. Wang, X. Kang, L. Zhou, Z. Chai, Q. Wu, R. Huang, H. Xu, M. Hu, X. Sun, S. Sun, J. Li, R. Jiao, P. Zuo, L. Zheng, Z. Yue, Z. Zhou, Synaptotagmin-11 is a critical mediator of parkin-linked neurotoxicity and Parkinson’s disease-like pathology. Nat. Commun. 9, 1–14 (2018).

165. M. Jakimiec, J. Paprocka, R. Smigiel, CDKL5 deficiency disorder—a complex epileptic encephalopathy. Brain Sci. 10, 1–9 (2020).

166. S. Chauvin, A. Sobel, Neuronal stathmins: A family of phosphoproteins cooperating for neuronal development, plasticity and regeneration. Prog. Neurobiol. 126, 1–18 (2015).

167. W. Jiang, M. Wei, M. Liu, Y. Pan, D. Cao, X. Yang, C. Zhang, Identification of protein tyrosine phosphatase receptor type O (PTPRO) as a synaptic adhesion molecule that promotes synapse formation. J. Neurosci. 37, 9828–9843 (2017).

168. Y. Ranjbar-Slamloo, Z. Fazlali, Dopamine and noradrenaline in the brain; overlapping or dissociate functions? Front. Mol. Neurosci. 12, 1–8 (2020).

169. E. Boudin, T. R. de Jong, T. C. R. Prickett, B. Lapauw, K. Toye, V. Van Hoof, I. Luyckx, Verstraeten, H. S. A. Heymans, E. Dulfer, L. Van Laer, I. R. Berry, A. Dobbie, E. Blair, Loeys, E. A. Espiner, J. M. Wit, W. Van Hul, P. Houpt, G. R. Mortier, Bi-allelic loss-of-function mutations in the NPR-C receptor result in enhanced growth and connective tissue abnormalities. Am. J. Hum. Genet. 103, 288–295 (2018).

170. Y. Guo, W. Pan, S. Liu, Z. Shen, Y. Xu, L. Hu, ERK/MAPK signalling pathway and tumorigenesis (Review). Exp. Ther. Med., 1997–2007 (2020).

171. W. A. Buttemer, D. Abele, D. Costantini, From bivalves to birds: Oxidative stress and longevity. Funct. Ecol. 24, 971–983 (2010).

172. A. P. Halestrap, The SLC16 gene family; structure, role and regulation in health and disease. Mol. Aspects Med. 34, 337–349 (2013).

