## Supplementary Materials for "Signatures of adaptive evolution in platyrrhine primate genomes"

**This PDF file includes:**

Section 1: Extended methods & results: Robust capuchin reference genome
Section 2: Extended methods & results: Signatures of selection in platyrrhine genomes
Section 3: Extended results: Robust capuchins (*Sapajus*; H1) positive selection results
Section 4: Extended results: Gracile capuchins (*Cebus*; H2) positive selection results
Section 5: Extended results: Ancestral Cebinae (H3) positive selection results
Section 6: Extended results: Across-capuchins (H3a) positive selection results
Section 7: Extended results: Ancestral Cebidae (H4) positive selection results
Section 8: Extended results: Squirrel monkeys (*Saimiri*; H5) positive selection results
Figs. S1 to S3
Tables S1–S3, S5–S7, S10, and S23
References (94 to 172)

**Other Supplementary Materials for this manuscript include the following:**

Tables S4, S8, S9, S11–S22, and S24–S44

### 1) Extended methods & results: Robust capuchin reference genome

1.1 Genome assembly: versions and accessions

The version of the genome assembly used in this study, Sape_Mango_1.0, was uploaded to a Zenodo repository (see data availability). An assembly (Sape_Mango_1.1) with minor modifications including the removal of two short scaffolds and the addition of the mitochondrial genome assembly was uploaded to NCBI under the accession JAGHVQ. The BioProject and BioSample NCBI accessions for this project and sample (Mango) are PRJNA717806 and SAMN18511585. See Table S1 for NCBI's sequence read archive (SRA) accessions for the raw short-read data for the shotgun and Chicago libraries.

1.2 Genome completeness

We evaluated completeness of the genome assembly by its estimated gene content using CEGMA v2.5 (Conserved Eukaryotic Genes Mapping Approach) (*15*) and BUSCO v3.0.2 (Benchmarking Universal Single Copy Orthologs) (*14*), which calculate the proportion of 248 core eukaryotic genes (CEGs) or 6,192 Euarchontoglires-specific conserved single copy orthologs, respectively, that were either complete, fragmented, or missing. We ran BUSCO starting with gene finding parameters optimised for the human genome from the *ab initio* HMM-based gene predictor, Augustus v3.3 (*90*). We specified the "long" option, instructing BUSCO to use the initial gene models it creates to then retrain the human HMM search model and optimise the parameters for the robust capuchin genome (*94*). The internal training that BUSCO performs is an automated five-round Augustus gene finder training pipeline. Our goal here was two-fold; improve BUSCO's ability to estimate gene content, and produce a trained HMM for capuchins to be used in the first pass of genome annotation pipeline (see below). We identified 91.5% (N = 5,666) of BUSCO's Euarchontoglires-specific conserved single copy orthologs (N = 6,192) in the assembly including 85% (N = 5,264) complete (with 0.6% duplicated) and 6.5% (N = 402) fragmented. We detect 90.3% (N = 224) of CEGMA's CEGs (N = 248) in the final assembly in at least partial status with 77.4% (N = 192) as complete CEGs.

1.3 Genome size estimation

We processed the raw, shotgun short read pairs to get clean, nuclear only, endogenous reads for genome size estimation. First, we trimmed for quality and adapter contamination using trimmomatic v.0.36 (*95*) with the options "ILLUMINACLIP:2:30:10 LEADING:3 TRAILING:3

SLIDINGWINDOW:4:20 MINLEN:36”. We then screened for vectors and contaminants using Kraken2 (https://github.com/DerrickWood/kraken2) following the standard build (viral, archaea, bacteria and UniVec_Core libraries) except excluding the human library given the similarity to capuchins. We removed read pairs that mapped to our mitochondrial genome assembly (see below) with a minimum identity of 85% using bbmap.sh from bbmap tools v.37.99 (https://sourceforge.net/projects/bbmap). We assessed quality metrics for the trimmed, filtered reads using FastQC (https://www.bioinformatics.babraham.ac.uk/projects/fastqc). We retained a total of ~ 2.1 billion reads (266.5 billion bases) for genome size estimation. We performed *k*-mer counting with the quality-filtered, clean reads with Jellyfish v.2.2.6 (*85*), generating a *k*-mer frequency distribution of 31-mers, and then we estimated genome size using four approaches.

The first method estimated genome size and other parameters based on a mixture model of four negative binominal distributions implemented by the GenomeScope 1.0 (*96*), which calculates the relative abundance of heterozygous and homozygous unique and two-copy sequences to estimate the heterozygosity and repeat fraction as well as the error rate. We did not apply a maximum *k*-mer frequency filter in GenomeScope as we had already removed sequences from contaminant or organelle sources. GenomeScope estimated a genome size of 2,917,676,754 bp with approximately 77.6% unique content (22.4% repeat), a heterozygosity level of 0.287% and an error rate of 0.0824% (Figure S1). For the second method, we used the Jellyfish stats command to calculate the total number of *k*-mers with a minimum frequency of 11 to exclude likely error *k*-mers (which was estimated from the *k*-mer frequency distribution) and then divided this total by the *k*-mer peak frequency (198263930448/66) for an estimate of 3,003,998,946 bp. The third method we used was implemented in the R package findGSE v.1.94 (*97*) which estimated a genome size of 3,029,414,613 bp with a heterozygosity rate of 0.25423% and 76.5% unique content (23.5% repeat). Finally, we also manually calculated genome size using the formula ((R*(L-K+1)-B)/M) (*98*) where R is the total number of reads, M is the *k*-mer peak frequency, K is the *k*-mer size, L is the average read length, and B the number of *k*-mers at very low frequency (< 4) (which is a corrective factor for sequencing errors) [(2111490333*(126.222-31+1)-4225744143)/66] giving an estimated size of 3,014,334,525 bp. Thus, across the four methods, the estimated haploid genome length for our *Sapajus apella* reference individual was between 2,918 and 3,029 Mbp (Table S2).

We also mapped the quality filtered, clean shotgun reads back to the genome assembly as an assessment of assembly quality with a minimum identity of 90% using bbmap.sh from bbmap tools v37.99, which showed 90% of reads (and bases) mapped successfully. A further assessment of genome assembly quality is contained within the rnaQUAST (*17*) analyses performed on the seven transcript assemblies (see the next section), which indicated that upwards of 94% of the transcripts in the various assemblies aligned to the genome with an average aligned percentage of greater than 92.7% (Table S4).

#### 1.4 RNAseq: filtering & trimming

We assessed quality metrics for the raw sequence data for each of the 17 tissues (temporal lobe, cerebellum, cerebrum, midbrain, hippocampus, pituitary, thymus, bone marrow, mesenteric lymph node (LN), skeletal muscle, aorta, ovary, lung, kidney, liver, duodenum, and colon) using FastQC (https://www.bioinformatics.babraham.ac.uk/projects/fastqc) and then combined all samples for downstream analyses. We used rCorrector (*99*) to correct for rare *k*-mers as they can adversely impact transcriptome assembly using a De Bruijn Graph approach and are often due to sequencing error in a deeply sequenced data set. Read pairs for which a read was deemed unfixable by rCorrector were flagged and subsequently removed with a python script from the Harvard Informatics GitHub repository TranscriptomeAssemblyTools (https://github.com/harvardinformatics/TranscriptomeAssemblyTools). We trimmed for adapters and very low-quality bases (phred < 5) and discarded reads shorter than 36 bp in length using TrimGalore v0.4.4 (https://www.bioinformatics.babraham.ac.uk/projects/trim_galore). We then mapped the trimmed reads to an rRNA database (Vertebrata SSU and LSU downloaded from SILVA; (*100*)) using Bowtie2 v2.3.4 (*101*) with the "very-sensitive-local option", and retained only the unmapped read pairs. We reassessed quality metrics for the trimmed, filtered reads using FastQC which revealed two overrepresented sequences. We used BLAST (*102*) to identify these as rRNA sequences which were not filtered by mapping to the SILVA database and we used bbduk.sh from bbmap tools v37.99 (https://sourceforge.net/projects/bbmap) to remove them. We then performed default trimmomatic quality filtering and *in silico* normalisation with Trinity v.2.5.1 (*103*), retaining ~ 341 million read pairs (95 billion bp) after these trimming and quality filtering steps, and 27 million normalised read pairs (Table S3).

#### 1.5 RNAseq: Transcriptome assembly

We assembled a *de novo* transcriptome with Trinity v.2.5.1 (*103*) using the normalised, quality-filtered RNAseq read pairs with the Jaccard clip option, referred to as TrinDNv1 (Table S4). We estimated abundance with Trinity using RSEM to filter isoforms with < 1% expression levels for that gene to generate the final *de novo* transcriptome assembly for downstream analyses, referred to as TrinDNv2. We mapped the normalised, quality-filtered RNAseq reads to the reference genome assembly using STAR v020201 (*104*) with the following settings: “outFilterMismatchNmax 999, outFilterMismatchNoverLmax 0.06, outFilterMultimapNmax 5, alignIntronMin 20, alignIntronMax 500000, outFilterIntronMotifs RemoveNoncanonical, alignEndsType EndToEnd.” We then assembled a reference-based transcriptome from the aligned RNAseq reads using Cufflinks v2.2.1 (*105*) with the option “-I 500000” (max. intron length), referred to as Cuffv1.

We combined the Trinity *de novo* and Cufflinks reference-based transcript assemblies and, as a preliminary assessment of quality, we used Bowtie2 (*101*) to map the normalised, quality-filtered reads to the combined transcript assembly. Overall, 99.44% of the reads aligned successfully, which is extremely high quality. We then built a comprehensive transcriptome database with PASA v2.2.0 (*106*) using both the Trinity *de novo* (TrinDNv2) and Cufflinks reference-based (Cuffv1) transcript assemblies, referred to as the PASAv1 assembly with 684,426 transcripts (Table S4).

In addition, we followed Carruthers et al. (*107*) to generate a high quality non-redundant RNAseq transcript assembly (referred to as NRv1) to aid gene prediction in the final iteration of the genome annotation pipeline with Maker v3.01.02 (*19*) (see below). We generated a new genome-guided transcript assembly with Trinity v.2.5.1 using the normalised, quality-filtered RNAseq reads mapped to the reference genome assembly with STAR v02020 (see above for Cuffv1 assembly) as input and a max intron length setting of 500,000 bp, referred to as TrinGGv1. We built a second transcriptome database with PASA v2.2.0 using the TrinDNv2, Cuffv1, and Trinity genome-guided (TrinGGv1) transcript assemblies, referred to as the PASAv2 assembly with 744,790 transcripts (Table S4). We then used TransDecoder v5.5.0 (https://transdecoder.github.io) to identify likely coding regions within the PASAv2 transcript assembly, remove transcripts with open reading frames (ORFs) < 210 bp (70 amino acids) in length, and then select the single best ORF per transcript (–single_best_orf). We further reduced redundancy in the remaining transcripts by using CD-Hit v4.7 (*108*) to cluster highly similar sequences using an amino acid sequence identity threshold of 1.00. We calculated coverage against the NCBI gracile capuchin (*Cebus capucinus imitator*) proteins database (GCF_ 001604975.1) using Diamond v0.9.24.125 (*109*) with the options “blastx, max_target_seqs 1, evalue 1e-3, more-sensitive” to assess how successfully our set of non-redundant CDS transcripts were reconstructed to full- or near full-length. We used the “analyze_blastPlus_topHit_coverage.pl” script from the Trinity package (v2.5.1) to identify the number of aligned transcripts across varying length thresholds, and retained all non-redundant CDS transcripts with an alignment length > 30%. As these steps would inadvertently remove transcripts that do not align to the *Cebus* protein set and thus exclude genes that are, for example, missing from the *Cebus* genome assembly, we also performed the Diamond search against a combined human and *Cebus* protein set. The results, however, were highly similar with slightly fewer transcripts retained in the combined set and we proceeded with the *Cebus* only results. Finally, we used this set of length-filtered non-redundant CDS transcripts to generate our final non-redundant (NRv1) transcript assembly for the ultimate iteration of Maker by filtering the equivalent nucleotide transcripts (which include non-CDS sequence) from the original PASAv2 transcript assembly file, with 73,436 transcripts in this final assembly (Table S4). A workflow summary graphic for the RNAseq filtering and assembly steps is shown in Figure S2.

We ran rnaQUAST v1.5.0 (*17*) with BUSCO v3.0.2 (*14*) in transcriptome mode using the Euarchontoglires-specific single copy orthologs database (6,192 orthologs) and with the BLAT v.36x2 alignment tool (*110*) to align the transcripts to the reference genome to check quality metrics and completeness of the seven transcriptome assemblies: TrinDNv1 (Trinity *de novo* assembly); TrinDNv2 (abundance filtered Trinity *de novo* assembly); Cuffv1 (Cufflinks reference based assembly); PASAv1 (PASA assembly with TrinDNv2 and Cuffv1 as input); TrinGGv1 (Trinity genome guided assembly); PASAv2 (PASA assembly v2 with TrinDNv2, Cuffv1, and TrinGGv1 as input); and NRv1 (non-redundant transcript assembly).

Results from the rnaQUAST run indicated the final assemblies used in downstream analyses (TrinDNv2, PASAv1, and NRv1) were high quality, near complete transcriptomes (~ 96 to 97% complete). The NRv1 assembly shows the largest percentage of transcripts longer than 500 bp and 1,000 bp, and the longest average transcript length, transcript N50, and average aligned length (Table S4). Although NRv1 contains less than 10% the number of transcripts contained in some of the larger assemblies including the input assembly (PASAv2), completeness of this transcript assembly as assessed by BUSCO revealed only a minor reduction in the proportion of BUSCO's Euarchontoglires-specific conserved single copy orthologs recovered (97% in PASAv2 vs. 95.9% in NRv1). Comparing our non-redundant transcript assembly (NRv1) with the metrics for the *Cebus capucinus imitator* genome annotation by NCBI (annotation release 100) adds further support for the quality of this transcriptome; for example, average length of *Cebus* mRNA transcripts is 3,513 bp, which is highly similar to the 3,505 bp recovered for the average length of the transcripts in our NRv1 transcriptome, and similarly, the number of exons per transcript and the mean length of exons in coding transcripts are also highly comparable (11.38 vs. 10.25 and 332 vs. 346 bp in the NCBI annotation vs. NRv1, respectively).

#### 1.6 Repeat content

To assess the repeat content of the robust capuchin genome, we first performed a homology-based repeat annotation of our genome assembly with RepeatMasker v4.0.7 (*88*) using the RepBase RepeatMasker library (dc20170127-rb20170127) (*111*) with the options "-sp 'Sapajus apella' -s -nolow -no_is -cutoff 255 -frag 20000", skipping the annotation of low complexity and simple repeats in this first iteration. Using this masked genome as input, we then performed *de novo* modelling of unknown repeat elements using RepeatModeler v1.0.11 (*89*) and ran RepeatMasker again with this *de novo* repeat library with the options "-cutoff 255 -frag 20000", annotating low complexity and simple repeats this round, to create a twice-masked genome. Finally, we combined the list of repeat regions found in the genome assembly (.cat files) from both masking runs and ran ProcessRepeats from RepeatMasker to summarise all annotated repeats in the genome.

The homology-based repeat identification using known RepBase elements annotated 42.15% of the genome assembly as interspersed repeats i.e., transposable elements (TEs) including DNA transposons and retrotransposons (long terminal repeat (LTR) elements, long interspersed nuclear elements (LINEs), and short interspersed elements (SINEs)) (Table S5). This masking run also estimated a GC content of 40.08%, which is very similar to human whole diploid genome GC content of 40.09% (*112*). *De novo* repeat identification using the RepeatModeler library provided a small improvement to the annotations of several classes of transposable elements, recovering an additional 0.87% (21.8 Mbp) of the assembly as *de novo* interspersed repeats. After combining and summarising all repeat regions found in the assembly from both masking runs, total interspersed content of the genome was 43.02% (1.06 Gbp), and total annotated repeat content (including transposable elements as well as small RNA, satellites, simple repeats, and low complexity repeats) was 44.63% (1.12 Gbp). For TEs, LINEs (LINE1 in particular) comprised the greatest genome length, and the greatest number of TEs were SINEs (particularly ALUs). Non-interspersed repeat elements including small RNA elements, satellites, simple repeats, and low complexity repeats comprised 1.61% of the assembly. Information on the homology-based, *de novo*, and combined repeat annotations can be found in Table S5.

#### 1.7 Genome annotation

We annotated the robust capuchin genome assembly in three iterations of Maker v3.01.02 (*18*, *19*) to predict gene models using both homology-based and *ab initio* gene modelling. For the first pass of the Maker pipeline, we incorporated: (i) direct evidence from the PASAv1 transcriptome assembly, as well as the TrinDNv2 Trinity *de novo* assembly; (ii) homology to SwissProt mammalian proteins in UniProtKB and the predicted proteomes of *Cebus capucinus imitator* (NCBI release 1.0), *Saimiri boliviensis* (Ensembl, SaiBol1.0), *Callithrix jacchus* (NCBI release 3.2), *Aotus nancymaae* (NCBI release 2.0), and human (GRCh38.p7); and (iii) *ab initio* predictions from Augustus v3.3 (*90*) using the robust capuchin-specific HMM that we trained in BUSCO (Table S6). We allowed direct prediction from aligned protein and transcript evidence (est2genome and protein2genome), and specified "primates" as the organism for the RepeatMasker library. We also specified the following settings, with other options as default; max_dna_len=500000, min_contig=1000, split_hit=100000, min_intron=20, single_exon=1, single_length=250, and keep_preds=1.

Following the first pass of Maker, we filtered the predicted gene models based on a maximum AED (Annotation Edit Distance) of 0.25 and a minimum length of 50 amino acids (aa) to generate a high-quality set of strongly supported gene models. We used the autoAug.pl script within Augustus to retrain the robust capuchin-specific HMM using the high-quality gene models from the first pass of Maker as the training set, as well as evidence from the comprehensive transcriptome generated in PASA (*106*) (PASAv1), with three rounds of optimisation. The setup for the second pass of Maker was identical except we used the capuchin-specific HMM that was retrained in Augustus, and we no longer allowed direct prediction from aligned protein and transcript evidence (est2genome and protein2genome).

After the second pass in Maker, there was a notable increase in the number of predicted gene models owing to an inflated number of short single exon predictions with high AED scores seemingly driven primarily by aligned RNAseq evidence from the Trinity and PASA transcriptomes. This appeared to be a result of noise and spurious alignments from the deeply sequenced transcript assemblies. One proposed solution was to turn off single exon predictions in Maker (*19*); however, we opted instead to further process our transcriptome to generate a refined high-quality non-redundant transcript assembly (NRv1) as described in a previous section. Additionally, in our downstream pipeline to assess signatures of positive selection, we pull the longest isoform for each gene for other primates. As we are only annotating a single isoform per gene in Maker pipeline, this NRv1 transcript assembly pushed Maker to predict the longest transcript per gene that had good evidence.

Following the second pass of Maker, we again filtered the predicted gene models, this time based on a stricter maximum AED of 0.10 and a minimum length of 70 amino acids to generate a set of strongly supported gene models. We employed this set of gene models to again retrain the robust capuchin-specific HMM using the autoAug.pl script within Augustus along with evidence from NRv1 transcript assembly and another three rounds of optimisation. In the third and final pass of Maker, we incorporated: (i) direct evidence from the NRv1 transcript assembly; (ii) homology to the predicted proteomes of *Cebus capucinus imitator, Callithrix jacchus*, and human (releases as above), and *Saimiri boliviensis* but with NCBI release 1.0 rather than Ensembl as in previous rounds; and (iii) *ab initio* predictions from Augustus using our final retrained robust capuchin-specific HMM (Table S6). We also specified the following settings for the third pass, with other options as default; max_dna_len=350000, min_contig=1000, min_protein=30, split_hit=100000, min_intron=20, single_exon=1, single_length=250, and keep_preds=1.

We allowed Maker to retain *ab initio* predictions that had no supporting evidence (keep_preds=1) which were assigned the highest AED of 1. We then searched our set of predicted gene models from the third pass of Maker for Pfam domains using InterProScan 5 (*113*). We removed all gene models that had an AED of 1 except for those with a Pfam domain from our final set of gene model predictions allowing the retention of novel gene models that have an AED of 1 but are likely true genes owing to the existence of known protein domains. After also removing non-coding tRNA predictions, we retained 26,592 gene model predictions in our Maker gene set.

### 1.8 Functional annotation

We obtained annotated motifs and domains in available databases (Pfam, PANTHER, ProDom, PRINTS, SMART, and PROSITE) for our gene models with InterProScan 5 (*113*). Gene functions were assigned according to the best match for each gene in alignment to the NCBI non-redundant (nr) protein database using Blast2GO v5.2.5 (*91*). We first downloaded a pre-formatted nr protein database and a GI list for all vertebrate proteins from Entrez (NCBI), and then modified the database to include only proteins in that list. We performed a BLASTP search against this vertebrate nr protein database using our set of final Maker gene predictions as the query with the options "-outfmt 14 -evalue .0001 -word_size 3 -show_gis -num_alignments 20 -max_hsps 20." Gene ontology analysis was performed with Blast2GO using default settings.

Of the 26,592 gene models in our final Maker gene set, 3,313 gene models were not annotated through our BLASTP search or with GO terms in Blast2GO. Many of the gene models lacking functional annotation, however, appeared to be uncharacterised proteins that had some BLASTP hits and/or known protein domains. The remaining 1,313 gene models that had no BLASTP hit and no annotated motifs or domains were removed from our gene model set. Our final set of gene models thus contained 25,279 predicted genes. We used Blast2GO in combination with our reference genome and updated GFF annotation file to generate FASTA files containing mRNA (transcript), CDS, and protein sequences for this final set of annotated gene models.

We note that Dovetail's HiRise scaffolding using the Chicago libraries increases contiguity but also adds a series of 100 N bases where joins are made resulting in long scaffolds containing numerous gaps. For example, our final draft genome assembly contains 118,679 more bases as gaps then the preliminary genome assembly input into HiRise for scaffolding. These gaps are disruptive to gene prediction with Maker when they occur within a gene leading to the fragmentation of some gene models as has been found in other studies (e.g., (*114*)). This gene model fragmentation likely partially explains the slightly inflated number of predicted models in our final gene set in comparison to the number of protein coding genes annotated by NCBI and Ensembl for other platyrrhine primate genomes.

### 1.9 Mitochondrial genome assembly

We generated a set of putative mitochondrial read pairs by mapping the short reads retained after screening with the Kraken2 database to a complete *Sapajus apella* mitochondrial genome (GenBank accession JN380205.1) using bbmap.sh (default settings) from bbmap tools v.37.99

(https://sourceforge.net/projects/bbmap). Read pairs (N = 795,689) that mapped to the reference were then used to assemble the mitochondrial genome for our reference individual, Mango, in a two-step procedure. In the first step, we prepared an initial assembly by mapping these read pairs to the same complete *S. apella* mitochondrial genome from GenBank using MIRA v.4.0.2 (*86*). We then performed baiting and iterative mapping using the MITObim v.1.9.1 (*87*) wrapper script to generate the final mitochondrial genome assembly for our reference individual, which was subsequently eye checked and manually annotated in Geneious R7.1 (Biomatters). The final assembly is 16,550 bp in length and was included as the final FASTA sequence in the version of the genome assembly (Sape_Mango_1.1) uploaded to NCBI under the accession JAGHVQ. The mitochondrial genome assembly is also available as a standalone FASTA along with the annotation file on a Zenodo repository (see data availability).

### 2) Extended methods & results: Signatures of selection in platyrrhine genomes

#### 2.1 Identification of orthologs

In order to assess signatures of positive selection in robust capuchin and other platyrrhine primates genomes, we first identified one-to-one orthologs across ten species. Our choice of species was partially influenced by the availability of Ensembl genome annotations. We downloaded predicted CDS and protein sequence files from Ensembl for the gracile capuchin (*Cebus imitator*; v.1.0), squirrel monkey (*Saimiri boliviensis*; v.1.0), marmoset (*Callithrix jacchus*; ASM275486v1), rhesus macaque (*Macaca mulatta*; v.8.0.1), chimpanzee (*Pan troglodytes*; v.3.0), human (*Homo sapiens*; GRCh38), tarsier (*Carlito syrichta*; v.2.0.1), mouse lemur (*Microcebus murinus*; v.3.0), and mouse (*Mus musculus*; GRCm38). For the robust capuchin (*Sapajus apella*), we included the final set of annotated *de novo* gene models from our Maker pipeline.

The protein and CDS sequence files from Ensembl were formatted as follows: (i) we reduced redundancy in the form of different isoforms for the same gene by retaining only the isoform with the longest protein sequence; (ii) we then modified the FASTA sequence headers to include only an organism ID, which was a unique four letter code derived from the first letter of the genus name and first three letters of the species name (Cimi, Sbol, Cjac, Mmul, Ptro, Hsap, Csyr, Mmur, Mmus), and sequence ID, which was the numeric portion of the ID given to each gene by Ensembl (e.g., >Cimi 00000000672.1), such that corresponding protein and CDS sequences had identical headers; and (iii) we removed sequences for proteins that were shorter than 30 amino acids (aa) in length. Our robust capuchin CDS and protein files did not require reformatting as we predicted only one isoform per gene, specified a minimum protein length of 30 aa in our annotation pipeline, and the FASTA headers were in the above format with Sape as the organism ID followed by a numeric sequence ID.

To identify orthologs across our proteins sets, we employed a custom configuration of OrthoMCL v.2.0.9 (https://github.com/apetkau/orthomclsoftware-custom) used within the OrthoMCL pipeline (*92*) with our reformatted FASTA sequence files for each species as input. We found 9,342 orthologs in all ten species, 5,652 of which were one-to-one orthologs, and 12,160 one-to-one orthologs that were recovered in at least two of our species. Ortholog IDs were assigned based on the output from OrthoMCL (referred to as group ID). For each species, the number of CDS/protein sequences input into OrthoMCL, orthologs, and one-to-one orthologs (found in at least two species) are shown in Table S7. We collected information for all one-to-one orthologs and generated protein and CDS FASTA files for each using OrthoMCL Tools v1.0 (https://github.com/guyleonard/orthomcl_tools).

These initial groups of ortholog sequences derived using OrthoMCL were, as described in the following sections, then filtered and processed into multi-species codon-based alignments, forming the basis of codon models to test for positive selection and are assigned to genes with symbols and IDs. These groups are referred to as orthologs, one-to-one orthologs, alignments, models, and genes depending on the stage of processing and relevance, however, they retain the same group ID as they progress through the pipeline and are sometimes called groups regardless of pipeline stage.

#### 2.2 Ortholog alignment and quality checks

Given our focus on capuchins in this study, we removed orthologs for which both the robust and gracile capuchin were missing (N = 1,229), retaining 10,931 one-to-one orthologs. We then sorted and filtered the one-to-one orthologs based the number of species each was recovered in, requiring a minimum of five species to be retained (N = 9,911). We aligned the CDS nucleotide sequences for these 9,911 groups by codon using Guidance2 v.2.02 (*20*) with the MAFFT aligner v.7.419 (*93*). We ran Guidance2 with 100 bootstraps, which allows the assignment of confidence scores to aligned sequences, columns, and residues. Unreliable columns (as codons) that do not align the same way greater than 93% of the time and unreliable sequences with alignment confidence scores < 0.6, as well as alignments with an incorrect number of nucleotides (i.e., not a multiple of three) or an internal stop codon, were then removed. We modified the sequence headers in the aligned

FASTA files to remove the numeric portion of the sequence ID corresponding to the Ensembl (or Sape) gene ID and keeping only the organism ID as the header for each species. The aligned FASTA files were then reformatted into PHYLIP alignments as required for downstream analyses.

An initial test run of the branch-site model (BSM) test with codeml from the PAML package v.4.9 (*21*) recovered a seemingly high proportion of genes with positively selected sites along each foreground branch. Through eye-checking some alignments in Geneious R7.1 (Biomatters), we found many groups with significant signatures of selection that were possibly false positives owing to apparent errors. Many studies have highlighted the tendency for positive selection tests, which are highly sensitive, to detect false positives due to primary sequencing, assembly, annotation, and alignment errors, such that not all columns in the alignment represent homologous protein-coding positions (*114–120*). Guidance was effective at minimising alignment errors and most of the spurious alignments appeared to be assembly and annotation artifacts including issues with predicted gene models, exon boundaries, pseudogenised genes, and the recovery of groups containing similar paralogous loci rather than orthologs, as well as sequencing and assembly errors.

As a result, to reduce the likelihood of false positives and true negatives, we decided to visually inspect all alignments and manually edit them in Geneious R7.1. We conservatively trimmed or masked dubious and unreliable regions, or excluded entire dubious sequences, as necessary, before running our positive selection analyses. Guidance cannot assign confidence scores to columns where there is only one sequence (and therefore doesn't remove them), thus these regions were removed while editing in Geneious. We also removed columns (by codon) that were missing in several sequences (depending on the number of species in the alignment), removed entire species/sequences that were missing from a significant portion of the alignment, and discarded entire alignments when the overall quality seemed poor or they became very short. We assigned an Entrez ID (DAVID's preferred ID) and gene symbol to each group based primarily upon the Ensembl gene ID, symbol, and description annotation for the human sequence in the alignment. If the human was missing, we used the Ensembl information for the gracile capuchin, marmoset, or squirrel monkey. We subsequently also removed groups (N = 364) that were missing the squirrel monkey and one capuchin because the requirements we defined for assessing each target lineage meant these alignments would not be analysed (see next section). Overall, this resulted in a set of 9,216 conservative, manually-curated CDS alignments which were highly likely to represent one-to-one orthologs across their length that were used in downstream analyses. Information on each of the 9,216 alignments, including group ID, assigned gene symbol and Entrez ID, and Ensembl (or Sape) gene ID for the original sequences for each species, can be found in Table S8. The platyrrhine species are found in the following number of these final alignments; *Sapajus* in 7,134, *Cebus* in 9,092, *Saimiri* in 9,003, and *Callithrix* in 8,921 (see Table S7 for ortholog and other counts for each species).

Species set IDs were generated for each combination of species found in the final alignments, with the first number in those species set IDs denoting the number of species in the set and numbers from after the first underscore represent missing species as follows *Sapajus* (Sape, _1), *Cebus* (Cimi, _2), *Saimiri* (Sbol, _3), *Callithrix* (Cjac, _4), *Macaca* (Mmul, _5), *Pan* (Ptro, _6), *Homo* (Hsap, _7), *Carlito* (Csyr, _8), *Microcebus* (Mmur, _9), *Mus* (Mmus, _10) (Table S7). An alignment with 8 species with Mmul (_5) and Csyr (_8) missing, for example, would be assigned to species set 8_5_8. The set with all ten species is referred to as the "full" set.

In total, there were 207 different combinations of species represented in the final alignments/groups (N = 9,216); one set with all species (full; N = 4,636), 10 sets of 9 species (N = 2,819), 41 sets of 8 species (N = 1,083), 61 sets of 7 species (N = 443), 59 sets of 6 species (N = 149), and 35 sets of 5 species (N = 86). Detailed information regarding these species sets including counts of groups for each species set, and counts of species sets per lineage analysed and per species can be found in Table S9.

#### 2.3 Branch model and branch site model tests for positive selection

We specified six target lineages (foreground branches) as follows: (H1) *Sapajus*; (H2) *Cebus*; (H3) Cebinae ancestor; (H3a) across-capuchins (all Cebinae; branches H1, H2, and H3 combined); (H4) Cebidae ancestor (i.e., ancestor to capuchins and squirrel monkeys); and (H5) *Saimiri* (Figure 1). Note, we consider Cebidae to be comprised of squirrel monkeys and capuchins with callitrichids in their own family (Callitrichidae). For H1, H2, and H3, both *Sapajus* and *Cebus* were required in the alignments. For H3, we accepted alignments without *Saimiri* given the much greater length of the ancestral Cebinae branch versus the ancestral Cebidae branch between the Cebinae/Saimirinae divergence and Callitrichidae, though these were a very small proportion of the total. For H3a, we accepted alignments with just one capuchin lineage, and required *Saimiri*; we aimed to uncover signatures generally selected for capuchins and that would be more comparable to studies analysing a single capuchin lineage. For H4, we required *Saimiri* and

*Callithrix* for this branch. Because all alignments had at least one capuchin, further requirements for H5 were simply the presence of *Saimiri*. Information on the lineages analysed for each group and each species set can be found in Tables S8 and S9, as well as group counts per lineage in Table S10.

We generated unrooted tree files, as required by codeml (*21*), for the full set of ten species, and each possible combination of species in the alignments including between five and nine species. In total, there were 207 different combinations of species (species sets) and thus, possible unrooted trees. The guide tree topologies were based on the well-accepted consensus primate phylogeny for the species included (Figure 1) (*1*, *121*) with the bifurification at the basal node removed. The unrooted tree in newick format for the full species set is ((((((Sape,Cimi),Sbol),Cjac),(Mmul,(Ptro,Hsap))),Csyr),Mmur,Mmus). See Tables S8 and S9 for rooted/unrooted newick trees per group and per species set, respectively. The specification of the various foreground branches (which are denoted with #1) for the lineages analysed according to the above rules resulted in 759 tree files. For each alignment, branch lengths were calculated from the data by codeml.

We ran two different tests for positive selection with codeml from the PAML package v.4.9 (*21*) which are based on rates of non-synonymous versus synonymous substitutions ($\omega$ or dN/dS ratio): the branch-site model (BSM), which tests for positively selected sites within the alignment in each target (foreground) lineage; and the branch model (BM), which tests for elevated (accelerated) or decreased (decelerated) dN/dS ratios across the alignment along the target branch versus the background rate along the other branches in the tree. For the across-capuchins lineage (H3a), we only performed the BM test as it was difficult to interpret the BSM test (which is for episodic selection) results when ran across multiple branches together. As such, a total of 11 lineage and test combinations were conducted.

For each branch model (BM) test, we assessed two models as follows; the alternative branch model which separates the tree into foreground and background branches which have distinct $\omega$ parameters ($\omega_0$, $\omega_1$) allowing them to evolve with separate dN/dS ratios; and the null model, which uses a single $\omega$ parameter for the entire tree. For each BSM test, we assessed an alternative branch-site model allowing for positive selection on the foreground branch and a null model allowing only for purifying and neutral selection on the foreground and background lineages. To achieve convergence, we carried out three replicates with different starting values for omega (0.2, 0.7, 1.2) or kappa (0.2, 2, 5) for the alternative model in each BM or BSM test, respectively, running four analyses per model test. After estimating the parameters and calculating the likelihood with codeml, we performed likelihood ratio tests (LRTs) by comparing the likelihood of the alignment under the alternative model (using the maximum of the log-likelihood scores across the three replicates) versus under the null. For the alignments that did not converge with the various starting omega or kappa values (i.e., the alternative model had a lower likelihood than the null model in all replicates, which occurred more frequently in BSM tests), we manually changed the LRT statistic of those tests to zero. We calculated p-values from the chi-square distribution with one degree of freedom. For both BM and BSM, we employed the LMAP package v.1.0.2 (*122*) to handle some aspects of the codeml workflow including initial directory organisation, generation of control files, task execution, and extraction of the likelihood estimates.

In total, considering alternative and null models, the three replicate starting values for omega or kappa for the alternative models, and the six or five different foreground branches for BM and BSM tests, respectively, we ran a total of 345,940 codeml analyses to test 86,485 models, 47,744 for BM and 38,741 for BSM (Table S10). Groups analysed per lineage varied between 6,978 and 9,003 of 9,216 total groups (Table S10), with averages of 7,957 BM and 7,748 BSM tests.

For the BM tests, we used the maximum likelihood estimate of the two $\omega$ parameters to identify genes where the estimated foreground $\omega$ ($\omega_1$) was higher (accelerated) or lower (decelerated) than the background ($\omega_0$), and then identified significantly accelerated and decelerated genes based on p-values (< 0.05). We generated "signed" LRT statistics such that accelerated genes were assigned a positive value and decelerated genes were assigned a negative value; these decelerated genes were not considered further in this study. A highly positive signed LRT score represents strong evidence for a lineage-specific elevated dN/dS ratio, which could be explained by the effects of both positive selection or relaxed constraint (*123, 124*). Alignments with the significant signatures of accelerated evolution in BM tests, however, are good candidates for adaptively evolving genes in the foreground branch of interest. The BSM specifies evidence for episodic positive selection by allowing $\omega$ to vary among sites and lineages, enabling detection of selection in a subset of sites within specific branches (*118*). Although BSM tests may be considered to be more directly indicative of positive selection, they are also more parameter rich than BM tests (with two more parameters) and parameter estimates may be noisier when sequences in an alignment are very similar (e.g., for the capuchin lineages). Indeed, previous studies have provided support for the limited power in BSM tests to detect strong signatures of positive selection across closely related lineages (e.g., *125*, *126*). In line with this, across the six lineages analysed, we recovered 248 to 552 (avg. 351) models (BM) with significant signatures of accelerated evolution, and 75 to 186 (avg. 113) models (BSM) with significant signatures of episodic positive selection, much fewer than for BM tests and particularly for the shorter branches. Between 17 and 34 (avg. 25) groups are significant for both BM and BSM tests for the same lineage (Table S10).

We used the Benjamini-Hochberg method (*127*) to correct for multiple testing within each foreground branch for the BSM and BM tests by controlling the false discovery rate (FDR). There were no or very few (≤ 5) significant genes after FDR correction for all BSM tests, and most lineages for BM (except for across-capuchins (H3a) and squirrel monkeys (H5)). Most foreground lineages defined in our analyses represent short branches in the phylogeny for which the power of these tests to detect selection is reduced (especially BSM tests) leading to lower LRT scores (see Table S10). High LRT statistics, however, are a cause for concern as they are often indicative of inflated dN/dS ratios that result from errors with the predictions or alignments (e.g., the issue addressed by manually editing and removing errors from the alignments). A low number of genes with high LRT scores is biologically realistic for shorter branches given the limited timeframe for mutations to accumulate. We here consider the significant FDR-corrected genes to be very strong candidates for adaptively evolving genes and the overall set of significant genes according to uncorrected p-values to be the most likely set of genome-wide candidates from our background set. Our gene set enrichment analyses (discussed below) are conducted on the set of significant genes according to uncorrected p-values for each lineage. For each lineage and test (BM/BSM) combination, we also rank each significant gene based on p-value with the gene with the smallest p-value being ranked first. While we focus primarily on the results of our gene set enrichment analyses, rather than individual genes, the ranks allow assessment of the strength of our results when focusing on specific genes of interest.

Lists of all groups (genes) analysed for each of the six lineages, along with significance for BM and/or BSM tests, can be found in Tables S11 (H1), S12 (H2), S13 (H3), S14 (H3a), S15 (H4), and S16 (H5). More detailed information for the groups with significant evidence of accelerated evolution or episodic selection from the BM and/or BSM tests including p-value, LRT statistic, likelihood scores, and FDR significance, is located in Tables S17 (H1), S18 (H2), S19 (H3), S20 (H3a), S21 (H4), and S22 (H5).

2.4 Gene set enrichment analysis

We conducted 11 gene set enrichment analyses, one for the set of significant genes from each combination of lineage and test (BM or BSM) using DAVID v.6.8 (*22*). For each enrichment analysis, we used the Entrez ID for each group in the gene set and the entire human gene set as the background population of genes. All lineages had much fewer (~1/2 to 1/3) total groups/genes analysed (between 6,978 and 9,003) than the human gene set used as the background, thereby reducing the power of the gene set enrichment analyses, and we primarily consider the enriched GO terms and other annotations based on DAVID's enrichment p-values (EASE scores) (< 0.05).

In DAVID, we assessed functional annotation clustering across BP (biological process), CC (cellular component), and MF (molecular function) gene ontology (GO) terms (the "all" option) together under the high classification stringency criteria with an EASE score of < 0.05 required for each enriched GO term in the clusters. All clusters recovered under these criteria had an enrichment score of greater than 1.3, which is a statistical support metric for clusters corresponding to the geometric mean of all EASE scores for each annotation term in the cluster and equivalent to non-log scale 0.05 significance score (*22*). We also assessed lists/charts of the enriched (i) BP, CC, and MF GO terms, (ii) UniProt (UP) keywords, (iii) KEGG pathways, (iv) Reactome pathways, and (v) disease annotations, with a minimum EASE score of 0.05 for all annotations. A workflow summary graphic for the ortholog identification, alignment, codeml, and gene set enrichment analysis steps is shown in Figure S3.

Across the BM gene sets for the six lineages, between 2 and 13 (avg. 6) GO clusters, and 68 and 189 (avg. 103) enriched terms (all annotation categories) are recovered. Across the BSM gene sets for the five lineages, between 0 and 9 (avg. 3) GO clusters, and 10 to 123 (avg. 60) enriched terms (all annotation categories) are recovered. Counts of GO clusters and all enriched annotation categories for each gene set enrichment analysis are listed in Table S23.

Lists of all enriched annotated terms and GO clusters including annotation category, term description, term ID, gene counts and hits, and statistical results (e.g., EASE score and fold enrichment) for each gene set enrichment analysis are found in Tables S24–27 (H1), S28–30 (H2), S31–34 (H3), S35–36 (H3a), S37–40 (H4), and S41–44 (H5) (see also the guide in Table S23).

Extended results from the gene set enrichment analyses for each lineage are presented below in the supplement.

All gene function information contained throughout the supplemental sections and the main text discussion was retrieved from the GeneCards (*128*) page for that gene (https://www.genecards.org/cgi-bin/carddisp.pl?gene=XXX, with XXX being the gene symbol), accessed between Nov-2020 and Feb-2021, unless otherwise cited. Further research was carried out into genes of particular interest.

### 3) Extended results: Robust capuchins (*Sapajus*; H1) positive selection results

Lists of all enriched annotated terms and GO clusters including annotation category, term description and ID, gene counts and hits, and statistical results such as EASE score and fold enrichment, for the BM and BSM gene set enrichment analyses for robust capuchins (*Sapajus*: H1) are found in Tables S24–27.

#### 3.1 Face and skeletal system development

For robust capuchins (*Sapajus*), the BM gene set is enriched for genes related to face morphogenesis and skeletal system development. The top annotation cluster in DAVID for the BM gene set with an enrichment score of 2.56 contains four BP GO terms, which are also among the top ranked individual GO terms: "face morphogenesis" (top term, 11.1 fold enrichment), "head morphogenesis" ($2^{nd}$ term, 9.5 fold enrichment), "face development" ($6^{th}$ term, 6.9 fold enrichment) and "body morphogenesis" ($7^{th}$ term, 6.9 fold enrichment). These four GO terms include the same five genes in this gene set: *SGPL1*, *RRAS*, *NIPBL*, *CSRNP1*, *TIPARP*.

Delangin, the protein encoded by *NIPBL*, plays a critical role in the regulation of the cohesion complex which functions in sister chromatid cohesion and transcriptional regulation of genes that are essential for normal development, growth, and patterning, and is found prenatally in the developing limbs and bones of the skull and face. Mutations or other defects in this gene are the primary cause of Cornelia de Lange syndrome, a multisystem disorder characterised by distinctive facial features and skeletal dysmorphology, developmental delay/intellectual disability, slow postnatal growth, limb malformations, and hirsutism, among other anomalies (*76*). *CSRNP1* (*AXUD1*) encodes a transcription factor that plays a central role in mediating neural crest cell development as a downstream effector of Wnt signalling (*129*). The neural crest is a progenitor cell population that gives rise to craniofacial cartilage and bones, peripheral and enteric neurons and glia, and melanocytes. *RRAS* encodes a small GTPase involved in diverse processes controlling cell adhesion, migration, and proliferation, and plays an important role in development. Mutations in this gene cause dysregulation of RAS signalling by disrupting signal flow in the MAPK/ERK cascade and underlie a Noonan syndrome-like disorder within the RASopathy family (*130*). The RASopathy family of disorders are characterised by facial dysmorphism, cognitive deficits, cardiac defects, defective postnatal growth, and skeletal and ectoderm anomalies. *TIPARP* protects the pluripotency of embryonic stem cells and plays a role in transcription factor regulation, among other functions. It is highly expressed in the brain with recent studies on knockout mice indicating it is required for correct development of the cortex (*131*). *SGPL1*, which is found in both gene sets, encodes an endoplasmic reticulum enzyme with a central role in sphingolipid catabolism, catalysing the irreversible degradation of sphingosine-1-phosphate (S1P) and other phosphorylated long-chain bases. S1P is a secreted bioactive signalling molecule that critically regulates many physiological and immune related processes, and is particularly important in the development of the vascular system and central nervous system (CNS) (*132*).

Among the other most significantly enriched GO terms according to EASE score in the BM gene set are several related to the skeletal system including "skeletal system development" and "skeletal system morphogenesis" (16 and 9 genes, 2.2 and 2.8 fold enrichment), with four genes also found in the face morphogenesis cluster noted above. Other genes include *RAB23*, *SIX4*, *WNT10B*, *GDF5*, *DYM*, *NLE1*, and *PDGFC*. *RAB23* encodes another small GTPase of the RAS superfamily that plays essential roles in embryogenesis as an upstream negative regulator in important signalling cascades including the sonic hedgehog (Shh) pathway (as an inhibitor of the transcription factor *GLI1*) and fibroblast growth factor (FGF) pathway. It plays a well-established role in heart and limb patterning, neural tube closure, and skeletal development, and recently has been shown to coordinate early osteogenesis, showing activity in osteoblasts, and controlling the growth and fusion of skull bones in developing animals (*75*). Premature fusion of multiple skull sutures owing to elevated osteogenesis is seen in Rab23-deficient mice, which agrees with the implication of *RAB23* mutations in patients with Carpenter syndrome, a developmental disorder characterised by craniosynostosis and polysyndactyly, among other symptoms (*75*). Craniosynostosis is the premature fusion of craniofacial sutures causing major disruptions to growth of the face and skull. *SIX4* encodes a sine oculis homeobox (SIX) transcriptional regulator targeting processes like cell differentiation, migration, and survival, and plays an important role in embryonic development, notably in cranial morphogenesis and synchondrosis development. Synchondroses formed by endochondral ossification in the cranial base are an important neurocranial growth centre with abnormalities impacting cranial base elongation and the development of the craniofacial bones (*74*, *133*).

*WNT10B* of the Wnt ligand gene family encodes a secreted protein that specifically activates canonical Wnt/β-catenin signalling. It is well established to be involved in the control of stemness, pluripotency, and cell fate decisions, particularly in bone, as well as the immune system, mammary gland, skin, and adipose tissue (*134*). *WNT10B* is implicated in osteoporosis and breast cancer, among other disorders and associated with abnormal jaw, dental, and digit morphologies (HPO). *GDF5* encodes a secreted growth/differentiation factor of transforming growth factor-beta (TGFB) family that is essential for normal skeletal and cartilage development, with an important role in joint formation, maintenance, and remodelling/repair, and is a major risk locus for osteoarthritis (*135*, *136*). *DYM* encodes a protein which is essential to endochondral bone formation during early development and is expressed throughout the entire growth process of embryonic and foetal tissues. Both *DYM* and *GDF5* are implicated in various skeletal dysplasias, and associated with a large number of bone-related human phenotypes (HPO). *NLE1*, notchless, encodes a regulator of Notch activity, as well members of the Wnt pathway, and it is required during embryogenesis for inner mass cell survival and formation of the axial skeleton (*137*). *PDGFC* encodes a platelet-derived growth factor (PDGF) required for normal skeletal formation during embryonic development, particularly the craniofacial skeleton, palate, and CNS (*138*). Other genes in these GO terms include the homeobox gene *HOXA6* (along with another homeobox gene not in these terms, *HOXD1*), the cartilage-specific lectin *CLEC3A*, and *COL19A1*, which encodes the alpha chain of type XIX collagen.

Although not annotated with these enriched GO terms, *DKK3* is a related gene found in the BM and BSM gene sets. DKKs are central to vertebrate development, locally inhibiting Wnt-regulated processes such as limb development, and are associated with bone formation and disease in adults (*139*). Other interesting genes in the BSM gene set include another growth differentiation factor, *GDF2*, which is a potent inhibitor of angiogenesis, as well as regulates cartilage and bone development, and differentiation of cholinergic neurons. Similarly, other interesting genes in the BM gene set include *STMN2*, thought to be involved in osteogenesis; *TMEM57*, associated with acrofacial dysostosis which is characterised by distinctive craniofacial malformations; *WISP2*, which encodes a member of the connective tissue growth factor family with an important role in modulating bone turnover and promoting the adhesion of osteoblast cells; and *FBLN7*, which is involved in tooth development and the differentiation/maintenance of odontoblasts in dentin formation.

A related enriched GO term that shares all genes with some aforementioned terms include "platelet-derived growth factor receptor signalling pathway" with the genes *TIPARP*, *PDGFC*, *SGPL1*, and *CSRNP1* (5.6 fold enrichment). PDGF receptor signalling plays a crucial role in specifying mesenchymal stem cell commitment to mesenchymal lineages such as osteoblasts, chondrocytes, myocytes, and adipocytes (bone, cartilage, muscle, and fat cells, respectively). Furthermore, the BM gene set also contains the enriched BP GO term "regulation of bone morphogenetic protein signalling pathway" with 5 genes (4 fold enrichment). In addition to *GDF5* (discussed above), these genes include *SMAD4*, which encodes a Smad signal transduction protein that is a crucial component of the bone morphogenetic protein (BMP) signalling pathway; *FBXL15*, which acts as a positive regulator of the BMP signalling, and is required for bone mass maintenance and for dorsal/ventral pattern formation; *HFE2*, which acts as a BMP coreceptor and regulates iron homeostasis; and *SKOR2*, which plays an essential role in development of the cerebellum and represses BMP signalling.

Other related enriched BP GO terms for the BM gene set include "skeletal muscle tissue (organ) development" (8 genes, ~3 fold enrichment). Unsurprisingly, there is some overlap with the skeletal system terms with three shared genes: *SIX4*, *WNT10B*, and *COL19A1*. *SIX4* has multiple important functions at various stages of muscle development. Other genes in these GO terms include *ANKRD2*, which encodes a muscle ankyrin repeat protein that functions as a negative regulator of myocyte differentiation, regulating gene expression during muscle development and in response to muscle stress; *USP2*, a ubiquitin-specific protease that plays a role in the regulation of myogenic differentiation of embryonic muscle cells; *MYF6*, a myogenic factor involved in muscle differentiation, inducing fibroblasts to differentiate into myoblasts; and *MYORG*, which promotes myogenesis. Other related GO terms include "muscle tissue development" and "striated muscle tissue development" (12 and 11 genes, ~2.1 fold enrichment), and "regulation of skeletal muscle tissue development" (4 genes, 5.2 fold enrichment).

#### 3.2 Other development and morphology related

The third cluster for the BM gene set (enrichment score 1.7) is comprised of four enriched BP GO terms related to the female development; "female gonad development", "development of primary female sexual characteristics", "female sex differentiation", and "ovulation cycle process" (5 to 6 genes, 3.7 to 4.4 fold enrichment). Interesting genes specific to the female development GO terms include *NRIP1*, involved in the regulation of ovarian function and modulates transcriptional activity of the oestrogen receptor; and *FSHB*, which encodes the beta subunit of the follicle-stimulating hormone (FSH) involved in follicle development and spermatogenesis. Related genes in the BM gene set include *PTX3*, which is implicated in female fertility, and *PANX1*, which plays a critical role in oogenesis; and in the BSM gene set include *HSD17B12*, which catalyses the conversion of oestrone into oestradiol in ovarian tissue, and may play a role in oestrogen formation; and *PRLH*, which encodes prolactin releasing hormone that stimulates prolactin release and regulates the expression of prolactin. There is also the enriched GO term “regulation of hair cycle” (3 genes, 8.7 fold enrichment) in the BM gene set.

The second annotation cluster for the BM gene set according to DAVID’s enrichment score (2.5) is comprised of kidney, urogenital system, and renal system development BP GO terms (12 to 13 genes, 2.7 to 3 fold enrichment). There are also some broad GO annotations related to these terms including “system development” (78 genes, 1.2 fold enrichment), “regulation of localisation” (49 genes, 1.3 fold enrichment), and “organ morphogenesis” (25 genes, 1.7 fold enrichment). For the BSM gene set, there are multiple enriched GO terms related to the endothelium including the top two BP terms “negative regulation of blood vessel endothelial cell migration” and “negative regulation of endothelial cell proliferation” (3 genes, 29.6 and 23.2 fold enrichment), as well as several other highly similar terms with the same angiogenesis inhibiting genes (*APOE*, *GDF2*, *MMRN2*).

#### 3.3 Cilia

The fifth BM cluster according to DAVID’s enrichment score (1.66) is comprised of four GO terms related to cilia; “cilium movement”, “regulation of cilium beat frequency”, “regulation of cilium movement”, and “regulation of microtubule-based movement” (5.2 to 18.1 fold enrichment). This cluster of GO terms annotates four genes in the BM set: *DNAAF1*, *BBS2*, *CFAP20*, and *TTLL1*. *BBS2* is part of the BBSome complex which is required for ciliogenesis and an associated disease in humans is Bardet-Biedl Syndrome (type 2), which causes progressive visual impairment due to cone-rod dystrophy, polydactyly, hypogonadism, kidney abnormalities, and learning difficulties. The protein encoded by *DNAAF1* is cilium-specific, required for the stability of the ciliary architecture, and involved in the regulation of microtubule-based cilia. *CFAP20* encodes another cilium-specific protein that plays a role in axonemal structure organisation and motility, and in the regulation of cilium size and morphology. *TTLL1* plays a role in post-translational modifications of tubulin, specifically in polyglutamylation, which functions in regulating motile cilia. Loss of function or absence of the *TTLL1* gene in mice is associated with reduced motility of respiratory cilia, and infertility owing to sperm with truncated axonemes. The terms in this cluster relate more specifically to motile cilia which are found in sperm cells and in epithelial cells lining the oviduct (to move the ova to the uterus), the airways (to clear mucous), and the brain ventricles where they provide planar polarity essential for cerebrospinal fluid (CSF) circulation, with defects in motile cilia resulting in an excessive accumulation of CSF causing congenital hydrocephalus and severe neurological damage (*140*).

Among the most enriched CC GO terms is "MKS complex", a protein complex that organises the inner structure (Y-shaped links) of the ciliary transition zone. This GO term includes three genes in the BM gene set, *AHI1*, *CEP290*, and *TMEM67*. Each of these genes are involved in ciliogenesis and are required for the formation of primary non-motile cilium. *AHI1* is required for ciliogenesis and both cerebellar and cortical development in humans, and may play a crucial role in ciliary signalling during cerebellum embryonic development as a positive modulator of classical Wnt signalling (*35*). *CEP290* is the most frequently mutated gene in ciliopathies and implicated in specific forms of Joubert syndrome related disorders (*141*). Related enriched UP keywords in the BM gene set include "cilium" (8 genes, 2.7 fold enrichment) and "Bardet-Biedl syndrome" (3 genes, 9.3 fold enrichment), with "Joubert syndrome" falling just above the EASE score threshold (0.055).

#### 3.4 Neurodegeneration and the nervous system

There are several important brain-related genes in the BM and BSM gene sets, in particular with a relevance to Alzheimer's disease (AD), including genes found in both gene sets such as *ITPKB*, which has high expression in the brain and is upregulated in AD, *HVCN1*, and *PROX2*, which is associated with longevity in humans and with neuron differentiation. Interesting genes in the BSM gene set include *RNF103*, which is highly expressed in the cerebellum; *APOE*, which is associated with AD and many related human phenotypes (HPO) including cognitive and memory impairment, neurofibrillary tangles, and senile plaques; and *NCAN*, neurocan, an important chondroitin sulphate proteoglycan (CSPG) found in the extracellular matrix (ECM) of the brain which may modulate neurite growth during development. For the BSM gene set, enriched disease annotations include various interesting brain and neurological related terms including some related to Alzheimer's disease and dementia, and "amyotrophic lateral sclerosis" (ALS). The sole enriched UP keyword for the BSM gene set is "neuropathy" (8.6 fold enrichment) with three genes; *SNAP29*, which binds syntaxins and mediates synaptic vesicle membrane trafficking, and is implicated in the neuropathy CEDNIK; *ELP1*, which regulates the migration and branching of projection neurons in the developing cerebral cortex and is implicated in Charcot-Marie-Tooth (CMT) disease; and *IGHMBP2*, also implicated in CMT disease.

The enriched BP GO term in the BM gene set, "negative regulation of glycoprotein metabolic process" (3 genes, 13.3 fold enrichment), contains two genes with a role in the regulation of amyloid-beta precursor protein (APP). *ITM2B* regulates APP processing and acts as an inhibitor of the amyloid-beta peptide aggregation and fibrils deposition, and *ITM2C* is a regulator of amyloid-beta peptide production, with the products of both genes blocking access of secretases to the APP cleavage site and implicated in various forms of dementia and cerebral amyloid angiopathy.

Other interesting genes in the BM gene set include *FOXN4*, a forkhead transcription factor essential for the development of neural tissues, particularly the retina and spinal cord; *MYRF*, which encodes a transcription factor that is required for CNS myelination, specifically activating transcription of CNS myelin genes, and may regulate oligodendrocyte differentiation; *DOK5* and *NRN1L*, which play roles in neurite outgrowth; *STMN2* which plays a regulatory role in neuronal growth and is associated with Down's syndrome and AD; *KIDINS220*, which is preferentially expressed in the nervous system where it controls neuronal cell survival, dendrite differentiation, synaptic plasticity, and axon guidance, and is associated with various neuropsychiatric disorders and neurodegenerative diseases including AD; *5HTR3B*, which encodes subunit B of the type 3 receptor for the biogenic hormone, serotonin, with this receptor causing fast depolarising responses in neurons after activation; and *NMUR2*, which encodes a G-protein coupled receptor (GPCR) for the neuromedin-U and neuromedin-S neuropeptides, which are widely distributed in the gut and CNS, playing an important role in the regulation of food intake and body weight. Another interesting gene in the BSM gene set is *GDF2*, which regulates the differentiation of cholinergic neurons, along with other functions.

#### 3.5 Lipid and other metabolic processes

Several enriched GO terms in the BSM gene set relate to metabolic processes and specifically to lipid metabolism including the BP GO terms triglyceride, acylglycerol, and lipid homeostasis with three genes; *APOE*, *CETP*, and *GCKR*. Two of these genes also comprise the BSM GO terms "triglyceride-rich lipoprotein particle remodelling" and "very-low-density lipoprotein particle remodelling” (2 genes, 44.9 fold enrichment). The majority of the most enriched disease terms for the BSM gene set are related to lipid metabolism including various terms describing LDL and HDL cholesterol, lipid profiles and levels, hyperglyceridaemia, waist circumference, and heart diseases. Similarly, the BM gene set contains related enriched terms including the BP GO terms “ketone biosynthetic process”, “glycerol metabolic process”, and “alditol metabolic process” (3 to 4 genes, 7.4 to 11.1 fold enrichment), the enriched UP keyword “obesity” (4 genes, 5.2 fold enrichment), and enriched disease annotations “lipid profiles” and “dyslipidaemias | hypertriglyceridemia” (4 and 3 genes, 12.8 and 14.1 fold enrichment).

There are also two BM clusters (enrichment scores ~1.6) of BP GO terms with broad/general terms regarding the regulation of metabolic processes (47 to 111 genes, 1.2 to 1.4 fold enrichment), as well as other broad individual GO terms including “positive regulation of nitrogen compound metabolic process”, “carbohydrate derivative metabolic process”, and “positive/negative regulation of nucleobase-containing compound metabolic process” (29 to 38 genes, 1.4 to 1.5 fold enrichment). The second cluster for the BSM gene set (enrichment score 1.62) also contains four broad metabolic process related BP terms (50 to 52 genes, 1.2 fold enrichment), as well as other similar individual GO terms not in the cluster.

#### 3.6 Other

There are several enriched GO terms for the BM gene set relating to DNA replication and repair including “DNA replication” (11 genes, 2.5 fold enrichment), “DNA repair” (15 genes, 1.9 fold enrichment), “cellular response to DNA damage stimulus” (20 genes, 1.6 fold enrichment), “post replication repair” (4 genes, 4.8 fold enrichment), and the MF term “ERCC4-ERCC1 complex” (2 genes, 45.5 fold enrichment), as well as the UP keyword and KEGG pathway “DNA replication” (5 and 3 genes, 3.8 and 8.4 fold enrichment).

Another BM cluster (enrichment score 1.7) includes several BP GO terms related to the regulation of sodium ion transport (4 to 5 genes, 4 to 6.3 fold enrichment), with further overlapping terms including the MF GO term “ion channel binding” (6 genes, 3.6 fold enrichment). For the BM gene set, there are general gene regulatory signatures including the UP keyword “activator” (18 genes, 1.9 fold enrichment), and the MF GO terms “transcription corepressor activity” (9 genes, 2.7 fold enrichment) and “RNA polymerase II transcription factor activity sequence-specific DNA binding” (17 genes, 1.7 fold enrichment). Other enriched BM terms include the BP term “enzyme linked receptor protein signalling pathway” (25 genes, 1.7 fold enrichment), the CC terms “proteinaceous extracellular matrix” (12 genes, 2.3 fold enrichment) and “Golgi associated vesicle” (5 genes, 4.1 fold enrichment), the MF terms “sulphur compound binding” (10 genes, 2.9 fold enrichment) and “heparin binding” (7 genes, 3 fold enrichment), and the UP keywords “signal-anchor” (13 genes, 2 fold enrichment) and “extracellular matrix” (9 genes, 2.5 fold enrichment). Other interesting BM genes include *TYR*, tyrosinase, which plays a role in the formation of pigments such as melanin; and *VEZT*, which is involved in morphogenesis of the preimplantation embryo and the implantation process.

For the BSM gene set, two other interesting GO terms are “T cell differentiation in thymus” and “thymocyte aggregation” (11.6 and 10.7 fold enrichment) with three genes (*CCR7*, *NKAP*, and *ITPKB*), and another enriched disease annotation is “HIV Infections | human immunodeficiency virus disease” (7 genes, 5 fold enrichment). Another notable gene in the BSM gene set is *NUCB2*, which may play a role in calcium level maintenance, eating regulation in the hypothalamus, and release of tumour necrosis factor from vascular endothelial cells.

**4) Extended results: Gracile capuchins (*Cebus*; H2) positive selection results**

Lists of all enriched annotated terms and GO clusters including annotation category, term description and ID, gene counts and hits, and statistical results such as EASE score and fold enrichment, for the BM and BSM gene set enrichment analyses for gracile capuchins (*Cebus*: H2) are found in Tables S28–30.

<u>4.1 Limb and skeletal system development</u>

For *Cebus*, the BM gene set is enriched for genes related to limb and skeletal system development. The top GO cluster in DAVID for the BM gene set (enrichment score 1.97) contains six BP GO terms: limb / appendage morphogenesis / development, and embryonic limb / appendage morphogenesis, which are also individually among the most enriched BP GO terms (3.3 to 4.3 fold enrichment). These GO terms are hit by the same seven genes in this gene set: *RSPO2*, *TBC1D32*, *C5orf42*, *HAND2*, *HOXC11*, *HOXD10*, and *SHOX2*. Many of these genes are known to play a crucial role in embryonic limb development including *RSPO2*, which is involved in limb specification through amplification of the Wnt signalling pathway. There are three homeobox genes of the Hox and Shox families that play fundamental roles in embryonic pattern formation and are required for normal limb development and growth including the short homeobox gene, *SHOX2*, which is expressed in the developing stylopod, and the Hox transcription factors, *HOXD10* and *HOXC11*, with the former expressed in the developing limb bud (*78*, *142*). Several genes are involved in Shh signalling such as *HAND2*, which functions as an upstream regulator of Shh induction in the limb bud; *TBC1D32*, which is required for high-level Shh responses in the developing neural tube, and plays a role in control of primary cilium structure allowing *GLI2* to be properly activated; and *C5orf42* (*CPLANE1*), which is involved in ciliogenesis and therefore important in Shh signalling. Shh signalling plays a central role in limb development in the establishment of anterior-posterior polarisation in both a concentration- (paracrine) and time-dependent (autocrine) manner involving complex spatiotemporal regulation, with altered Shh signalling implicated in disorders with congenital limb defects and in the evolution of the morphological diversity of vertebrate limbs (*143*). Mutations in genes in this cluster are associated with various skeletal dysmorphologies and congenital limb defects in humans including vertical talus (*HOXC11*, *HOXD10*), humerofemoral hypoplasia (*RSPO2*), and orofaciodigital syndrome (*TBC1D32*, *C5orf42*) while *SHOX2* is implicated in the short stature of Turner syndrome.

Other related enriched GO annotations for the BM gene set include "embryonic skeletal system morphogenesis" (4.3 fold enrichment) with five genes including *SHOX2*, *HOXC11*, and *HOXD10* from previous cluster as well as two other Hox genes, *HOXB5* and *HOXB2*; "skeletal system development" (13 genes, 2.1 fold enrichment), including the genes *SH3PXD2B*, *HAND1*, *NOV*, *PEX7* and *TRIM45*; "cartilage morphogenesis" (3 genes, 19.8 fold enrichment); and "connective tissue development" (8 genes, 2.7 fold enrichment). It is notable that these limb and skeletal system development GO terms include five accelerated Hox genes from a total of 21 Hox genes analysed for *Cebus* in BM tests. Hox genes encode transcription factors that regulate the expression of downstream target genes to control axes during development and are required to promote the proliferation and differentiation of skeletal progenitor cells in the mesoderm, as well as the recruitment of mesenchymal cells into pre-cartilaginous condensations during limb development (*144*, *145*).

Some of the other genes in these skeletal system related GO terms include *SH3PXD2B*, which encodes an adapter protein required for podosome formation involved in cell adhesion and migration, with mutations in this gene associated with skeletal dysplasia found in Frank-Ter Haar syndrome (*146*); *HAND1* encodes a basic helix-loop-helix transcription factor that is involved in the development and morphogenesis of long bones, and regulates bone size and morphology by suppressing postnatal expression of collagen fibrils in the cortical bones (*147*); *PEX7* (also in the BSM gene set) plays an essential role in peroxisomal protein import with mutations in this gene causing rhizomelic chondrodysplasia punctata type 1, which is characterised by disturbed endochondral bone formation, shortening of the femur and humerus, vertebral disorders, dwarfism, facial dysmorphism, and intellectual disabilities; and *NOV* (*CCN3*) encodes a member of the CCN family of regulatory proteins with various roles in regulating cells within the bone microenvironment including promoting osteoclast and chondrocyte differentiation, and impairing osteoblast differentiation by neutralising BMP and Wnt and activating Notch signalling (*148*).

Other interesting genes in the BM gene set not in these GO terms include *SFRP4*, which functions as a modulator of Wnt signalling, playing a role in bone morphogenesis during post-natal development, and is associated with Pyle's disease, which is characterised by cortical-bone thinning, limb deformity, and fractures (*149*, *150*); *LEO1*, a component of the PAF1 complex required for transcription of Hox and Wnt target genes; *CRTAC1*, which encodes an ECM protein found in articular deep zone cartilage; and *PRG4*, which encodes a large proteoglycan made by chondrocytes located at the surface of articular cartilage, functioning as a boundary lubricant and contributing to the elastic absorption and energy dissipation of synovial fluid. Mutations in *PRG4* cause CACP syndrome which is characterised by the childhood onset joint abnormalities.

#### 4.2 Embryonic development

There are several enriched GO terms related to vasculature and heart development in the *Cebus* BM gene set including "coronary vasculature development" (5 genes, 9.2 fold enrichment), "cardiac chamber development" (8 genes, 4.1 fold enrichment), "cardiac septum development" (6 genes, 5 fold enrichment), "vasculature development" (14 genes, 1.8 fold enrichment), and "regulation of vascular endothelial growth factor receptor signalling pathway" (3 genes, 8.8 fold enrichment), as well as the enriched UP keyword "angiogenesis" (6 genes, 4 fold enrichment). Another related GO term is regulation of "p38MAPK cascade" (3 genes, 8.5 fold enrichment), which is a major endothelial cell signalling pathway.

Several of the genes found in these GO terms, particularly in the cardiac terms, overlap with the limb and skeletal development GO terms, such as *HAND1* and *HAND2*. These basic helix-loop-helix genes are also essential for cardiac morphogenesis, particularly for the formation of the right ventricle and of the aortic arch arteries, with *HAND2* also required for vascular development and regulation of angiogenesis likely through a VEGF signalling pathway. Notably, cardiac muscle and blood vessel endothelium develop from the mesoderm, along with bone and cartilage, which may underly some of these overlapping developmental signatures. Some other genes in these vasculature and cardiac terms include *ID2*, which negatively regulates basic helix-loop-helix transcription factors (such as *HAND1* and *HAND2*), inhibiting skeletal muscle and cardiac myocyte differentiation, and is implicated in the regulation of angiogenesis; *VEGFA*, which encodes a member of the PDGF/VEGF growth factor family essential for angiogenesis and endothelial cell growth; *MMRN2*, which inhibits endothelial cells motility and acts as a negative regulator of angiogenesis by binding *VEGFA*; *CCM2*, a scaffold protein of the CCM signalling pathway and a crucial regulator of heart/vessel formation; and *MFGE8* and *ANGPTL6*, both involved in neovascularisation.

More generally, there are recurring signatures of accelerated evolution in genes involved in embryonic development in *Cebus*, for example, the third most significantly enriched BP GO annotation is "chordate embryonic development" with 17 genes (2.3 fold enrichment), and the sixth is "embryo development" (23 genes, 1.9 fold enrichment), among other highly similar terms. Notably, 13 of the top 14 BP GO terms describe the development or morphogenesis of the embryo, limbs, or heart. Similarly, one of the most enriched UP keywords is "developmental protein" with 19 genes (1.7 fold enrichment), including many of those noted in this and the previous sections.

#### 4.3 Endosomes and vacuoles

There are five enriched CC GO terms related to endosomes and vacuoles including "endosome", "endosome membrane", and "endosomal part". Similarly, the most enriched individual BP GO term is "vacuole organisation", (10 genes, 3.7 fold enrichment) sharing many genes with the endosome CC terms. Endosomes play active roles in many important physiological processes including nerve impulse transmission, and the importance of endosomes for proper brain function is underscored by the implication of endosome dysfunction in many neurodegenerative disorders including AD. One of the genes in these terms is *GM2A*, which plays a role in binding gangliosides and stimulating ganglioside GM2 degradation. Gangliosides are highly abundant in the nervous system, and their importance in the brain is highlighted by the severe neurodegenerative disorders (e.g., Tay-Sachs Disease) caused by loss of function mutations in ganglioside biosynthetic enzymes (*151*). Another related gene in the BM and BSM gene sets, though not in these GO terms, is *PLAA* which plays a role in synaptic vesicle recycling through the trafficking of ubiquitin-mediated membrane proteins to late endosomes, as well as a role in cerebellar Purkinje cell development, and is associated with several neurodevelopmental disorders. An interesting gene in the vacuole organisation GO annotation is *PINK1*, which protects against mitochondrial dysfunction during cellular stress by phosphorylating mitochondrial proteins and is implicated in Parkinson's Disease.

#### 4.4 Brain and neuronal-related

Some interesting genes annotated by the UP keyword "developmental protein" are involved in neural and brain development including *TBX6*, a T-box transcription factor that plays an essential role in determining the neural vs mesodermal fate of axial stem cells; *VEGFA*, which initiates a signalling pathway needed for motor neuron axon guidance including for the caudal migration of facial motor neurons during embryonic development; *ZNF335*, with an important role in neural progenitor cell proliferation and self-renewal through the regulation of specific genes involved in brain development; *ATOH8*, a transcription factor with many roles including in the specification and differentiation of neuronal cell lineages in the brain; *SIX6*, which is required to maintain expression of gonadotrophin-releasing hormone (GnRH), and for the development and survival of GnRH neurons, which themselves are crucial to the hypothalamic-pituitary-gonadal system that regulates mammalian fertility, and may also be involved in eye development (*152*); and finally, *ID2* is notable for its role in regulating the circadian clock.

A related gene among those found in both gene sets is *SSPO*, which may play a role in neurogenesis in early brain development and the formation of the CNS (*153*). Notable genes in the BSM gene set include *BCAN*, a CSPG specifically expressed in the CNS, serving as guidance cues during development and modulating synaptic connections postnatally; and *DLX4*, a distal-less homeobox gene which are postulated to play a role in forebrain and craniofacial development.

For the BM set, other interesting genes include *KHDRBS1* (2nd ranked), which may regulate alternative splicing of *NRXN1* and *NRXN3*, important cell surface receptors involved in neurotransmission; *ULK4*, which encodes a protein involved in neurite branching, neurite elongation, and neuronal migration, and is associated with schizophrenia and bipolar disorder; *VSTM5*, which plays several important roles including in modulating the position and complexity of central neurons, the formation of neuronal dendrites, and regulation of neuronal morphogenesis and migration during cortical development in the brain; *RASSF10*, which plays an important role in regulating embryonic neurogenesis; *NPFFR1*, a receptor for NPAF and NPFF neuropeptides implicated in hormonal modulation, regulation of food intake, thermoregulation, and nociception; *BSX*, brain specific homeobox, a DNA binding protein that functions as a transcriptional activator, essential for normal postnatal growth and nursing, and is an essential factor for the function of neuronal neuropeptide Y and agouti-related peptide; *HTR1E*, a GPCR for serotonin which is primarily located in the frontal cortex, caudate putamen, claustrum, hippocampus, and amygdala; *SSTR1*, a receptor for the peptide hormone somatostatin that regulates diverse cellular functions such as neurotransmission, cell proliferation, and endocrine signalling, as well as inhibiting the release of various hormones and other secretory proteins; and *P2RX2*, which encodes a gated ion channel involved in a variety of processes such as excitatory postsynaptic responses in sensory neurons, neuromuscular junction formation, perception of taste, peristalsis, and auditory neurotransmission.

4.5 Other

There are several enriched annotations in the BM gene set related to DNA damage/repair including the BP GO term “double strand break repair” (7 genes, 2.7 fold enrichment), and the UP keywords “DNA repair” and “DNA damage” (9 and 10 genes, 2.6 and 2. 4 fold enrichment), and “mutator protein” (2 genes, 55.8 fold enrichment).

There are several enriched GO terms related to pigment-related biological processes for the BM gene set including “heme / pigment biosynthetic process” (3 and 4 genes, 10.8 and 5.9 fold enrichment), which together form the second GO cluster (enrichment score 1.47). One of the genes in these GO terms is *TYRP1*, which encodes a melanosomal enzyme that plays an important role in the melanin biosynthetic pathway, and is implicated in various forms of albinism. Another gene not in this cluster but in both gene sets is *HPS3*, which is involved in melanosome biogenesis and associated with Hermansky-Pudlak Syndrome, characterised by oculocutaneous albinism causing light pigmentation of the skin, hair, and eyes. Other genes in these GO terms are putatively related to the respiratory pigment heme (*COX10*, *UROS*) and a related gene in the BM set is *TSPO*, which may play a role in the transport of heme.

Other interesting genes that are among the 18 found in both gene sets that have not been mentioned include *CLUL1*, which encodes a glycoprotein that is expressed predominantly by cone photoreceptors of the retina, and *SLC19A1*, which is a folate transporter and involved in the regulation of intracellular concentrations of folate. Another folate transporter found in the BM gene set, *SLC46A1*, is expressed in the brain and choroid plexus where it transports folates into the CNS. Across the BM gene set, there are at least seven genes implicated in deafness: *OPA1*, *DFNB59*, *HOMER2*, *PEX7*, *P2RX2*, *PJVK*, and *TMPRSS3*. The sole enriched disease annotation for the BM gene set is “menarche” (4 genes, 6.1 fold enrichment).

There are broad cellular metabolism related BP GO terms in the BM gene set such as “cellular catabolic process”, “cellular macromolecule catabolic process”, “organic cyclic compound catabolic process”, and “RNA catabolic process” (8 to 34 genes, 1.3 to 2.5 fold enrichment). Other enriched terms for the BM gene set include the BP GO terms “negative regulation of mitochondrion organisation” (4 genes, 7.4 fold enrichment), “respiratory chain complex IV assembly” (3 genes, 9.9 fold enrichment), “regulation of mitophagy” (4 genes, 7 fold enrichment), and “autophagy” (12 genes, 2.3 fold enrichment), the MF GO terms “activating transcription factor binding” (5 genes, 6.9 fold enrichment), “polysaccharide binding” (3 genes 11.1 fold enrichment), and “amide binding” (8 genes, 2.5 fold enrichment), and the UP keywords “protein transport” (14 genes, 1.9 fold enrichment) and “ANK repeat” (8 genes, 2.5 fold enrichment).

For the BSM gene set, there are only four enriched GO terms with the CC terms “microbody” and “peroxisome” (4 genes, 8.7 fold enrichment) and “cilium” (6 genes, 4.3 fold enrichment), and the MF term “ligase activity” (6 genes, 3.4 fold enrichment), with no significant GO clusters or enriched BP GO terms, KEGG or Reactome pathways, or disease annotations for this gene set. Enriched UP keywords for the BSM gene set include “cilium” (6 genes, 8 fold enrichment), “ligase” (6 genes, 4.7 fold enrichment), “leucine-rich repeat” (5 genes, 4.4 fold enrichment), “cell projection” (7 genes, 2.7 fold enrichment), and “immunoglobulin domain” (6 genes, 3 fold enrichment).

**5) Extended results: Ancestral Cebinae (H3) positive selection results**

Lists of all enriched annotated terms and GO clusters including annotation category, term description and ID, gene counts and hits, and statistical results such as EASE score and fold enrichment, for the BM and BSM gene set enrichment analyses for ancestral Cebinae (H3) are found in Tables S31–34.

5.1 Mitochondria

For ancestral Cebinae, the BM gene set is enriched for genes related to the mitochondrion. The top GO annotation cluster in DAVID for the BM gene set with an enrichment score of 1.96 contains four CC GO terms: “mitochondrial inner membrane”, “mitochondrial envelope”, “mitochondrial membrane”, and “organelle inner membrane” (16 to 20 genes, 1.8 to 2.1 fold enrichment). Similarly, many of the other most significantly enriched GO terms are related to the mitochondrion including the CC GO terms “mitochondrion” (1$^{st}$; 46 genes, 1.8 fold enrichment) and “mitochondrial part” (4$^{th}$; 28 genes, 19 fold enrichment), and the BP GO term “mitochondrion organisation” (8$^{th}$; 20 genes, 2 fold enrichment). Other enriched mitochondrial GO terms include the BP term “mitochondrial morphogenesis” (3 genes, 10.3 fold enrichment) and the CC term “mitochondrial matrix” (14 genes, 2.2 fold enrichment).

These GO terms cover several genes that are found in both gene sets including *BAX*, which plays a role in the mitochondrial apoptotic process, and *SUPV3L1*, which is a major helicase involved in mitochondrial RNA metabolism, serving as an assembly factor that is required for formation of the membrane arm of the complex. In addition, other genes highly ranked in the BM gene set include *FOXRED1* (6$^{th}$), which is required for assembly of mitochondrial complex I; *COX15* (8$^{th}$), which may be required for the biogenesis of the terminal component of the mitochondrial respiratory chain (COX); *ETFDH* (11$^{th}$), which encodes a component of the electron-transfer system; and *LRPPRC* (15$^{th}$) which localises primarily to mitochondria and plays a role in translation or stability of mitochondrially encoded COX subunits. “Mitochondrion” is also the most significantly enriched UP keyword (33 genes, 2 fold enrichment), with many of the same genes also found in another enriched UP keyword “transit peptide” (18 genes, 2.3 fold enrichment), including *TWNK*, a gene involved in mitochondrial DNA (mtDNA) metabolism, considered critical for lifetime maintenance of mtDNA integrity, and a key regulator of mtDNA copy number in mammals.

Several of these aforementioned genes (*BAX*, *SUPV3L1*, *FOXRED1*, *ETFDH*) are included in more specific mitochondrial-related GO term such as the CC term "mitochondrial protein complex" (8 genes, 3.1 fold enrichment), along with *NDUFB9*, *NDUFS6*, *MCCC1*, and *MCCC2*, with the latter two genes comprising the related CC term "3-methylcrotonyl-CoA carboxylase complex mitochondrial" and MF term "methylcrotonoyl-CoA carboxylase activity" (~65.8 fold enrichment). This signal is also found in disease annotations with "mitochondrial complex I deficiency" as the most enriched disease term containing three of these genes (13.5 fold enrichment). A relevant gene, highly ranked in both gene sets and annotated by the mitochondrial membrane related CC GO terms, is *TMEM126B*, encoding a mitochondrial transmembrane protein component of the assembly complex for mitochondrial complex I. Another gene in both gene sets but not found in these GO terms is *TMEM135*, which is involved in mitochondrial metabolism through regulation of the balance between mitochondrial fusion and fission, with a related gene in the BSM set, *GDAP1*, which regulates the mitochondrial network and promotes mitochondrial fission. The balance of mitochondrial fission and fusion dynamics may be especially important in neurons given its association with impaired development of the nervous system in humans (*154*).

In the BM gene set for ancestral Cebinae, there are interesting related enriched disease annotations; "cognitive trait" with six genes, and "aging" and "telomere length" with the same six genes (both 2.9 fold enrichment), though the EASE scores for the latter two are just above significance (0.057 and 0.059). These genes are *NDUFB9*, *NDUFS6*, *GSTO2*, *GSR*, *SLC25A27*, and *UCP2*, with *GSTO2* the only gene not found in a mitochondrion related GO term. Both NADH:ubiquinone oxidoreductase genes (*NDUFB9*, *NDUFS6*) are central to the mitochondrial complex 1 and implicated in adult onset neurodegenerative disorders. The two mitochondrial uncoupling proteins (UCPs), *SLC25A27* and *UCP2*, create proton leaks across the inner mitochondrial membrane, thus uncoupling oxidative phosphorylation from ATP synthesis, and may play a role in thermoregulatory heat production and metabolism in brain. Another gene related to telomere length in the BM gene set is *STN1*, which encodes a component of a complex that protects telomeres from DNA degradation and functions in telomere replication and length homeostasis.

#### 5.2 Hormones, neuropeptides, and other neuromodulators

For ancestral Cebinae, the BM gene set is enriched for genes related to hormones. The third cluster for the BM gene set is a large cluster containing nine BP GO terms (enrichment score 1.56) including “hormone secretion”, “hormone transport”, “signal release”, “peptide secretion”, and the terms describing the regulation of these processes (8 to 13 genes, 2 to 2.6 fold enrichment). Related enriched terms not in this cluster include the MF term “hormone activity” (6 genes, 3.3 fold enrichment), the BP term “negative regulation of secretion by cell” (7 genes, 2.7 fold enrichment), and the Reactome pathway “androgen biosynthesis” (3 genes, 19.5 fold enrichment). Interesting hormone related genes in these annotations include *SOX4*, which may mediate downstream effects of parathyroid hormone in bone development; *HMGN3*, which binds thyroid hormone receptor beta in the presence of thyroid hormone (TH), impacts insulin and glucagon levels, and modulates the expression of pancreatic genes involved in insulin secretion; *TRH*, which is involved in the secretion of thyroid-stimulating hormone (TSH), TH synthesis regulation, and the modulation of hair growth; *CGA*, which encodes the alpha subunit of the four pituitary glycoprotein hormones (chorionic gonadotropin, luteinising hormone, FSH, and TSH); *HSD17B12*, which encodes a hydroxysteroid that converts oestrone into oestradiol in ovarian tissue; and *SRD5A3*, involved in the production of androgen 5-alpha-dihydrotestosterone from testosterone, and maintenance of the androgen-androgen receptor activation pathway. Another notable BM gene is *POU1F1*, a member of the POU family of transcription factors that regulate mammalian development. *POU1F1* is involved in pituitary development through the specification of somatotroph, lactotroph, and thyrotroph cells (growth hormone, prolactin, and TSH producing cells, respectively) in the developing anterior pituitary, and in hormonal expression and the activation of growth hormone and prolactin genes. There are two genes encoding synaptotagmins in this cluster that are also found in the across-capuchins gene set (*STY11* and *SYT3*).

Several of the genes in the BM hormone/peptide related cluster and GO terms are neuropeptides or other neuromodulators, or their receptors. *NPFF*, which encodes neuropeptide FF, is involved in a range of physiologic roles including nociception and pain modulation, and in the central processing of visceral autonomic signals related to feeding, cardiovascular responses, stress, neuroendocrine regulation, and hormonal modulation (*155*). *NPVF*, also found in the BSM gene set, encodes a propeptide that is cleaved to form the neuropeptides NPSF and NPVF, also referred to as the RFamide-related peptides RFRP-1 and RFRP-3, respectively, which are mammalian homologs of the avian neuropeptide gonadotropin-inhibitory hormone (GnIH). These neuropeptides act as potent negative regulators of gonadotropin synthesis and secretion with a range of functions in the modulation of reproduction, which appear to vary across lineages but linked to the regulation of sexual behaviour, sexual maturation, ovulatory cycle, gonadal function, reproductive seasonality, and stress induced reproductive suppression, among others, as well as a role in nociception and sleep regulation (*156*, *157*). *CARTPT* encodes the propeptide for the neuropeptide CART that is expressed abundantly in the brain, functions as a neuromodulator of dopamine signalling, and plays a role in appetite, energy balance, maintenance of body weight, reward, addiction, and the stress response. *RLN3* encodes a member of the relaxin family that is expressed predominantly in the brain and modulates a range of physiological processes such as stress, arousal, memory, and appetite regulation. There are also two serotonin receptors in the gene sets: *HTR5B* (BSM gene set) which is pseudogenised in humans but has important functions in mice, and *HTR1F* (BM gene set), which is located primarily in the hippocampus, cortex, and dorsal raphe nucleus. An interesting neuromodulatory gene in the BSM gene set is *CALY*, which interacts with the D1 dopamine receptor and may interact with other dopamine receptor subtypes.

Among the top most enriched UP terms for the BM gene set is "neuropeptide" with four genes (10.6 fold enrichment), including *CARTPT*, *NPVF* and *NPFF*, as well as *PROK2*. The other prokineticin gene, *PROK1*, is found in the BM gene set. Prokineticins are widely expressed with *PROK1* predominantly expressed in peripheral tissues, especially steroidogenic organs, and *PROK2* is mainly expressed in the CNS. Prokineticin signalling has been associated with many important functions including the contraction of gastrointestinal smooth muscle, circadian rhythm regulation (discussed below), neurogenesis, angiogenesis, pain perception, mood regulation, and reproduction, and their dysregulation has been associated with diseases such as neurodegeneration and cancer (*158*). Among the enriched terms in the BM gene set is the Reactome pathway "agmatine biosynthesis" (71.5 fold enrichment) with two genes, *AGMAT* and *AZIN2*, and an identical BP GO term (65.3 fold enrichment). Another similar enriched BP GO annotation is "cellular biogenic amine metabolic process" (4 genes, 5.4 fold enrichment), which includes *AGMAT* and *AZIN2*, as well as *TRH* and *CHKA*.

5.3 Extracellular matrix of the brain

*NCAN*, the 4th ranked gene in the BSM gene set, and *BCAN*, the 2nd ranked gene in the BM gene set and also in the BSM gene set, encode CSPGs of the lectican family that are specifically expressed in the CNS serving as guidance cues during development and modulating synaptic connections in the adult. They are abundant components of the brain's ECM forming a condensed lattice-like structure known as perineuronal nets (PNNs) that play important roles in many diverse CNS functions. *NCAN* may also modulate neuronal adhesion and neurite growth during development by binding to neural cell adhesion molecules, and is implicated in the psychological condition known as Capgras syndrome. All of the most enriched Reactome pathways for the BSM gene set are a result of *BCAN* and *NCAN*, including the pathway "ECM proteoglycans" (9.1 fold enrichment), which also includes *VTN*. *VTN* encodes a glycoprotein that is marker for brain-resident pericytes, which play a crucial role in the formation and functionality of the blood-brain barrier, and is also expressed by pericytes of subventricular zone in the mouse where neurogenesis continues throughout life (*159*).

5.4 CNS development and other neuronal signatures

For the BSM gene set, the BP GO term "central nervous system development" is enriched (2 fold enrichment) and contains 10 interesting genes: *BAX*, *BRCA2*, *BBS7*, *GDF7*, *KNDC1*, *SPEF2*, and *TPP1*, as well as the three genes mentioned in the brain's ECM section, *BCAN*, *NCAN*, and *VTN*. *BRCA2* is involved in maintenance of genome stability, specifically the homologous recombination pathway for double-strand DNA repair, and required for neurogenesis particularly during embryonic and postnatal neural development (*160*). *GDF7* may play an active role in the motor area of the primate neocortex, and contributes to neuronal cell identity in its selective expression in the roof plate in the developing embryonic nervous system, inducing the formation of sensory neurons (*161*). *KNDC1* encodes brain specific Ras guanine nucleotide exchange factor that controls the negative regulation of neuronal dendrite growth, may be involved in cellular senescence, and likely serves an important role in regulating neuronal dendrite development. *BBS7* is required for ciliogenesis and discussed more in the cilium-related section below. Another cilium-related gene, *SPEF2*, is required for motile cilia function, which also play an important role in the brain. *BAX* plays a role in the mitochondrial apoptotic process with potentially crucial functions (though somewhat redundant with *BAK*) during development including maintenance of homeostatic mitochondrial morphology, which is essential for proper development of cortex neurons (*162*). *TPP1* encodes a lysosomal serine protease implicated in CLN2 disease, which is characterised by epilepsy, language development delay, visual impairment, and developmental regression.

Other genes not included in this GO term but found in both gene sets are *LNPK*, which is involved in CNS development and implicated in neurodevelopmental disorders; *ANKRD11*, which has a role in proliferation and development of cortical neural precursors; and *FBXW8*, involved in dendrite patterning in the brain. Interesting genes in the BSM gene set include *ENAH* (3rd rank), which encodes a protein involved in a range of processes dependent on cytoskeleton remodelling and cell polarity such as axon guidance and lamellipodial/filopodial dynamics in migrating cells; *GPR37*, which encodes a receptor for the neuro- and glio-protective factor prosaposin, and is associated with both juvenile and late onset Parkinson's disease; and *GDAP1*, which may play a role in signal transduction during neuronal development.

Other interesting genes in the BM gene set include *DBX1*, developing brain homeobox 1, a transcription factor proposed to play a role in patterning of the CNS during embryogenesis; *POU1F1*, involved in the specification of somatotroph, lactotroph, and thyrotroph cells in the developing anterior pituitary; *ADGRG6*, which is essential for normal differentiation of promyelinating Schwann cells and for normal myelination of axons, and regulates neural, cardiac, and ear development; *ARTN*, which encodes a glial cell derived neurotrophic factor member that supports the survival of sensory and sympathetic peripheral neurons and also supports the survival of CNS dopaminergic neurons of the ventral mid-brain; *TRIM44*, which may play a role in the differentiation and maturation of neuronal cells, and act as a negative regulator of *PAX6* expression; *CXXC5*, which acts as a mediator of Wnt signalling activity in neural stem cells, among other functions; and *AHSG*, which is involved in brain development as well as the formation of bone tissue and endocytosis.

#### 5.5 Circadian rhythms

There are signatures of selection on circadian rhythms in the BM gene set including as an enriched BP GO term (6th; 3 fold enrichment) with eight genes: *PROK1*, *PROK2*, *PER3*, *TIMELESS*, *METTL3*, *CARTPT*, *BHLHE41*, and *HNRNPD*. Six of these genes are also annotated by the enriched UP keyword "biological rhythms" (3.3 fold enrichment). Interestingly, *PER3* is found in both gene sets and is the top ranked most significant gene in the BSM gene set. *PER3* is a core component of the circadian clock expressed in a circadian pattern in the suprachiasmatic nucleus (SCN), the primary circadian pacemaker in the mammalian brain. It is a member of the Period family that encode components of the circadian rhythms of locomotor activity, metabolism, and behaviour. *PROK2* is thought to function as an output molecule from the SCN that transmits behavioural circadian rhythms. *TIMELESS* interacts with Period genes in its role in the autoregulatory loop of the circadian rhythm, and is associated with psychiatric disorders such as bipolar disorder. *METTL3* plays a role in the regulation of various processes including the circadian clock, as well as differentiation of embryonic and haematopoietic stem cells, cortical neurogenesis, response to DNA damage, and primary miRNA processing. *BHLHE41* acts as a transcriptional repressor involved in the regulation of the circadian rhythm by repressing the activity of the clock and clock-controlled genes. *HNRNPD* plays a role in the regulation of the rhythmic expression of circadian clock core genes.

#### 5.6 Fertilisation

For ancestral Cebinae, the top ranked individual GO term in the BSM gene set is “fertilisation” (5.5 fold enrichment) with five genes: *BAX*, *CATSPER3*, *FETUB*, *PRSS7*, and *SPEF2*. Three of these genes are found in both gene sets (*BAX*, *CATSPER3*, *PRSS37*). *CATSPER3* encodes a voltage-gated calcium ion channel that plays a central role in calcium-dependent physiological responses such as sperm hyperactivation, acrosome reaction, and chemotaxis towards the oocyte, which are essential for successful fertilisation. *PRSS37* is involved in the activation of the proacrosin/acrosin system, may play a role in sperm migration or binding to zona-intact eggs, and is implicated in male fertility. *FETUB* encodes a protease inhibitor required to prevent premature zona pellucida hardening before fertilisation. *SPEF2* is important for development of the axoneme, manchette, and sperm head, and is essential for male fertility. Three of these genes (*CATSPER3*, *FETUB*, *PRSS7*) comprise another enriched BP GO term “sperm-cell recognition” (11.4 fold enrichment).

Other related Cilia And Flagella Associated Protein (CFAP) genes not found in these GO terms are *CFAP65* (BSM gene set), which plays a role in flagellar formation and sperm motility, and *CFAP70* (BM gene set), which is an axoneme-binding protein that plays a role in the regulation of ciliary motility and cilium length. Both these genes are implicated in male infertility and sperm motility disorders. Related genes in the BM gene set include *ADCY10*, which induces the capacitation process that sperm undergo prior to fertilisation; and *TDRD6*, involved in the formation of chromatoid body (during spermiogenesis), Balbiani body (during oogenesis), and germ plasm (upon fertilisation).

5.7 Cilium

For the ancestral Cebinae BSM gene set, the top and only significant GO cluster (enrichment score 1.58) contains five terms related to the cilium with the CC terms “ciliary plasm” and “axoneme”, and the BP terms “cilium assembly”, “cilium organisation”, and “cilium morphogenesis” (4 to 5 genes, 3,7 to 7.2 fold enrichment). The five genes found across this cilium-specific cluster are *SPEF2* (discussed in the fertilisation section), *CEP162*, *BBS7*, *CFAP46*, and *TRAF3IP1*. While there is some overlap with the fertilisation GO, most (4 of 5) of the genes hit by these GO terms are distinct, with broader roles in primary cilium function and often implicated in ciliopathies. *BBS7* is required for proper assembly of the BBSome complex, which in turn is essential for ciliogenesis, and mutations in *BBS7* are implicated in Bardet-Biedl syndrome, a disorder with varying symptoms including obesity, retinal degeneration, polydactyly, intellectual disability, and nephropathy, among others. *CEP162* is required to promote assembly of the transition zone in primary cilia and implicated in Seckel syndrome, which is characterised by growth delays, dwarfism, microcephaly, intellectual disability, and unique facial features. *CFAP46* is important to the cilium axoneme and cilium movement, while *TRAF3IP1* plays a role in ciliogenesis.

For the BSM gene set, related enriched terms include the UP keyword “cilium” (5 genes, 4.1 fold enrichment), as well as the BP GO term “cell projection assembly” (7 genes, 3 fold enrichment), which contains all the genes in the cilium-related GO terms. Among the enriched BSM GO terms are less specific BP terms like “cell part morphogenesis” and “cellular component morphogenesis”, which include several genes discussed above related to fertilisation/sperm and cilium morphogenesis (11 and 14 genes, 2.5 and 1.9 fold enrichment). Other interesting BM genes are *TCTN3*, which encodes part of the tectonic-like complex required for tissue-specific ciliogenesis and Shh signalling; *CEP97*, which acts as a key negative regulator of ciliogenesis; and *KIAA0556* (*KATNIP*), also in the BSM gene set, which encodes a ciliary protein associated with Joubert Syndrome.

5.8 Embryonic development

Other signals for ancestral Cebinae in the BSM gene set include enriched BP GO terms "embryo development" (11 genes, 2.1 fold enrichment) and "embryonic digit morphogenesis" (3 genes, 8.8 fold enrichment). Interesting genes related to embryonic development in the BM gene set (aside from those mentioned previously) include *SUCO*, required for bone remodelling during late embryogenesis (also in the BSM gene set); *ZNF322*, a transcriptional activator important for maintenance of pluripotency in embryonic stem cells; *CTCF*, which plays a critical role in the epigenetic regulation and in activating/repressing transcription in oocyte/preimplantation embryo development; *CDX4*, which may regulate homeobox gene expression during patterning and haematopoiesis; *GPATCH3*, which may control neural crest cell migration involved in ocular and craniofacial development; *RNF111*, required for mesoderm patterning during embryonic development; *FOXN1*, a transcriptional regulator that regulates the development, differentiation, and function of thymic epithelial cells both in the prenatal and postnatal thymus; and *POU2F3*, which plays a critical role in keratinocyte proliferation and differentiation, and regulates expression of a number of genes including placental lactogen, which modifies the mother's metabolic state during pregnancy to facilitate the energy supply of the foetus. In the BM gene set, another enriched BP GO term is "labyrinthine layer development" (4 genes, 5.4 fold enrichment).

5.9 Metabolic processes & protein modification

Recurrent signatures of selection on various metabolic processes are found in both the BM and BSM gene sets. For the BM gene set, the fourth cluster (enrichment score 1.5) contains three BP GO terms related to organic acid metabolism (21 to 22 genes, 1.6 to 1.7 fold enrichment). Other enriched BP GO terms related to metabolic processes in the BM gene set include "primary amino compound metabolic process" (4 genes, 18.7 fold enrichment), "organonitrogen compound metabolic process" (47 genes, 1.4 fold enrichment), "sulphur compound biosynthetic process" (8 genes, 2.6 fold enrichment), "cellular amino acid metabolic process" (9 genes, 2.4 fold enrichment), "lipid modification" (10 genes, 2.5 fold enrichment), and "cellular lipid metabolic process" (25 genes, 1.6 fold enrichment). Other enriched terms for the BM gene set include the KEGG pathways "metabolic pathways" (25 genes, 1.5 fold enrichment), "valine, leucine and isoleucine degradation" (4 genes, 6.2 fold enrichment), and "other types of O-glycan biosynthesis" (3 genes, 10 fold enrichment), and the second most enriched disease category "waist-hip ratio" (8 genes, 2.5 fold enrichment).

Other related enriched BP terms for the BSM gene set include "positive regulation of cellular protein metabolic process", "positive regulation of protein modification", "regulation of proteolysis", "proteolysis", and "protein phosphorylation" (10 to 17 genes, 1.7 to 2.6 fold enrichment). Another enriched BSM GO term is "SCF ubiquitin-ligase complex" (10.2 fold enrichment) with three genes: *FBXO6* (6th ranked), *FBXW8* (also the top ranked gene in the BM gene set), and *ABTB1*. The SCF complex plays an important role in the ubiquitination of proteins involved in the cell cycle and in nearly all aspects of reproduction such as gametogenesis, oocyte-to-embryo transition, embryo development, and the regulation of oestrogen and progestin (*163*). Finally, the BM gene set also includes the enriched BP GO term "protein complex oligomerisation" (14 genes, 1.9 fold enrichment).

5.10 Other

The second cluster for the BM gene set (enrichment score 1.65) is comprised of four GO terms related to the immune response: "activation of immune response" (which is the 3rd ranked BP term), "immune response-activating signal transduction", "immune response regulating signal pathway", and "positive regulation of immune response" (15 to 18 genes, 1.7 to 2 fold enrichment). In addition, "acquired immunodeficiency syndrome | disease progression" is among the enriched disease categories (24 genes, 1.5 fold enrichment).

Other enriched terms for BM gene set include the MF GO term "G-protein coupled receptor binding" (10 genes, 2.6 fold enrichment), and the enriched Reactome pathway "G alpha (q) signalling events" (7 genes, 3 fold enrichment), which include several of the genes in the hormone/peptide GO cluster and terms. Other interesting genes in the BM gene set include *UACA*, which has been implicated in the regulation of mammary gland involution; two of the eight genes in humans that encode components of the troponin regulatory complex that regulates striated muscle contraction (*TNNT3* and *TNNC2*); and *CRYBA1*, which encodes crystallin proteins, the dominant structural components of the vertebrate eye lens.

Other enriched terms for the BSM gene set include the BP GO term "annotation response to gamma radiation" (3 genes, 10.7 fold enrichment), and the disease annotation "breast neoplasms" with three genes (*BRCA2*, *IL17RB*, and *NR1I2*) (20.6 fold enrichment). Another gene not included in this annotation but strongly implicated in breast cancer is *BCAR3*, which is ranked 11th in the BSM gene set. Other notable BSM genes include *CYBRD1*, which is highly expressed in the duodenal brush border membrane and thought to play a physiological role in dietary iron absorption; *BMP3*, which suppresses osteoblast differentiation and negatively regulates bone density by modulating the availability of the TGFB receptor; and *OTOL1*, a collagen-like protein that provides a scaffold for otoconia, crystalline structures of the inner ear involved in the perception of gravity. Finally, a related gene found in the BM gene set, *NOXO1*, is required for the biogenesis of otoconia.

**6) Extended results: Across-capuchins (H3a) positive selection results**

Lists of all enriched annotated terms and GO clusters including annotation category, term description and ID, gene counts and hits, and statistical results such as EASE score and fold enrichment, for the BM gene set enrichment analysis for across-capuchins/Cebinae (H3a) are found in Tables S35 and 36.

6.1 Neurotransmission

For across-capuchins, the BM gene set is enriched for genes related to the vesicle fusion with the top GO cluster (enrichment score 2.07) containing five BP GO terms related to the fusion of organelles, membranes, and vesicles (9 to 13 genes, 2.5 to 3 fold enrichment). Nine of the genes in this cluster are common to all GO terms including six synaptotagmin (*SYT3*, *SYT11*, *SYT14*) and synaptotagmin-like (*SYTL1*, *SYTL3*, and *SYTL5*) genes encoding C-type tandem C2 proteins known to play important roles in regulated exocytosis, neurotransmitter release, and hormone secretion. *SYT3* encodes a calcium sensor involved in the calcium-dependent exocytosis of secretory vesicles through calcium, phospholipid, and SNARE-complex binding to the C2 domain. It is abundantly expressed in all brain regions with expression increasing in parallel with synaptogenesis during postnatal development (of the mouse brain), and also plays a role in dendrite formation by melanocytes. *SYT11* does not bind calcium, phospholipids, or SNARE proteins, but plays an important role in dopamine transmission by regulating endocytosis and vesicle-recycling, and forms an essential component of a neuronal vesicular trafficking pathway that differs from the synaptic vesicle trafficking pathway but is crucial for development and synaptic plasticity (*34*, *164*). *SYT11* is implicated in schizophrenia and late-onset Parkinson’s disease. Like *SYT11*, *SYT14* encodes another calcium-independent synaptotagmin that likely mediates membrane trafficking in synaptic transmission. Mutations in *SYT14* cause a form of the cerebellar disorder spinocerebellar ataxia, and translocation of the gene is associated with neurodevelopmental abnormalities.

Three genes encode synaptotagmin-like proteins (*SYTL1*, *SYTL3,* and *SYTL5*) that likely function as Rab effector proteins with a role in Rab27-dependent vesicle trafficking. *SYTL1* and *SYTL3* are associated with Griscelli syndrome type 1, which is characterised by dilution of pigment in the skin and hair, and neurological impairment including delayed motor development and intellectual disability. Of the other three genes common to all GO terms in this cluster, two are related to SNARE complexes; *YKT6* encodes a SNARE recognition molecule implicated in vesicular transport, docking, and fusion between secretory compartments; and *TSNARE1*, a vertebrate-specific gene of unknown function but likely binds SNARE proteins, playing a role in synaptic vesicle exocytosis, and is implicated in schizophrenia. The final gene common to all GO terms in this cluster is *VPS41*, which plays a role in vesicle-mediated protein trafficking to lysosomal compartments.

The second cluster for the BM gene set (enrichment score 1.69) is related to the first with five GO terms; “calcium ion-regulated exocytosis of neurotransmitter”, “synaptic vesicle exocytosis”, and the MF terms “clathrin binding”, “calcium-dependent phospholipid binding”, and “syntaxin binding” (6 to 7 genes, 2.8 to 5.7 fold enrichment). All these GO terms also contain the six SYT/SYTL genes, with several also including *CPLX3*, which regulates SNARE complex-mediated synaptic vesicle fusion.

Other related genes in the BM gene set not found in these GO terms/clusters include *KCNE4*, which encodes a voltage-gated potassium channel which have diverse functions including regulating neurotransmitter release and insulin secretion (see below), and is predominantly expressed in the embryo and adult uterus; *GABRP*, a subunit of the GABA A receptor, a chloride channel that mediates the fastest inhibitory synaptic transmission in the CNS, with this subunit altering the sensitivity of recombinant receptors to modulatory agents such as pregnanolone, and also playing a role in tissue contractility in the uterus; *PDLIM4*, which is involved in regulation of the synaptic AMPA receptor transport in dendritic spines of hippocampal pyramidal neurons; *SHISH8*, which may regulate trafficking and kinetics of AMPA-type glutamate receptor at synapses; *CA11*, a secreted synaptic protein that functions as neurexin (neuronal cell surface proteins) ligands; *KHDRBS1*, which can regulate alternative splicing of some neurexins involved in neurotransmission and synaptic contacts; and *CASK*, which encodes a calcium/calmodulin-dependent serine protein kinase scaffold protein located at synapses in the brain. *CASK* plays a role in synaptic transmembrane protein anchoring and ion channel trafficking, binds to cell-surface proteins including APP and neurexins, and contributes to neural development and regulation of gene expression via interaction with the transcription factor *TBR1*. Mutations in *CASK* are associated with FG syndrome 4, intellectual disability and microcephaly with pontine and cerebellar hypoplasia, and a form of X-linked intellectual disability.

6.2 Hormones, neuropeptides, and behaviour

The fourth and fifth clusters for the across-capuchin BM gene set are related to peptide and hormone secretion. The fifth cluster contains three BP GO terms (enrichment score 1.54) describing the negative regulation of hormone/peptide secretion (5 to 6 genes, 3.4 to 4.4 fold enrichment). All of the genes in this cluster are also found in the fourth cluster, which contains nine BP GO terms (enrichment score 1.55) also related to hormone/peptide secretion such as “insulin secretion”, “hormone secretion”, “peptide secretion”, the regulation of these processes, and “hormone transport” (10 to 16 genes, 1.9 to 2.7 fold enrichment). The 16 genes in these GO clusters include *LEP*, *GHRL*, *TRH*, *PAX8*, *CGA*, *HMGN3*, *CRH*, *NPFF*, *NPVF*, *CPLX3*, and *TMF1*, with several of these genes overlapping with the results for ancestral Cebinae. There are numerous other related BP GO annotations including the 3rd most enriched BP term “gonadotropin secretion” (5 genes, 11.5 fold enrichment), “luteinising hormone secretion” (4 genes, 13.4 fold enrichment), “endocrine hormone secretion” (5 genes, 4.9 fold enrichment), “endocrine process” (7 genes, 3.1 fold enrichment), “regulation of hormone levels” (24 genes, 1.9 fold enrichment), “hormone metabolic process” (11 genes, 2.2 fold enrichment), “regulation of gonadotropin secretion” (3 genes, 10 fold enrichment), “positive regulation of insulin secretion” and “positive regulation of hormone secretion” (6 and 8 genes, 3.2 and 2.5 fold enrichment), “negative regulation of secretion by cell” (10 genes, 2.2 fold enrichment), and “positive regulation of insulin receptor signalling pathway” (3 genes, 8.5 fold enrichment).

Other related enriched terms include the MF GO term “hormone activity” (10 genes, 3 fold enrichment), and the UP keywords “hormone” (7 genes, 2.8 fold enrichment) and “amidation” (6 genes, 5 fold enrichment); amidation refers to a post-translational modification which is essential to the activity of many neuropeptides and hormones. Some other genes found across these enriched terms that are not in the two hormone clusters include *PRLH*, *FSHB*, *CHST9*, *HSD17B12*, and *SRD5A3*. Another more general BP term is “signal release” (23 genes, 2 fold enrichment), which, in addition to the 16 genes found in the hormone related clusters, also contains the six SYT/SYTL genes in the vesicle fusion cluster and the gene *CASK*.

Many of the genes in these GO terms/clusters relate to hormones produced in the anterior pituitary including the four pituitary glycoprotein hormones (chorionic gonadotropin, luteinising hormone, FSH, and TSH) and prolactin. For example, *CGA*, which encodes the alpha subunit of the four pituitary glycoprotein hormones; *FSHB* encodes the beta subunit of FSH, which is involved in follicle development and spermatogenesis in reproductive organs along with luteinising hormone; *TRH*, which is involved in the secretion of TSH, TH synthesis regulation, and the modulation of hair growth; *PRLH*, prolactin releasing hormone, stimulates prolactin release and regulates the expression of prolactin, which promotes lactation, as well as regulates behaviour, metabolism, and the immune and reproductive systems; and *CHST9*, which participates in biosynthesis of luteinising hormone and TSH by mediating sulphation of their carbohydrate structures. Other genes are related to the thyroid gland including *PAX8*, which is expressed during embryonic development and involved in thyroid follicular cell development and expression of thyroid-specific genes, and is implicated in hypothyroidism and other thyroid-related disorders; and *HMGN3*, which encodes a protein that binds thyroid hormone receptor beta in the presence of TH, impacts insulin and glucagon levels, modulates the expression of pancreatic genes involved in insulin secretion, regulates the expression a glycine transporter that mediates glycine concentration in synaptic junctions in the CNS, and may play a role in ocular development and astrocyte function.

Another notable gene is *CRH*, corticotropin releasing hormone, a major regulator of homeostasis, mediating the autonomic, behavioural, and neuroendocrine responses to stress. CRH is implicated in depression, some forms of epilepsy, and AD. It is also highly expressed in the placenta where it serves as a marker determining the length of gestation, with a rapid increase in circulating levels occurring at the onset of parturition suggesting it may act as a trigger.

Some interesting feeding behaviour related genes in these terms include *GHRL* and *LEP*. *GHRL* encodes a preproprotein that is cleaved into two peptides, ghrelin and obestatin. Ghrelin is a powerful appetite stimulant, plays an important role in energy homeostasis, and is implicated in the regulation of multiple processes including hunger, reward perception, gastric acid secretion, gastrointestinal motility, and pancreatic glucose-stimulated insulin secretion. Obestatin may play an opposing role to ghrelin by promoting satiety and reducing food intake. *LEP*, leptin, is a key player in the regulation of energy balance and body weight control. Leptin activates downstream signalling pathways that inhibit feeding and promote energy expenditure, has several endocrine functions, and is involved in the regulation of immune and inflammatory responses, haematopoiesis, angiogenesis, reproduction, bone formation, and wound healing.

Other genes in these GO terms include *TMF1*, a potential coactivator of the androgen receptor, and *SRD5A3*, which is involved in the production of androgen 5-alpha-dihydrotestosterone from testosterone, and maintenance of the androgen-androgen receptor activation pathway. Other related genes not in these GO terms/clusters include *GPR39*, a member of the ghrelin receptor family that is involved in regulation of body weight, gastrointestinal mobility, hormone secretion, and cell death; *NPFFR1*, a receptor for NPAF and NPFF neuropeptides implicated in hormonal modulation, regulation of food intake, thermoregulation, and nociception; *SCGB3A2*, a secreted lung surfactant protein and a downstream target of thyroid transcription factor that may inhibit production of FSH and luteinising hormone in the pituitary; and two genes encoding serotonin receptors, *HTR3B* and *HTR1F*.

In relation to the above, among the enriched BP GO terms are several related to behaviour including “feeding behaviour” (8 genes, 2.9 fold enrichment), “regulation of behaviour” (6 genes, 3.5 fold enrichment), and “negative regulation of behaviour” (3 genes, 8.5 fold enrichment). Several of these genes are included in the hormone GO clusters/terms (*LEP*, *PRLH*, *TRH*, *GHRL*). Other genes in these GO terms include *BSX*, brain specific homeobox, encoding a DNA binding protein that functions as transcriptional activator, required for normal postnatal growth and nursing, and is an essential factor for neuropeptide Y and agouti-related neuropeptide function, which together act to increase appetite and decrease metabolism and energy expenditure; *AHI1*, important for cortical and cerebellar development; and *GALR3*, which encodes a receptor for the neuropeptide galanin which is widely distributed in the central and peripheral nervous systems and the endocrine system, and modulates a variety of physiologic processes including cognition/memory, sensory/pain processing, hormone secretion, and feeding behaviour.

#### 6.3 Circadian rhythms

As in the ancestral Cebinae BM gene set, the BM gene set for across-capuchins is enriched for genes related to circadian rhythms, with “circadian rhythm” found among the enriched BP GO terms with 12 genes (2.6 fold enrichment), five of which are also found in the ancestral Cebinae BM gene set and annotated by the “circadian rhythm” GO term (*BHLHE41*, *HNRNPD*, *METTL3*, *PER3*, *PROK2*). The seven genes not found for ancestral Cebinae are *KLF9*, *RELB*, *NRIP1*, *USP2*, *CRH*, *LEP*, and *GHRL*. Nine of these genes (except the latter three hormone related genes) are found in the enriched UP keyword “biological rhythms” (2.7 fold enrichment). *RELB*,

*NRIP1*, and *USP2* play a role in the regulation of the core circadian clock and clock-controlled genes, and *KLF9* is as an epidermal circadian transcription factor regulating keratinocyte proliferation.

6.4 Mitochondrion

The signal on the mitochondrion for the ancestral Cebinae lineage is also found more generally across-capuchins. “Mitochondrion” appears as an enriched CC GO term (64 genes, 1.4 fold enrichment), along with other mitochondrion-related CC terms including “mitochondrial matrix” and “mitochondrial part” (22 and 41 genes, 1.9 and 1.5 fold enrichment). Similarly, among enriched UP keywords are the terms “mitochondrion” (47 genes, 1.6 fold enrichment) and “transit peptide” (29 genes, 2 fold enrichment). Two of the genes in the vesicle fusion cluster described above relate to mitochondrial fission and fusion; *MIEF1*, which regulates mitochondrial fission, and *MIGA1*, which regulates mitochondrial fusion.

The second cluster of GO terms with an enrichment score of 1.56 contains five BP GO terms related to the translation of mitochondrial loci: “mitochondrial translational elongation”, “mitochondrial translational termination”, “mitochondrial translation”, “translational termination”, and “translational elongation” (7 to 9 genes, 2.4 to 3.5 fold enrichment). Four of the genes common to all these GO terms are nuclear-encoded mitochondrial ribosomal proteins which form mitochondrial ribosomes (mitoribosomes). These genes are *MRPL9*, *MRPL37*, and *MRPL44*, which encode proteins that form the large 39S subunit of mitoribosomes, and *MRPS28* which encodes a small 28S subunit protein. Mitoribosomes are comprised of a small 28S subunit and a large 39S subunit, and function in protein synthesis within the mitochondrion. Other genes in this GO cluster are *PTCD3*, which encodes a mitochondrial RNA-binding protein that plays a role in mitochondrial translation, and *GFM1*, which encodes a mitochondrial translation elongation factor that catalyses the GTP-dependent ribosomal translocation step. Similar terms related to mitochondrial translation are found as three enriched Reactome pathways ($1^{st}$, $2^{nd}$, and $3^{rd}$; 3.5 to 4 fold enrichment), with seven or eight of the same genes found in the GO cluster. Another related gene found in the BM gene set but not included in this cluster is *ENDOG*, a nuclear encoded endonuclease that is localised in the mitochondrion and plays a role in initiating replication of mitochondrial DNA.

6.5 Other brain and neuronal related

A particularly interesting gene is *CERS*, a ceramide synthase enzyme that catalyses the synthesis of ceramide, the hydrophobic structure of sphingolipids, specifically 18-carbon (C18) ceramide in brain neurons. Elevated expression of this gene has been associated with increased longevity, and decreased expression with myoclonus epilepsy with dementia in humans. Another is *SRPX2*, which promotes synapse formation and is thought to play a role in the development of the perisylvian language region, critical for language and cognitive development, with mutations in this gene causing bilateral perisylvian polymicrogyria, rolandic epilepsy, speech dyspraxia, and cognitive disability. *KIDINS220* encodes a transmembrane protein expressed in the nervous system where it controls neuronal cell survival, differentiation, neurite outgrowth, and synaptic plasticity, and may play a role in axon guidance during neural development and regeneration. It serves as a scaffold that mediates crosstalk between intracellular signalling pathways, and is implicated in various neuropsychiatric disorders and neurodegenerative diseases including AD.

The top ranked gene in the BM gene set is *TTLL1*, which encodes a catalytic subunit of the neuronal tubulin polyglutamylase complex that polyglutamylates alpha subunits of tubulin in the brain. Tubulin proteins form microtubules, which are essential for generation, migration, and differentiation of neurons, with glutamylation being the most prevalent tubulin post-translational modification. Other brain development and neuronal related genes include *EFNA4*, a GPI-bound ligand for ephrin receptors, which are crucial for migration, repulsion, and adhesion during neuronal development; *STMN2*, a stathmin protein that functions in microtubule dynamics, playing a regulatory role in neuronal growth, particularly controlling neurite length in cortical neurons, and also involved in brain development; *C12orf57*, required for the development of the corpus callosum; *AHI1*, involved in vesicle trafficking, neuronal differentiation, and the formation of primary cilia, and it may play a crucial role in ciliary signalling during cerebellum embryonic development as a positive modulator of Wnt signalling; *SSPO*, involved in the modulation of neuronal aggregation and the development of the CNS; *DPF3*, a member of the neuron-specific chromatin remodelling complex, the post-mitotic chromatin remodelling mechanism related to the switch/transition from proliferating neural stem/progenitor cells to committed neurons; *VSTM5*, which plays several important roles including modulating the position and complexity of central neurons, the formation of neuronal dendrites, regulating synapse formation, and regulation of neuronal morphogenesis and migration during cortical development in the brain; and *SZT2*, which is expressed predominantly in the parietal and frontal cortex of the brain, as well as in dorsal root ganglia, localises to the peroxisome, and is implicated in resistance to oxidative stress.

6.6 Branched chain amino acids and metabolic processes

There are signatures in the BM gene set related to branched chain amino acids (BCAAs). The first and second most enriched BP GO terms are "leucine catabolic process" and "leucine metabolic process", both with the same four genes (29.5 and 21 fold enrichment, respectively). Other similar enriched GO terms describe BCAA catabolic and metabolic processes (4 genes, 6.7 to 7.8 fold enrichment). The genes in these GO terms are *AUH*, *MCCC1*, *MCCC2*, *HMGCL*, and *ACADSB*. Other related enriched terms include the KEGG pathway "valine, leucine and isoleucine degradation" (8 genes, 7.7 fold enrichment), and the Reactome "BCAA catabolism" (4 genes, 9 fold enrichment). BCAAs (leucine, isoleucine, and valine) are all essential amino acids required in the diet, found in protein-rich food sources such as eggs and meat, and are major constituents of muscle protein.

Two genes, *FLAD1* and *RFK*, encompass the enriched KEGG pathway "riboflavin metabolism" (30 fold enrichment). These genes, along with *VCP*, also form the enriched BP GO term "flavin-containing compound metabolic process" (18.4 fold enrichment). Riboflavin is a B vitamin involved in many processes in the body and necessary for normal cell growth and function. It is found in certain foods such as meat, eggs, and nuts. Similarly, three genes related to biotin consumption are found in the BM gene set and encompass the enriched Reactome pathway "biotin transport and metabolism" (11.6 fold enrichment): *MCCC1*, *MCCC2*, and *SLC5A6*. Biotin, another B vitamin also known as vitamin H, is an essential nutrient that is involved in the conversion of food to energy, and is important for embryonic growth. It is found in egg yolk, organ meats, nuts, as well as some grains and other plant-based sources.

Other enriched metabolic process related terms include BP GO terms describing glycoprotein metabolic and biosynthetic processes, nucleobase-containing compound, carboxylic acid, and aromatic compound catabolic processes, as well as "fatty acid metabolic process", "fatty acid oxidation", and "lipid oxidation" (7 to 19 genes, 1.7 to 2.7 fold enrichment), the MF GO term "sulphur compound binding" (13 genes, 2 fold enrichment), and the KEGG pathway "metabolic pathways" (40 genes, 1.5 fold enrichment).

6.7) Other

An interesting enriched UP keyword is “deafness” (11 genes, 2.1 fold enrichment) including the genes *GRXCR2*, which could play a role in maintaining cochlear stereocilia bundles involved in sound detection, and *TMPRSS3*, a serine protease that plays a role in hearing, possibly acting as a permissive factor for cochlear hair cell survival and activation, and required for saccular hair cell survival. *TMPRSS3* was first identified through its association with congenital and childhood onset deafness. Some enriched developmental related BP terms include “connective tissue development” (13 genes, 2 fold enrichment), “cartilage morphogenesis” (3 genes, 9.2 fold enrichment), and “cardiac ventricle development” (8 genes 2.5 fold enrichment). Some other interesting developmental genes include three Hox genes, *HOXD9*, *HOXD1*, and *HOXB6*; *DKK3*, which play an important role embryonic development through the inhibition of Wnt regulated processes, and is implicated in bone formation/disease and AD; and *SP7*, a bone specific transcription factor required for osteoblast differentiation and bone formation. Other interesting genes include the olfactory receptor *TAAR5*, which is specific for trimethylamine, a trace amine and bacterial metabolite found in some animal odours, and associated with bad breath and spoiled food for humans; *UROC1*, which encodes an enzyme involved in histidine catabolism and is known to protect the skin from ultraviolet rays; and *ACBD3*, involved in hormone-induced steroid biosynthesis in testicular Leydig cells.

Other enriched terms for the BM gene set include the BP terms “protein hexamerisation” (4 genes, 18.4 fold enrichment), “cellular protein complex disassembly” (11 genes, 2.1 fold enrichment), “positive regulation of transcription from RNA polymerase II promoter transcription factor activity” (40 genes, 1.4 fold enrichment) and “RNA polymerase II core promoter proximal region sequence-specific binding” (18 genes, 1.9 fold enrichment); the UP keywords “activator” (27 genes, 1.5 fold enrichment) and “DNA replication” (8 genes, 3.2 fold enrichment); the Reactome pathways “antigen processing: ubiquitination & proteasome degradation” (14 genes, 1.9 fold enrichment) and “peptide ligand-binding receptors” (7 genes, 2.8 fold enrichment); and the enriched disease terms “acquired immunodeficiency syndrome | disease progression” (37 genes, 1.5 fold enrichment), “triglycerides” (21 genes, 1.7 fold enrichment), and “lipids” (10 genes, 2.2 fold enrichment).

### 7) Extended results: Ancestral Cebidae (H4) positive selection results

Lists of all enriched annotated terms and GO clusters including annotation category, term description and ID, gene counts and hits, and statistical results such as EASE score and fold enrichment, for the BM and BSM gene set enrichment analyses for ancestral Cebidae (H4) are found in Tables S37–40.

#### 7.1 Cilium

The ancestral Cebidae BM gene set is strongly enriched for cilium-related genes. The top GO cluster contains five terms (enrichment score 3.37): "cilium morphogenesis" (also the most significantly enriched BP term), "cilium assembly", "cilium organisation", "cell projection assembly", and "cellular component assembly involved in morphogenesis" (12 to 16 genes, 2.7 to 4.1 fold enrichment). Some of the genes in this cluster include *WDR35* ($2^{nd}$ rank in the BM gene set), *DZIP1* ($8^{th}$ rank in the BM gene set), *SCLT1*, *TMEM67*, *KIAA0586*, *CLUAP1*, and *IFT122* (also found in the BSM gene set), all of which are involved in ciliogenesis with several encoding components of, or associated with, the intraflagellar transport machinery (IFT) and implicated in forms of Joubert's syndrome. *IFT122* is involved in cilia formation during neuronal patterning; *TMEM67* is required for ciliary structure/function and forms part of the tectonic-like complex required for tissue-specific ciliogenesis; and *KIF24* acts as a negative regulator of ciliogenesis. Others genes in this cluster are central to the function of motile cilia including *DRC1*, which encodes a component of the nexin-dynein regulatory complex, a key regulator of ciliary/flagellar motility; *TEKT3*, required for normal sperm motility; and *ZMYND10*, *CCDC151*, and *LRRC6*, each thought to play a role in dynein arm assembly, essential for axoneme building for cilia motility. These motility-related genes are implicated in ciliary dyskinesia disorders and comprise the enriched UP keyword "primary ciliary dyskinesia" (4 genes, 9.3 fold enrichment). A second cluster contains three BP GO terms related to the axoneme (enrichment score 1.99): "axonemal dynein complex assembly", "axoneme assembly", and "microtubule bundle formation"; 5 genes, 4.5 to 11 fold enrichment). Another enriched related BP GO term not found in these clusters is "outer dynein arm assembly" with three of the same genes (14 fold enrichment).

The single most enriched GO annotation is the CC GO term "cilium" and the second most enriched CC GO term is "ciliary part" (22 and 15 genes, 3 fold enrichment). Other related enriched CC GO terms include "motile cilium" (8 genes, 4.3 fold enrichment), and "ciliary tip"

(4 genes, 6.4 fold enrichment). Genes include *ADCY10*, which plays a critical role in mammalian spermatogenesis by producing cAMP and inducing sperm maturation prior to fertilisation, and is involved in ciliary beat regulation; *DRC7*, which encodes a component of the nexin-dynein regulatory complex (as *DRC1* above); and *CATSPER3* (also found in the BSM gene set) (see below). Other related signatures including many of the same genes as in the cilium-related GO cluster can be found as enriched UP keywords "cilium", "ciliopathy", and "cilium biogenesis/degradation" (8 to 13 genes, 4 to 4.7 fold enrichment), and enriched Reactome pathways "intraflagellar transport" and "anchoring of the basal body to the plasma membrane" (4 and 5 genes, 5.5 and 3.8 fold enrichment).

Given the strong signatures of accelerated evolution in cilia-related genes, it is unsurprising that there are overlapping signatures related to microtubules including the BP GO terms "microtubule-based process", "microtubule-based movement", "microtubule cytoskeleton organisation", as well as related CC terms including "microtubule cytoskeleton", "microtubule organising centre", "centrosome", "centriole", "cytoskeleton", and "cell projection", and the UP keywords "cytoskeleton" and "cell projection" (8 to 42 genes, 1.4 to 5 fold enrichment). There are also similar signatures in the BSM gene set such as the enriched CC GO terms "microtubule organising centre", "cytoskeleton", "centrosome", "microtubule cytoskeleton", and "cytoskeletal part" (8 to 20 genes, 1.8 to 3.1 fold enrichment), and the UP keyword "cell projection" (9 genes, 2.5 fold enrichment).

#### 7.2 Sperm development and reproduction

Both the BM and BSM gene sets for ancestral Cebidae are enriched for sperm development and reproduction related genes, with some overlap with the cilium-related clusters and GO terms. For the BM gene set, there is a cluster of three BP GO terms (enrichment score 1.37): "spermatid development", "spermatid differentiation", and "germ cell development" (6 to 8 genes, 2.4 to 3.2 fold enrichment). Some other genes in these GO terms include *PLD6*, which encodes an endonuclease that plays a critical role in piRNA biogenesis during spermatogenesis; *TOPAZ1*, which is required for progression to post-meiotic stages of spermatocyte development and thereby important for normal spermatogenesis and male fertility; and *CATSPER3*, a voltage-gated calcium channel that plays a central role in calcium-dependent physiological responses essential for successful fertilisation, such as sperm hyperactivation, acrosome reaction and chemotaxis towards the oocyte. Similarly, for the BM gene set, the enriched UP keyword "spermatogenesis" (7 genes, 2.7 fold enrichment) contains many of the same genes as these GO terms as well as *TDRD12*, which encodes an ATP-binding RNA helicase required during spermatogenesis to repress transposable elements and prevent their mobilisation, which is essential for the germline integrity, and *CALR3* (11$^{th}$ rank in BM gene set, also found in BSM gene set), which is required for sperm fertility.

Other related enriched BP GO terms in the BM gene set are "male meiosis" and "male meiosis I" (4 and 3 genes, 6.8 and 11.7 fold enrichment). These genes include *DMC1* and *TDRD12*, both found in the spermatid GO terms, as well as *MEIOB* (also found in the BSM gene set), which encodes a single-stranded DNA binding protein required for homologous recombination in meiosis I and implicated in male infertility, and *MOV10L1* (13$^{th}$ rank in BM gene set), which encodes another ATP-dependent RNA helicase required during spermatogenesis to prevent mobilisation of transposable elements. Similarly, the enriched BP GO terms "piRNA metabolic process" and "DNA methylation involved in gamete generation" (14 and 11.7 fold enrichment) are comprised of three aforementioned genes *PLD6*, *TDRD12*, and *MOV10L1*. The piRNA pathway is involved in the epigenetic and post-transcriptional silencing of transposable elements, with a role in safeguarding genome integrity and fertility. Several of the genes mentioned above are also found in the BP term "cellular process involved in reproduction in multicellular organism" (9 genes, 2.4 fold enrichment). This GO term also includes *ZMYND15*, *RNF17*, and *INHBA*, the latter encoding a subunit of activin and inhibin protein complexes that activate and inhibit, respectively, FSH secretion from the pituitary gland, as well as playing a role in eye, tooth, and testis development.

Similar signatures regarding spermatogenesis, male meiosis, and reproduction are also found in the BSM gene set with some overlapping genes found in both gene sets. For the BSM gene set, the top cluster (enrichment score 1.56) contains BP GO terms such as "spermatogenesis", "male gamete generation", "sexual reproduction", and other similar GO terms (8 to 10 genes, 2.1 to 3.1 fold enrichment). Three of these genes are also found in the BM gene set (*MEIOB*, *CALR3*, *CATSPER3*), while the other eight genes include *CATSPERD* (4$^{th}$ rank in BSM gene set), which is involved in sperm cell hyperactivation (needed for sperm motility and thus, sperm preparation for fertilisation); *SPATA5* (3$^{rd}$ rank in BSM gene set), which may be involved in morphological and functional mitochondrial transformations during spermatogenesis; and *TEX15*, required during spermatogenesis for normal chromosome synapsis and meiotic recombination in germ cells. A particularly interesting gene found in the reproduction related GO terms in this cluster is *DEFB126*, which encodes an atypical beta-defensin involved in several aspects of sperm function including facilitating sperm transport in the female reproductive tract, contributing to sperm protection against immunodetection, binding sperm to oviductal epithelial cells to form a sperm reservoir until ovulation, and release from the sperm surface during capacitation and ovulation allowing sperm to bind to the zona pellucida of the oocyte. Some of the GO terms in this cluster are also among the top ranked individual BP GO terms for the BSM gene set.

Other related enriched GO terms for the BSM gene set include “fertilisation”, which is the second ranked individual GO term (6 genes, 7.1 fold enrichment), “male meiosis” (3 genes, 14.3 fold enrichment), and “spermatid development” and “spermatid differentiation” (4 genes, 6 and 5.8 fold enrichment). Enriched CC GO terms include “motile cilium” (4 genes, 5.6 fold enrichment), and “CatSper complex” (40.5 fold enrichment) with two genes (*CATSPER3, CATSPERD*), which also comprise the enriched Reactome pathway “sperm motility and taxes” (42.9 fold enrichment). Further signals include the enriched UP keywords “flagellum” (3 genes, 19.8 fold enrichment) and “spermatogenesis” (5 genes, 5.1 fold enrichment). All except one of the genes in these enriched GO terms and UP keywords are also found in the spermatogenesis/reproduction cluster. Another related BP term in the BSM gene set is “cell recognition” (4 genes, 5 fold enrichment), with three genes overlapping with these terms (*CATSPER3*, *CATSPERD*, *NECTIN3*).

Another interesting gene in the BSM gene set is *PRSS8*, a serine protease highly expressed in prostate epithelia and is one of several proteolytic enzymes found in seminal fluid. Other BM genes include *EFCAB9*, a pH-dependent calcium ion sensor required to activate the CatSper complex involved in sperm cell hyperactivation; *PRSS55,* another serine protease involved in sperm migration and sperm-egg interaction; *HYAL3,* which facilitates sperm penetration into the layer of cumulus cells surrounding the egg by digesting hyaluronic acid, and is involved in follicular atresia and in induction of the acrosome reaction in sperm; and *TTLL9,* which mediates tubulin polyglutamylation thereby playing a role in the establishment of microtubule heterogeneity in sperm flagella. Together the concurrent signals in both gene sets, with several highly ranked genes and some overlapping genes across both sets, suggests strong signatures of selection related to sperm development and motility for the ancestral Cebidae branch.

#### 7.3 Immune system

There are various signatures related to immune system processes across both gene sets. The BM gene set contains a cluster of three BP GO terms (enrichment score 1.66) related to I-kappaB kinase/NF-kappaB signalling and regulation (8 to 9 genes, 2.5 to 3.1 fold enrichment). These genes include two serine/threonine protein kinases (*MAP3K7*, and *RIPK2*, also found in the BSM gene set), as well as *NKIRAS2* and SQSTM1, which all play essential roles in the activation or regulation of NF-kappaB and the modulation of immune responses (innate and adaptive). Another cluster in the BM gene set (enrichment score 1.58) includes three BP GO terms related to the cellular response to endotoxins/lipopolysaccharides (7 genes, 2.8 to 3.3 fold enrichment). Other enriched immune-related individual GO terms in the BM gene set are “cellular response to interleukin-4” (4 genes, 9.7 fold enrichment) and “macrophage fusion” (2 genes, 46.6 fold enrichment).

The BSM gene set is enriched for T cell and adaptive immunity related genes. Among the most enriched GO annotations is the CC GO term “T cell receptor complex” (3 genes, 28.8 fold enrichment). Enriched BP terms describing the positive regulation of T cell, lymphocyte, mononuclear cell, and leukocyte proliferation (4 genes, 5.6 to 8 fold enrichment) form the second GO cluster (enrichment score 1.55). Other enriched GO terms include “adaptive immune response” (7 genes, 3.3 fold enrichment) and “acute inflammatory response” (3 genes, 5.7 fold enrichment), and the CC term “membrane attack complex”, which is also found as an enriched UP keyword (2 genes, 52.1 and 56.5 fold enrichment). Further overlapping signals include the enriched UP keywords “adaptive immunity” and “immunity” (5 and 7 genes, 5.7 and 2.8 fold enrichment, respectively). The genes in these enriched terms are *CD4*, *CD6* (10$^{\text{th}}$ ranked gene in the BSM gene set), *CD8B*, *C7*, *C8A* (also found in the BM gene set), *ICOSLG*, *RIPK2*, and *OSMR*.

#### 7.4 Brain/neuronal development and plasticity

Among the enriched individual GO annotations for the BM gene set is the CC GO term “growth cone” with seven interesting genes (3.5 fold enrichment). Growth cone refers to the migrating motile tip of a growing neuron projection. These genes include *ADCY10*, which encodes a soluble adenylyl cyclase that catalyses the formation of the signalling molecule cAMP, enhancing neurite outgrowth and facilitating regeneration after injury, thus playing a key role in neuronal survival and axon growth. *CTTN* contributes to the organisation of the actin cytoskeleton and plays a role in the regulation of neuron morphology, axon growth, and formation of neuronal growth cones. *CDKL5* encodes a serine-threonine kinase that is highly expressed in the brain with the highest concentrations during peri- and post-natal stages of the rapid development of the nervous system (particularly in the cerebral cortex and the hippocampus). Although the exact molecular function is unknown, it is involved in proliferation, neuronal migration, neuronal formation, and neuronal growth, as well as in the development and functioning of synapses in brain maturation. Mutations in this gene cause a rare developmental epileptic encephalopathy characterised by early-onset, intractable epilepsy and neurodevelopmental delay impacting cognitive, motor, speech, and visual function (*165*). *STMN4* encodes a stathmin tubulin-binding phosphoprotein highly expressed in the nervous system, particularly during brain development. Stathmins are essential regulators of neuronal differentiation (at the various developmental stages) and plasticity of the nervous system, and their expression is altered in numerous neurodegenerative diseases (*166*). *SIGMAR1* (5th ranked gene in the BM gene set) encodes a receptor protein involved in learning processes, memory, and mood alteration, with a potential role in modulating neurotransmitter release through the regulation of ion channels. *SIGMAR1* is implicated in forms of ALS, and ALS is also among the enriched disease annotations for the BM gene set (5 genes, 5.6 fold enrichment). The final genes in the growth cone CC GO term are *GPRIN1* (also found in the BSM gene set), which may be involved in neurite outgrowth, and *PTPRO* (4th ranked gene in the BM gene set), a cell adhesion molecule that has been shown to induce the formation of artificial synapse clusters; synaptic cell adhesion molecules play essential roles in initiating the formation of synapses, critical for brain function (*167*).

Other interesting BSM genes include *RAB18* (ranked 10th in the BSM gene set and also in the BM gene set), which plays a key role in eye and brain development as well as neurodegeneration; *ADGRG6* (also in the BM gene set), which regulates neural, cardiac, and ear development, and is essential for normal myelination of axons and differentiation of promyelinating Schwann cells; *NTF4*, a member of the neurotrophic factor family that control survival and differentiation of mammalian neurons; *LRIT3*, which plays an important role in synapse formation and synaptic transmission between cone photoreceptor cells and retinal bipolar cells, and is associated with night blindness; *ARL6IP5*, which regulates intracellular concentrations of taurine and glutamate, and implicated in neuronal ceroid lipofuscinosis; and *ASXL2*, which belongs to a family of epigenetic regulators that bind various histone-modifying enzymes and maintain repression of homeotic genes during development, with this gene playing an important role in neurodevelopment, cardiac function, adipogenesis, and osteoclastogenesis. A similar gene from the same family, *ASXL3*, is found in the BM gene set.

Other interesting BM genes include *TRIM44*, which is a negative regulator of *PAX6* expression, thought to play a role in neuronal differentiation and maturation; *DUSP15*, which may play a role in the regulation of oligodendrocyte differentiation and myelin formation; and *RNF112*, which encodes an E3 ubiquitin ligase that plays an important role in neuronal differentiation during brain development, as well as in the protection of the nervous tissue cells from oxidative stress-induced damage, and regulating dendritic spine density and synaptic neurotransmission.

#### 7.5 Neuromodulation & behaviour

Among the enriched Reactome pathways for the BSM gene set is “orexin and neuropeptides FF and QRFP bind to their respective receptors” with two genes (48.3 fold enrichment), which is related to the regulation of sleep and appetite. One of these genes, *NPFFR2*, the top ranked gene in the BSM gene set, encodes a GPCR activated by the neuropeptides NPAF and NPFF, and implicated in hormonal modulation, regulation of food intake, thermoregulation, and nociception. *HCRT* encodes a hypothalamic neuropeptide precursor that gives rise to two mature neuropeptides, orexin A and orexin B, which function in the regulation of sleep and arousal, and may also play a role in feeding behaviour, metabolism, and homeostasis. Another interesting gene in the BSM gene set is *DBH*, dopamine beta-hydroxylase, which catalyses the conversion of dopamine to norepinephrine thereby playing a role in the bioavailability of both. Dopamine and norepinephrine are crucial neuromodulators involved in major brain computation processes such as sensory processing, plasticity, memory encoding, learning, mood maintenance, motivation, and concentration, with norepinephrine also functioning as the main neurotransmitter of the sympathetic nervous system (*168*). Mutations in *DBH* are implicated in a range of psychiatric disorders. Another interesting related gene in the BSM gene set is *NMUR1*, a receptor for the neuromedin-U and neuromedin-S neuropeptides.

A particularly interesting BM gene is the serotonin receptor, *HTR1A*, primarily located in limbic brain areas, notably the hypothalamus and cortical areas, playing a role in the response to anxiogenic stimuli, regulation of serotonin release, and regulation of serotonin and dopamine metabolism and levels in the brain, thereby affecting neural activity, mood, and behaviour. Inactivation of this gene in mice leads to increased anxiety and stress responses, and it is implicated in generalised anxiety disorder. Other interesting related genes in the BM gene set include *VIPR2*, a receptor for the neuropeptide vasoactive intestinal peptide that is widely distributed throughout the CNS and involved in smooth muscle relaxation, exocrine and endocrine secretion, and water/ion stasis in lung and intestinal epithelia; *XBP1*, which has many functions including a role in the survival of dopaminergic neurons of the substantia nigra pars compacta; and *RPH3A*, which plays an important role in neurotransmitter release and synaptic vesicle traffic, and involved in the exocytosis of arginine vasopressin hormone.

#### 7.6 Aging

The BSM gene set includes the enriched disease annotation "aging" with 12 genes (2.2 fold enrichment) and six of the same genes are found in the disease annotation "longevity" (2.5 fold enrichment), though the EASE score for "longevity" falls above significance. The "aging" annotation includes diverse genes with a variety of biological functions including some with a role in apoptosis (*TNFRSF10B*, *RIPK2*), metabolism (*MMUT*, *GSTO2*), immunity (*OSMR*, *C7*), DNA repair and genome integrity (*WRN*), and ATP synthesis (*ATP5F1*), among other functions (*CMA1*, *EMP1*, *DBH*). Notably, several of these genes are also found in the BM gene set (including *RIPK2*, *GSTO2*, *WRN*, and *ATP5F1*). *GSTO2* and *C7* are implicated in the age at onset for AD (and Parkinson's for *GSTO2*), and similarly, *ATP5F1* is associated with AD and Huntington's disease. *WRN* plays a major role in genome stability, particularly during DNA replication and telomere metabolism. Mutations in *WRN* are associated with defective telomere maintenance and cause Werner syndrome, which is characterised by rapid onset of cellular senescence, early cancer onset, and premature aging (*50*). Another gene involved in telomere maintenance in the BSM gene set is *NSMCE4A*. Similarly, an enriched disease annotation for the BM gene set is "chromosome aberrations | DNA damage" (3 genes, 9.6 fold enrichment).

Another important gene implicated in aging found in the BSM gene set (ranked 6th) is *SMPD1*, a lysosomal acid sphingomyelinase (ASM), one of the significant sphingolipid-metabolising enzymes which catalyses the conversion of sphingomyelin, a significant component of membranes, into ceramide and phosphocholine. ASM also plays a role in multiple signalling processes, including cell survival, permeability, and proliferation, and is vital in mediating senescence and apoptosis. The expression and activity of ASM changes with age showing marked elevation in the brains of old mice, and recent studies have highlighted the importance of ASM as a critical mediator that contributes to pathologies in aging and age-related neurodegenerative diseases, with ASM viewed as a promising drug target for anti-aging and the treatment of age-related neurodegenerative diseases (*51*). There are also several other genes in the BM gene set related to sphingolipid and ceramide metabolism including the ceramide synthase, *CERS4*; *TEX2*, which prevents toxic ceramide accumulation when cellular ceramide levels increase by facilitating non-vesicular transport of ceramides from the ER to the Golgi complex where they are converted to complex sphingolipids; and *ORMDL2*, a negative regulator of sphingolipid synthesis.

An important gene found in both the BSM (ranked 9th) and BM gene sets is *ADAM10*, a member of the ADAM family of cell surface proteins possessing both adhesion and protease domains; it is an alpha secretase responsible for the cleavage of APP thereby preventing the generation of amyloid beta peptides associated with the development of AD. Other substrates for *ADAM10* include Notch, growth factors, adhesion molecules, ephrins, and their receptors, playing a role in development and neurogenesis via Notch processing. Notably, *TSPAN17*, which regulates *ADAM10* maturation, is also found in both gene sets. Furthermore, another important alpha secretase from the same family, *ADAM9*, is found in the BM gene set (ranked 10th), along with *RASD1*, a small GTPase that interacts with nuclear adaptor protein *FE65*, which interacts with APP. Another interesting related BM gene is *ACMSD*, which plays an important role in preventing the accumulation of quinolinate, a key precursor of NAD, and a potent endogenous excitotoxin of neuronal cells implicated in various neurodegenerative disorders.

#### 7.7 Other

In the BM gene set, the enriched UP keywords “blood coagulation” and “haemostasis” contain the same five genes (both 7.7 fold enrichment): *F2*, *F2R*, *F13B*, *FGB*, and *SERPIND1*. These five genes are also found in the most enriched KEGG pathway “complement and coagulation cascades” (8.1 fold enrichment), along with *C8A*, and the enriched Reactome pathway “common pathway of fibrin clot formation” (20.1 fold enrichment), along with *CD177*. *F2*, which encodes coagulation factor II (thrombin), plays an important role in thrombosis and haemostasis by converting fibrinogen to fibrin during blood clot formation, stimulating platelet aggregation, and activating additional coagulation factors. Thrombin also plays a role in cell proliferation, tissue repair, angiogenesis, and maintaining vascular integrity. *F2R* encodes a thrombin receptor involved in the regulation of thrombotic response. *F13B* encodes a subunit of coagulation factor XIII, the last zymogen to become activated in the blood coagulation cascade, and plays a role in stabilising the fibrin clot. *FGB* encodes the beta component of fibrinogen, a blood-borne glycoprotein cleaved by thrombin to form fibrin following vascular injury. *SERPIND1* encodes a thrombin inhibitor. “Thrombosis, deep vein” is also among the most enriched disease annotation (3 genes, 10 fold enrichment).

The fifth GO cluster in the BM gene set (enrichment score 2.09) includes three BP GO terms related to the specification of left/right symmetry (7 genes, 3.9 to 4.2 fold enrichment). Four of these genes are also found in the cilia-related GO cluster/terms (*CLUAP1*, *CCDC151*, *DRC1*, *LRRC6*) with three other genes: *NKX3-2,* a NK homeobox protein which acts as a negative regulator of chondrocyte maturation, and is required for development of some components of the middle ear; *TBX20*, essential for heart development; and *FOXN4*, which encodes a transcription factor essential for the development of neural tissues particularly the retina, as well as a role in spinal cord neurogenesis, and development of some non-neural tissues including the lung. Four of these genes, along with *IFT122*, also comprise the enriched BP GO term “embryonic heart tube development” (4.5 fold enrichment).

The third GO cluster in the BM gene set (enrichment score 2.71) contains four BP GO terms related to DNA alkylation, methylation, and modification (6 genes, 4.8 to 8.2 fold enrichment). Three of these genes are related to the piRNA pathway discussed in section 7.2 (*PLD6*, *TDRD12*, and *MOV10L1*), and the other three genes are *HEMK1*, *ATF7IP*, which modulates transcription regulation and chromatin formation, and *ATRX*, which is involved in chromatin remodelling and transcriptional regulation. Mutations in *ATRX* are associated with X-linked syndromes exhibiting cognitive disabilities and shown to cause diverse changes in DNA methylation patterns. For the BM gene set, there are several GO terms related to DNA metabolism including “DNA metabolic process” (24 genes, 1.7 fold enrichment), “DNA-dependent DNA replication” and “DNA replication” (7 and 10 genes, 3.4 and 2.4 fold enrichment), “DNA duplex unwinding” and “DNA geometric change” (5 genes, 4.6 and 4.3 fold enrichment), and “mitochondrial DNA metabolic process” (3 genes, 11.7 fold enrichment), as well as the enriched MF terms “nucleoside-triphosphatase activity” and “helicase activity” (20 and 7 genes, 1.7 and 3 fold enrichment), and UP keywords “nucleotide binding” and “helicase” (35 and 7 genes, 1.4 and 3.6 fold enrichment).

Similarly, there is a cluster of five BP GO terms in the BM gene set (enrichment score 1.62) related to nucleoside biosynthesis (6 genes, 3.2 to 4.2 fold enrichment), as well as various enriched ATP-related terms including the BP GO terms “regulation of ATP biosynthetic process” and “protein poly-ADP-ribosylation” (3 and 2 genes, 26.2 and 46.6 fold enrichment), an enriched KEGG pathway “AMPK signalling” (which promotes ATP-producing and inhibits ATP-consuming pathways) (5 genes, 3.8 fold enrichment), and the UP keyword “ATP binding” (28 genes, 1.5 fold enrichment). Further related MF GO terms in the BM gene set include various hydrolase activity terms, as well as “adenosine deaminase activity”, “deaminase activity”, and “pyrophosphatase activity” (3 to 53 genes, 1.4 to 18.4 fold enrichment), some of which form a GO cluster (enrichment score 1.68), and the UP keyword “hydrolase” (35 genes, 1.6 fold enrichment). Another related BP term “cellular amino acid metabolic process” is found in both gene sets (10 and 5 genes, 2.8 and 3.9 fold enrichment). An interesting gene in several of these terms is *PPARGC1A*, a transcriptional coactivator for steroid receptors that greatly increases the transcriptional activity of TH receptor, regulates key mitochondrial genes contributing to adaptive thermogenesis, plays an essential role in metabolic reprogramming in response to dietary availability by coordinating the expression of diverse genes involved in glucose and fatty acid metabolism, and is also involved in the integration of the circadian rhythms and energy metabolism, and required for oscillatory expression of some clock genes.

The second GO cluster for the BM gene set (enrichment score 2.82) contains a large number of genes related to macromolecular and protein complex assembly, biogenesis, and subunit organisation (36 to 51 genes, 1.4 to 1.7 fold enrichment). Other related GO terms include “cellular macromolecular complex assembly” and “cellular protein complex assembly” (24 and 15 genes, 1.7 and 1.8 fold enrichment). Signals related to protein complexes and cellular components underly the even broader GO terms “cellular component assembly” and “cellular component biogenesis” (60 and 65 genes, 1.6 fold enrichment), and “cellular component organisation or biogenesis” (107 genes, 1.2 fold enrichment). Among the enriched individual GO terms for the BM gene set are several CC terms related to the mitochondrion including “mitochondrial matrix” (also enriched for the BSM gene set), “mitochondrial part”, and “mitochondrion” (13 to 35 genes, 1.4 to 2.2 fold enrichment), and “mitochondrial nucleoid” (4 genes, 6.3 fold enrichment). Both gene sets also have the enriched UP keyword “mitochondrion” (26 and 12 genes, 1.7 and 2.1 fold enrichment), with the UP keyword “transit peptide” enriched in the BM gene set (15 genes, 2.1 fold enrichment). For the BM gene set, there are also signatures of accelerated evolution related to the endoplasmic reticulum, found as two enriched CC terms and a UP keyword (23 to 37 genes, 1.6 to 1.7 fold enrichment). Related to the combined signatures on cell organelles is the enriched BP GO term “organelle assembly” (19 genes, 2 fold enrichment), and another enriched cell related BP term is “cell-substrate adhesion” (10 genes, 2.3 fold enrichment).

An enriched BP GO term found in both gene sets is “water-soluble vitamin metabolic process” (5 and 4 genes, 4 and 8.9 fold enrichment) with two of those genes in the enriched BSM Reactome pathway “vitamin C (ascorbate) metabolism” (48.3 fold enrichment). There are also various signatures related to metabolic processes across both gene sets. The fourth cluster (enrichment score 2.49) for the BM gene set includes three BP GO terms related to organic acid metabolism (23 to 24 genes, 1.9 to 2 fold enrichment). Other metabolism related terms in the BM set include the BP GO terms “coenzyme metabolic process” and “carbohydrate derivative biosynthetic process” (11 and 19 genes, 2.4 and 1.8 fold enrichment), and KEGG pathway “metabolic pathways” (22 genes, 1.7 fold enrichment). Similar general signatures can be found for the BSM gene set with the third cluster of three BP GO terms (enrichment score 1.55) also related to organic acid metabolism (10 genes, 2.2 to 2.4 fold enrichment). There are some signatures related to lipids in the BM gene set including the BP term “positive regulation of lipid metabolic process” (6 genes, 3.3 fold enrichment), the CC term “lipid particle” (5 gene, 5.3 fold enrichment), and the UP keyword “lipid metabolism” (12 genes, 2.1 fold enrichment). For the BM gene set, the other MF GO terms are “oestrogen receptor binding” and “manganese ion binding” (4 genes, 7.3 and 5.4 fold enrichment), and enriched disease annotations include “myocardial infarction” (16 genes, 2.2 fold enrichment) and “metabolic syndrome” (7 genes, 3.1 fold enrichment), among others. Other enriched terms in the BSM gene set include the BP terms "protein processing" and "protein maturation" (5 genes, 4.1 and 3.6 fold enrichment), the MF terms "transition metal ion binding" and "zinc ion binding" (14 and 12 genes, 1.8 and 1.9 fold enrichment), the CC GO terms "plasma membrane protein complex", "intrinsic component of plasma membrane", and "secretory granule"  (6 to 16 genes, 1.7 to 3.1 fold enrichment), and the UP keywords "signal" and "glycoprotein" (35 to 36  genes, 1.5 to 1.7 fold enrichment).

Other interesting genes in the BSM gene set include *KEAP1*, which regulates the response to oxidative stress; *KIAA1551* (*RESF1*) plays a role in the regulation of imprinted gene expression; *KLF14*, a transcriptional co-repressor that exhibits imprinted expression from the maternal allele in embryonic and extra-embryonic tissues; *FPGS*, essential for folate homeostasis; and *TRIM25*, which mediates oestrogen action in various target organs.

### 8) Extended results: Squirrel monkeys (*Saimiri*; H5) positive selection results

Lists of all enriched annotated terms and GO clusters including annotation category, term description and ID, gene counts and hits, and statistical results such as EASE score and fold enrichment, for the BM and BSM gene set enrichment analyses for squirrel monkeys (*Saimiri*: H5) are found in Tables S41–44.

#### 8.1 Growth factors

Both the BSM and BM gene sets for *Saimiri* are enriched for signatures related to growth factors. For the BSM gene set, there are multiple similar GO annotations including the MF terms "growth factor activity" (6 genes, 4.6 fold enrichment) and "growth factor receptor binding" (5 genes, 4.8 fold enrichment), and the BP terms "cellular response to growth factor stimulus" and "response to growth factor" (11 genes, 2.1-2.2 fold enrichment), in addition to the enriched UP keyword "growth factor" (6 genes, 5.1 fold enrichment). Genes found among these GO terms include two fibroblast growth factor (FGF) family members which possess broad mitogenic and cell survival activities, and are involved in a variety of biological processes, including embryonic development, cell growth, morphogenesis, and tissue repair. *FGF1* plays an important role in the regulation of cell survival, division, differentiation, and migration, and angiogenesis. *FGF20* is expressed in the normal brain, particularly the cerebellum, and may regulate CNS development and function. These two FGF genes also form the enriched Reactome pathway "FGFR3b ligand binding and activation" (39.3 fold enrichment).

There are three genes from the TGFB superfamily in these GO terms, which play fundamental roles in the regulation of basic biological processes such as growth, development, and immune system function. *TGFB1*, which the protein family was named after, encodes a multifunctional protein that regulates the growth and differentiation of various cell types and is involved in various processes, such as normal development, immune function, and response to neurodegeneration. It plays a role in the activation of other growth factors and in bone remodelling, acting as a potent stimulator of osteoblastic bone formation causing chemotaxis, proliferation, and differentiation in committed osteoblasts. *BMP15* plays a role in oocyte maturation and follicular development. *GDF10*, growth differentiation factor 10, is involved in osteogenesis and adipogenesis. Other genes include *NTF4*, which encodes a neurotrophic factor that controls survival and differentiation of mammalian neurons; *NGFR*, a low affinity nerve growth factor receptor which can bind to the products of *NGF*, *BDNF*, *NTF3*, and *NTF4*, playing an important role in differentiation and survival of specific neuronal populations during development, and mediating cell survival / death of neural cells; *SOX11*, a transcription factor of the SOX family which are involved in the regulation of embryonic development and in the determination of cell fate, with a role in the developing nervous system for *SOX11*; *RGMB*, which encodes a member of the repulsive guidance molecule family that contributes to the patterning of the developing nervous system and acts as a BMP coreceptor with a role in BMP signalling; and *DUSP6*, a dual specificity protein phosphatase with specificity for the ERK family.

Several of the genes in the BSM growth factor related GO terms are also found in the KEGG pathway "PI3K-Akt signalling pathway" (6 genes, 2.8 fold enrichment) including *FGF1*, *FGF20*, *IL6R*, and *NGFR*. The other genes are *OSMR* and *COL4A4*. The PI3K-Akt pathway is a signal transduction pathway promoting metabolism, proliferation, cell survival, growth, and angiogenesis.

The BM gene set contains the enriched GO term "regulation of cellular response to growth factor stimulus" with 11 genes (2.3 fold enrichment) including *BMPER*, which encodes a secreted protein that inhibits BMP function and regulates BMP responsiveness of osteoblasts and chondrocytes. Mutations in *BMPER* are associated with a lethal skeletal disorder in humans called diaphanospondylodysostosis. Other genes include *CD63*, a cell surface protein which mediates cellular signalling cascades that play a role in the regulation of cell development, activation, growth, and motility; *FOLR1*, a member of the folate receptor family that binds folic acid, required for normal embryonic development and cell proliferation, with mutations in this gene associated with neurodegeneration due to cerebral folate transport deficiency; *ILK*, a protein kinase that regulates integrin-mediated signal transduction; and *KCP*, which enhances BMP signalling and inhibits activin-A and TGFB1-mediated signalling pathways.

Other interesting growth factor related genes in the BM gene set not found in these GO terms include *IGF2R* (also in the BSM gene set), which encodes a receptor for insulin-like growth factor (IGF) 2 and has various functions including the activation of TGFB and the degradation of IGF2; *PDGFB*, which encodes a growth factor that plays an essential role in the regulation of embryonic development, cell proliferation, migration, survival, and chemotaxis; several genes associated with epidermal growth factors, *GRB2*, *NRG3*, and *GAREM1*, with the latter also involved in the activation of the MAPK/ERK signalling; and *IGFALS*, which encodes a protein that binds IGFs, increasing their half-life and impacting their vascular localisation. Mutations in *IGFALS* cause acid-labile subunit deficiency characterised by a severe reduction in IGFI and disruption to the overall IGF circulating system, leading to short stature and delayed/slow puberty, and it is also implicated in Laron Syndrome, which is characterised by short stature.

Similarly, other interesting BM genes related to development/growth/body size include *SHOX*, short stature homeobox, which controls fundamental aspects of growth and development, and defects are associated with reduced growth, the short stature phenotype of Turner syndrome, and Leri-Weill dyschondrosteosis, a skeletal dysplasia characterised by shortened limbs and stature; *ING1*, inhibitor of growth family member 1, which encodes a tumour suppressor protein that can induce cell growth arrest and apoptosis; *GRAP2*, encoding an adaptor-like protein involved in protein tyrosine kinase signalling and associated with short stature; and *EIF2AK3*, which encodes a metabolic-stress sensing protein kinase that represses global protein synthesis through inactivation of the eukaryotic translation initiation factor 2 in response to various stress conditions. Mutations in *EIF2AK3* are associated with Wolcott-Rallison syndrome, which is characterised by neonatal diabetes, growth retardation, and skeletal dysplasia.

Other related BSM genes include *ANKRD27*, which is associated with parastremmatic dwarfism, a rare bone disease characterised by severe dwarfism and distortion of lower limbs; *LCORL*, with polymorphisms in this gene associated with skeletal frame size and adult height; and *NPR3*, encoding a receptor for natriuretic peptide hormones, binding the atrial, brain, and C-type natriuretic peptides. *NPR3* plays a role in clearing circulating and extracellular natriuretic peptides through endocytosis, regulating their local concentrations and effects, thus regulating blood pressure, growth, skeletal development, and other processes. It has been shown to play a role in the regulation of linear bone growth with loss of function mutations causing enhanced growth in humans characterised by tall stature, long digits, extra epiphyses in the hands and feet, and cardiovascular abnormalities (*169*).

#### <u>8.2 Signalling cascades</u>

The most significant selective signatures in the BSM gene set relate to signalling pathways and cellular signalling. There is strong overlap across several clusters in the BSM gene set, which we describe briefly here. The top GO cluster includes three BP GO terms (enrichment score 3.07) for the ERK1 and ERK2 cascade and its regulation with nine genes. Four of these genes are also found in the growth factor related GO terms discussed above (*FGF1*, *FGF20*, *TGFB1*, *DUSP6*), plus five other genes of which four encode chemokines (*CCL1, CCL17, CCL20, CCL8, ADCYAP1*). The second GO cluster includes five BP GO terms (enrichment score 2.4) for the MAPK cascade and its regulation with a total of 19 genes of which 13 are common to all GO terms. All nine genes in the ERK1/2 cluster are found in this cluster along with genes discussed above for the growth factor related GO terms (e.g., *BMP15*, *GDF10*, *NGFR*, *SOX11*) and some other genes (*CD40, DACT1*, *HRH4*, *RIPK3, TNFRSF1B, IL6R*). Most of the top individual GO terms are found in these signalling pathway (and cytokine, see below) clusters. The top GO annotation according to EASE score is the BP GO term "regulation of MAPK cascade" with 17 genes (3 fold enrichment), followed by MF GO terms "cytokine receptor binding" and "cytokine activity" (4.5 and 5 fold enrichment, 10 and 9 genes), then various BP GO terms related to the MAPK/ERK cascades, protein phosphorylation, chemotaxis, and interleukin-6 production.

In addition to overlap with growth factor related GO terms, the genes in the ERK1/2 and MAPK signalling pathway clusters overlap with other clusters relating to cytokines and specific clusters for interleukin and chemokine subclasses of cytokines (discussed in the next section). In combination, both the cytokine and growth factor signals appear to underly the particularly strong signatures for the ERK/MAPK signalling cascades. Notably, only three (of 19) genes in the MAPK cascade cluster are not found in either the growth factor related GO terms, or the chemokine and interleukin-6 related cytokine GO clusters (*CD40*, *DACT1*, *HRH4*). Thus, selection on ERK/MAPK cascades may be related to the reduced body size of squirrel monkeys owing to their role in cell differentiation/proliferation/growth in association with growth factors (which are discussed in section 8.1), and related to immunity owing to their role in inflammation and stress responses in association with cytokines (especially chemokines and interleukin-6; discussed in the following section) (*170*).

Similar signatures related to signalling cascades can be found in other annotation categories in the BSM gene set including the enriched UP keywords "signal" (64 genes, 1.7 fold enrichment) and "secreted" (36 genes, 2 fold enrichment). In addition, there are some genes involved in the MAPK and ERK1/2 cascades in the BM gene set including *MAPK7*, which plays a role in various cellular processes such as proliferation, differentiation, and cell survival, and phosphorylates the product of *SGK1* which is required for growth factor-induced cell cycle progression.

#### 8.3 Inflammation and immunity related

As discussed in the previous section, there are signatures of selection for the BSM gene set on cytokines. There are various enriched terms relating to cytokines including the BP GO terms “cytokine-mediated signalling”, “positive regulation of cytokine production”, and “cellular response to cytokine stimulus”, MF GO terms “cytokine receptor binding” and “cytokine activity”, and UP keyword “cytokine”. Many of the genes in these broader cytokine terms are found in specific clusters related to interleukin-6 and chemokine subclasses of cytokines, which form largely non-overlapping clusters of genes, as well as growth factor related genes.

Regarding the immunity related cytokine signals, the third cluster (enrichment score 2.17) contains three BP GO terms related to interleukin-6 production with five genes (5.5 to 9.2 fold enrichment); *IL6R*, *IL1RAP*, *IL36A*, *ADCYAP1*, and *PAEP*. *IL6R* encodes a subunit of the interleukin 6 receptor complex; interleukin 6 is an important multifunctional cytokine that plays an essential role in the immune response. *IL36A* encodes a cytokine that forms part of the IL-36 signalling system present in epithelial barriers, and can activate NF-kappa-B and MAPK signalling pathways to generate an inflammatory response.

The fifth cluster (enrichment score 1.88) contains 11 GO terms (BP and MF) that relate to chemokines, chemotaxis, and leukocyte migration (4 to 5 genes, 3.8 to 12.3 fold enrichment). The four genes common to all GO terms are chemokines *CCL1*, *CCL17*, *CCL20*, and *CCL8*. Chemokines’ main function is in chemotaxis; acting as a chemoattractant to manage the migration of leukocytes in both inflammatory and homeostatic processes, thus guiding cells of both innate and adaptive immune systems. *CCL1* shows chemotactic activity for monocytes; *CCL17* plays important roles in T cell development in thymus as well as in trafficking and activation of mature T cells; *CCL8* (also in the BM gene set) is chemotactic for many cell types including monocytes, lymphocytes, basophils, and eosinophils, and is a potent inhibitor of HIV1; and *CCL20* shows chemotactic activity for dendritic cells, effector/memory T-cells, and B-cells, and plays an important role at skin and mucosal surfaces under homeostatic and inflammatory conditions, as well as various other functions. Among the most enriched related GO terms not in this cluster is the BP term “chemotaxis” with 13 genes (2.8 fold enrichment) including these chemokine genes, some growth factor related genes, as well as others.

Another of the top most enriched BP GO terms for the BSM gene set is “inflammatory response” with 15 genes (2.8 fold enrichment) including most of the genes in the interleukin-6 and chemokine related clusters. Some other genes include *CD40* (also found in the BM gene set), which encodes a receptor found on antigen-presenting cells that is essential for mediating a wide range of immune and inflammatory responses including T cell-dependent immunoglobulin class switching and memory B cell development; *CD5L*, which encodes a key regulator of lipid synthesis that is primarily expressed by macrophages in lymphoid and inflamed tissues and regulates mechanisms in inflammatory responses, for example, participating in obesity-associated inflammation; and *HRH4* (also in the BM gene set), which encodes the H4 subclass of histamine receptors thought to play a role in inflammation and allergy responses.

Similar signatures related to inflammation and immunity can be found for the BSM gene set in the enriched UP keywords “cytokine”, “chemotaxis”, and “inflammatory response” (5 to 8 genes, 4.4 to 5.8 fold enrichment). These signatures are also reflected in the KEGG pathway “cytokine-cytokine receptor interaction” with 11 genes (7.4 fold enrichment) and the enriched Reactome pathway “signalling by interleukins” (4 genes, 11.5 fold enrichment). The BM gene set is enriched for the disease annotation “acquired immunodeficiency syndrome | disease progression” with 34 genes (1.8 fold enrichment). Most of the other enriched disease annotations for the BM gene set are derived from hits to the same few genes and not considered in detail.

#### 8.4 Brain development and neuronal related

The BSM gene set is also enriched for brain and neuronal related signatures. The 6$^{th}$ and 8$^{th}$ clusters consist of three and six BP GO terms (enrichment scores of 1.49 and 1.67) describing the regulation of neuron differentiation, nervous system development, neuron projection development, and neurogenesis (6 to 13 genes, 2 to 3.2 fold enrichment). Several of these genes are found in the growth factor related GO terms discussed in section one, including *NTF4*, *SOX11*, *NGFR*, *TGFB1*, and *FGF20*. Other aforementioned genes in the cytokine/interleukin GO terms include *ADCYAP1*, which promotes neuron projection development and plays a role in the neuroendocrine stress response (among other functions), and *IL1RAP*, which can bidirectionally induce pre- and post-synaptic differentiation of neurons. Distinct genes found in the neuron GO terms include *SEMA4F*, which plays a role in neural development; and two golgin family members, *GOLGA4* and *GORAB*, involved in vesicular trafficking.

Other related BP GO terms not in these clusters for the BSM gene set are “neuron projection development” (14 genes, 2.1 fold enrichment) and “neural tube development” (5 genes, 3.8 fold enrichment) with several of the same genes from the neural development cluster, and “photoreceptor cell maintenance” (3 genes, 10.7 fold enrichment). Several of the genes mentioned in this section are also found in the enriched disease annotation “Alzheimer’s disease” (16 genes, 1.9 fold enrichment). Among other enriched terms for the BM gene set is the UP keyword “non-syndromic deafness” (6 genes, 3.3 fold enrichment).

Other interesting related genes found in the BM gene set include *SIX3*, which plays many important roles in brain development including in forebrain patterning, and is required for ependymal cell maturation at postnatal stages of brain development and for neuroretina development with key roles in early lens formation, among others; *LHX4*, which encodes a transcription factor involved in the control of differentiation and development of the pituitary gland, and implicated in pituitary hormone deficiency; and *HMX2*, a NKL homeobox transcription factor involved in specification of neuronal cell types, required for inner ear and hypothalamus development, and implicated in inner ear malformations and hearing loss.

8.5 Neuromodulators

Among the most enriched BP GO terms for the BM gene set are “dopamine metabolic process” and “dopamine biosynthetic process” (7.1 and 13.2 fold enrichment, respectively). The four genes between these GO terms are *DAO*, which regulates the level of the neuromodulator D-serine in the brain, and contributes to dopamine synthesis; *GPR37*, a receptor for the neuro- and glio-protective factor, prosaposin, that interacts with Parkin and is implicated in juvenile Parkinson’s disease; *GCH1*, which may be involved in dopamine synthesis and pain sensitivity; and *SNCB*, which encodes a protein that may function in neuronal plasticity, and is a non-amyloid component of senile plaques found in AD. A related interesting gene in the BSM gene set is *LMX1A*, which plays a role in the development of dopamine producing neurons during embryogenesis; mutations in *LMX1A* are associated with an increased risk of Parkinson's disease.

Other interesting related genes found in the BM gene set include *NPSR1*, a GPCR for neuropeptide S associated with asthma susceptibility, panic disorders, inflammatory bowel disease, and rheumatoid arthritis; *NMU*, a neuromedin neuropeptide that plays a role in pain, stress, immune-mediated inflammatory diseases, and feeding regulation; and *NRG3*, a susceptibility locus for schizophrenia and schizoaffective disorder in linkage studies.

#### 8.6 Other anatomical and developmental signatures

For the BSM gene set, there is a cluster of four GO terms (enrichment score 1.61) including the BP GO terms "anatomical structure development" and "developmental process" (52 to 59 genes, ~1.3 fold enrichment). Some of the genes in this cluster are found in the growth factor and neural development related GO terms and clusters. A few other interesting genes include *HOXB4*, a homeobox gene involved in development; *COL4A4*, which encodes one of the six subunits of type IV collagen; *NOTO*, a transcription regulator with important roles in notochord development; *TMEM88*, which plays a crucial role in heart development; and *OPN1SW* (see below). Important developmental genes in the BM gene set includes another homeobox gene, *HOXB2*, and *SCX*, which plays an essential early role in the formation of the mesoderm and somite-derived chondrogenic lineages.

For the BM gene set, there are several enriched BP GO terms related to fibroblast proliferation and regulation with seven genes (3.8 to 3.9 fold enrichment). Fibroblasts are the main connective tissue cells, responsible for making the extracellular matrix and collagen, which together form the structural framework of tissues in animals and play an important role in tissue repair. There are also several genes related to plasmin in the gene sets; plasmin plays a key role in blood coagulation and fibrinolysis. These include *PLAUR* (BSM gene set), which plays a role in localising and promoting plasmin formation; and *PLAT* (BM gene set), which encodes a secreted serine protease that converts the proenzyme plasminogen to plasmin, thus playing an important role in tissue remodelling and degradation.

#### 8.7 Peroxisomes, mitoribosomes, and oxidative stress

Among the most enriched GO terms for the BM gene set is the CC term "peroxisome" (and "microbody") with 10 genes (3.5 fold enrichment). These same genes comprise the enriched KEGG pathway "peroxisome" (8 genes, 5.9 fold enrichment) and CC term "peroxisomal part" (6 genes, 3.1 fold enrichment). Peroxisomes are primarily involved in lipid metabolism, the conversion of reactive oxygen species (ROS), and the biosynthesis of plasmalogens, which are critical for brain and lung function in mammals. The selective signature on peroxisomes is interesting considering the reduced body size of squirrel monkeys; the metabolic rate and the rate of ROS formation and, thus, the potential for oxidative stress, is inherently greater in smaller-sized mammals (*171*)Indeed, several of the genes in the peroxisome annotations are specifically linked to the of ROS, such as *EPHX2*, which encodes a member of the epoxide hydrolase family that degrades toxic epoxides (alkene oxides, oxiranes) and arene oxides; *SOD1*, which encodes one of two isozymes responsible for converting naturally-occurring but toxic free superoxide radicals; and *MPV17*, see below. In line with this, among the most enriched MF GO terms for the BM gene set is "superoxide-generating NADPH oxidase activator activity" i.e., genes relating to increased activity of the enzyme superoxide-generating NADPH oxidase (3 genes, 14.2 fold enrichment). One of these genes is *NOXO1*, which encodes an NADPH oxidase (NOX) organiser, which positively regulates the superoxide-generating activity of NADPH oxidases, *NOX1* and *NOX3*. A related gene in the BSM gene set is *NOXA1*, which encodes a protein which activates *NOX1* (and putatively *NOX3*) and functions in the production of ROS.

Another related enriched GO annotation in the BM gene set is the BP term "cellular response to oxidative stress" with 11 genes (2.3 fold enrichment). These genes include *MPV17*, which encodes a non-selective channel that modulates inner mitochondrial membrane potential under normal conditions and oxidative stress, and is involved in mitochondrial homeostasis, mitochondrial DNA maintenance, the regulation of ROS metabolism, and the control of oxidative phosphorylation; *MGMT*, which encodes a DNA repair protein that is involved in cellular defence against mutagenesis and toxicity from alkylating agents (potent carcinogens), and repairs toxic lesions; *GLRX2*, which encodes an oxidoreductase involved in mitochondrial redox homeostasis, the protection against and recovery from oxidative stress, and the regulation of superoxide production by mitochondrial complex I; *HMOX1*, which encodes heme oxygenase, an essential enzyme in heme catabolism that cleaves heme to form biliverdin, and shows cytoprotective effects as excess of free heme sensitises cells to undergo apoptosis; and *OXR1*, thought to be involved in protection from oxidative damage.

Furthermore, potentially in relation to an increased metabolic rate, there are signatures of selection on mitochondria and mitoribosomes. Another of the most enriched GO annotations for the BM gene set is the CC term "large ribosomal subunit" (9 genes, 3.6 fold enrichment), which is also found in the top GO cluster along with two other CC terms, "ribosome" and "ribosomal subunit". Of the nine genes in this GO, six are nuclear-encoded mitochondrial ribosomal proteins that form the large (39S) subunit of the mitoribosome; *MRPL1*, *MRPL14*, *MRPL17*, *MRPL19*, *MRPL24*, and *MRPL55*. Other genes in this GO term are *RPL11*, which similarly encodes a protein that is a component of the large (60S) subunit of the ribosome; and *MRTO4*, which encodes a component of the ribosome assembly machinery. Other top GO terms related to the ribosome/mitoribosome include the MF term "structural constituent of ribosome" (12 genes, 2.6 fold enrichment), which also includes *ANKRD42* and three solute carrier family 25 genes (*SLC25A18*, *SLC25A21*, *SLC25A35*), which transport a variety of compounds across the inner mitochondrial membranes. Another gene in the BM gene set related to mitoribosomes is *NGRN*, which plays an essential role in mitoribosome biogenesis and is required for intra-mitochondrial translation of core subunits of the oxidative phosphorylation system. Other enriched GO terms comprised solely of the mitochondrial ribosomal proteins include the CC terms "mitochondrial large ribosomal subunit" and "organellar large ribosomal subunit", and the BP terms "mitochondrial translational elongation" and "mitochondrial translational termination" (5 to 6 genes, 3.4 to 4.9 fold enrichment), which together form the third GO cluster (enrichment score 1.56) for the BM gene set. Similarly, the second, third and fourth most enriched Reactome pathways describe mitochondrial translation initiation, termination, and elongation, with the same six genes encoding mitochondrial ribosomal proteins (all 3.7 fold enrichment). Other genes in the BM gene set related to the mitochondrion include *NDUFAF5*, which is essential for the assembly of mitochondrial complex I located in the inner mitochondrial membrane, with mutations in this gene causing mitochondrial complex I deficiency; *COX16*, which is essential for the assembly of the mitochondrial respiratory chain complex IV; and *COX11*, which plays a role in terminal stages of COX synthesis.

### 8.8 Nutrition and metabolism

The ninth GO cluster for the BM gene set (enrichment score 1.32) contains three terms related to cellular calcium and metal ion homeostasis (8 to 9 genes, 2.2 to 2.5 fold enrichment). These GO terms include several aforementioned genes (*CCL1*, *CCL8*, *CD40*, *TGFB1*, *ADCYAP1*, *HRH4*), as well as *HRC*, which plays a role in the regulation of releasable calcium into the sarcoplasmic reticulum of skeletal muscle, and *PTH1R*, which encodes a receptor for parathyroid hormone with a central role in regulating calcium ion homeostasis. Another gene in the BM gene set is *STOML2*, which encodes a mitochondrial protein that may regulate biogenesis and activity of mitochondria, and play a role in calcium homeostasis through negative regulation of calcium efflux from mitochondria. A related enriched KEGG pathway for the BM gene set is “mineral absorption” (4 genes, 5.5 fold enrichment) including the gene *FTH1*, which encodes the heavy subunit of ferritin, the major intracellular iron storage protein that stores iron in a soluble, non-toxic form, is important for iron homeostasis, and plays a role in delivery of iron to cells.

There are several GO terms for the BM gene set related to lactate transport including the BP terms “plasma membrane lactate transport”, “lactate transmembrane transport”, and “lactate transport”, as well as the MF term “lactate transmembrane transporter activity” (11.1 to 12.9 fold enrichment) all with the same three genes (*SLC16A3*, *SLC16A4*, *SLC16A11*) and forming the second GO cluster (enrichment score 1.59). These genes are all SLC16 family members, which catalyse rapid transport across the plasma membrane of many monocarboxylates such as lactate, pyruvate, branched-chain oxo acids derived from leucine, valine, and isoleucine, and some ketone bodies. They are involved in a range of metabolic pathways including energy metabolism of the brain, skeletal muscle, and heart, gluconeogenesis, bowel metabolism, TH metabolism, and drug transport (*172*).

Among the enriched GO terms for the BM gene set is the BP term “retinoid metabolic process” (6 genes, 3.3 fold enrichment). Retinoids (vitamin A) are necessary for growth, reproduction, differentiation of epithelial tissues, vision, and immune competence, with important roles in the developing nervous system, notochord, and other embryonic structures. Genes in this GO term include *OPN1SW*, opsin 1 short wave sensitive, which encodes the blue cone visual pigment (also in the BSM gene set); *RHO*, rhodopsin, which encodes the photoreceptor found in rod cells in the back of the eye that is required for vision in low light conditions and for photoreceptor cell viability after birth; and *RBP2*, which encodes a protein abundant in the small intestinal epithelium involved in the intracellular transport of retinol (vitamin A). There are also several lipid related enriched BP GO terms in the BM gene set including “cellular lipid metabolic process” (32 genes, 1.5 fold enrichment), “lipoprotein biosynthetic process” (6 genes, 3.1 fold enrichment), and “positive regulation of lipid metabolic process” (7 genes, 2.7 fold enrichment).

8.9 Other

For the BM gene set, the top nine individual GO terms are CC terms, and generally, there are a large number of enriched CC GO terms, for example, representing 34 of the 70 GO terms enriched for this gene set. Many of these CC terms have only a small-to-modest fold enrichment but the significance is driven by the large number of genes including “cytoplasm”, “cytoplasmic part”, “endomembrane system”, “organelle membrane”, “organelle part”, “intracellular part”, “intracellular organelle lumen”, and “intracellular membrane bound organelle” (76 to 316 genes, 1.1 to 1.4 fold enrichment). There are several enriched mitochondrion related CC terms such as “mitochondrion”, “mitochondrial part”, “mitochondrial matrix”, and “mitochondrial inner membrane” (16 to 49 genes, 1.4 to 1.8 fold enrichment). Other enriched CC terms include “Golgi apparatus” (47 genes, 1.5 fold enrichment), “envelope” (38 genes, 1.6 fold enrichment), “centriole” (8 genes, 3.4 fold enrichment), “actin cytoskeleton” (18 genes, 1.8 fold enrichment), “endoplasmic reticulum lumen” (10 genes, 2.3 fold enrichment), “nuclear membrane” (12 genes, 1.9 fold enrichment), “endoplasmic reticulum-Golgi intermediate compartment membrane” (5 genes, 3.7 fold enrichment), and “microtubule organising centre part” (8 genes, 1.8 fold enrichment), among others. Similarly, “mitochondrion” and “Golgi apparatus” are enriched UP keywords for the BM gene set (35 and 27 genes, 1.5 to 1.6 fold enrichment).

For the BM gene set, there are two BP GO terms related to glycosaminoglycan (GAG) biosynthesis with eight genes (3.4 to 3.5 fold enrichment). GAGs are polysaccharides consisting of repeating disaccharide units with four primary groups: heparan sulphate (HS), chondroitin sulphate, keratan sulphate, and hyaluronic acid. They are involved in cell hydration and structural scaffolding, and play a key role in cell signalling thus modulating a range of biological processes. Four genes in the GAG GO terms are specific to HS and appear to primarily drive the enriched GAG signal; *GLCE*, which modifies maturing HS and heparin allowing further modifications that determine the specificity of protein interactions; *SDC2*, which encodes a transmembrane HS proteoglycan that participates in cell proliferation, migration, and cell-matrix interactions; and two genes encoding HS biosynthetic enzymes, *HS3ST1* and *HS3ST5*, both are crucial rate limiting enzymes for the synthesis of anticoagulant HS. The signature of selection on HS biosynthesis is reflected by the enriched Reactome pathway “HS-GAG biosynthesis” with these same four genes (6.8 fold enrichment). Two other genes in the GAG GO terms are related to glycosphingolipid biosynthesis, *B3GNT3* and *B4GALT1*, the latter of which catalyses the production of lactose in the lactating mammary gland (the Golgi complex form). These two genes, along with two others (*FUT1* and *FUT4*) comprise the enriched KEGG pathway "glycosphingolipid biosynthesis - lacto and neolacto series" (9.4 fold enrichment).

Other enriched disease annotations for the BSM gene set include several related to viral respiratory infections (9 genes, 4.9 to 5.1 fold enrichment) and chorioamnionitis (8 genes, 4.9 fold enrichment), as well as the terms "rheumatoid arthritis" (7 genes, 4.6 fold enrichment), "multiple sclerosis" (13 genes, 2.6 fold enrichment), "premature birth" (5 genes, 5.9 fold enrichment), and "bone density" (7 genes, 3.8 fold enrichment).

**Table S1** NCBI SRA accessions and sequencing information for all WGS, Chicago, and RNAseq libraries for the *Sapajus apella* reference individual, Mango.

| Library | Accession | Read info. | Platform | # read pairs | # lanes |
|---|---|---|---|---|---|
| Whole genome shotgun (WGS) | SRR14087928, SRR14113847 | 150bp PE | HiSeq 4000 | 1,330,334,107 | 4 |
| Chicago library 1 | SRR14087927 | 100bp PE | HiSeq 4000 | 312,672,706 | 2 |
| Chicago library 2 | SRR14087937 | 100bp PE | HiSeq 4000 | 247,824,943 | |
| Chicago library 3 | SRR14087935 | 100bp PE | HiSeq 4000 | 240,427,607 | |
| | | | All Chicago: | 800,925,256 | |
| RNAseq: Cerebellum | SRR14087926 | 150bp PE | HiSeq 3000 | 19,288,395 | 1 |
| RNAseq: Kidney | SRR14087929 | 150bp PE | HiSeq 3000 | 17,731,806 | |
| RNAseq: Lung | SRR14087930 | 150bp PE | HiSeq 3000 | 19,626,504 | |
| RNAseq: Temporal lobe | SRR14087931 | 150bp PE | HiSeq 3000 | 24,670,464 | |
| RNAseq: Colon | SRR14087932 | 150bp PE | HiSeq 3000 | 16,796,196 | |
| RNAseq: Thymus | SRR14087933 | 150bp PE | HiSeq 3000 | 21,435,567 | |
| RNAseq: Pituitary | SRR14087934 | 150bp PE | HiSeq 3000 | 24,753,830 | |
| RNAseq: Liver | SRR14087936 | 150bp PE | HiSeq 3000 | 17,451,342 | |
| RNAseq: Cerebrum | SRR14087938 | 150bp PE | HiSeq 3000 | 23,357,124 | |
| RNAseq: Mesenteric LN | SRR14087939 | 150bp PE | HiSeq 3000 | 23,301,230 | |
| RNAseq: Muscle | SRR14087940 | 150bp PE | HiSeq 3000 | 27,385,357 | |
| RNAseq: Aorta | SRR14087941 | 150bp PE | HiSeq 3000 | 26,855,559 | |
| RNAseq: Ovary | SRR14087942 | 150bp PE | HiSeq 3000 | 18,782,482 | |
| RNAseq: Duodenum | SRR14087943 | 150bp PE | HiSeq 3000 | 23,438,737 | |
| RNAseq: Hippocampus | SRR14087944 | 150bp PE | HiSeq 3000 | 22,371,719 | |
| RNAseq: Midbrain | SRR14087945 | 150bp PE | HiSeq 3000 | 19,849,072 | |
| RNAseq: Bone marrow | SRR14087946 | 150bp PE | HiSeq 3000 | 19,700,417 | |
| | | | All RNAseq: | 366,795,801 | |

**Table S2** Genome size estimates from four methods.

| **Method** | **Genome size (bp)** |
|---|---|
| GenomeScope | 2,917,676,754 |
| Jellyfish stats | 3,003,998,946 |
| Formula[1] | 3,014,334,525 |
| findGSE | 3,029,414,613 |

[1]Liu et al. (2014)

**Table S3** RNAseq read pair and total base counts after quality filtering steps.

| Processing step | # Read pairs | # Bases |
|---|---|---|
| Raw | 366,795,801 | 102,498,243,858 |
| rCorrector & Trim 1 | 351,721,049 | 98,256,604,277 |
| rRNA removal | 341,717,917 | 95,454,377,196 |
| Trim 2 | 340,957,550 | 95,245,746,882 |
| Normalisation | 27,205,724 | 7,546,788,011 |

Table S4 is an external excel file

**Table S5** Repeat content of the robust capuchin (*Sapajus apella*) genome estimated with libraries of known repeats (RepBase) and *de novo* repeat identification (RepeatModeler).

| | **Final Combined** | | | **RepBase[2]** | | | **RepeatModeler[2]** | | |
|---|---|---|---|---|---|---|---|---|---|
| **Repeat Type[1]** | **# Elements** | **Length (bp)** | **% Genome** | **# Elements** | **Length (bp)** | **% Genome** | **# Elements** | **Length (bp)** | **% Genome** |
| SINEs: | 1,357,677 | 279,256,097 | 11.08% | 1,357,123 | 279,102,605 | 11.07% | 169 | 15,427 | 0.00% |
| ALUs | 908,257 | 212,167,670 | 8.42% | 907,882 | 212,014,178 | 8.41% | 169 | 15,427 | 0.00% |
| MIRs | 442,827 | 66,274,190 | 2.63% | 442,648 | 66,274,190 | 2.63% | 0 | 0 | 0.00% |
| LINEs: | 911,478 | 498,029,165 | 19.76% | 884,120 | 494,201,421 | 19.61% | 26,347 | 3,834,920 | 0.15% |
| LINE1 | 558,958 | 404,928,706 | 16.07% | 536,992 | 402,065,602 | 15.95% | 21,489 | 2,870,692 | 0.11% |
| LINE2 | 300,196 | 80,857,408 | 3.21% | 295,468 | 80,370,514 | 3.19% | 4,225 | 486,369 | 0.02% |
| L3/CR1 | 38,137 | 8,454,735 | 0.34% | 38,113 | 8,454,735 | 0.34% | 0 | 0 | 0.00% |
| LTRs: | 472,219 | 206,887,322 | 8.21% | 435,273 | 197,412,384 | 7.83% | 37,134 | 9,518,001 | 0.38% |
| ERVL | 99,972 | 47,811,480 | 1.90% | 99,757 | 47,806,711 | 1.90% | 87 | 4,769 | 0.00% |
| ERVL-MaLRs | 223,928 | 91,264,595 | 3.62% | 223,383 | 91,194,751 | 3.62% | 423 | 70,327 | 0.00% |
| ERV_classI | 113,563 | 58,031,173 | 2.30% | 87,702 | 51,460,224 | 2.04% | 26,163 | 6,600,128 | 0.26% |
| ERV_classII | 11,821 | 3,971,540 | 0.16% | 1,617 | 1,172,629 | 0.05% | 10,369 | 2,812,269 | 0.11% |
| DNA Transposons: | 424,122 | 95,927,338 | 3.81% | 385,929 | 90,582,536 | 3.59% | 38,199 | 5,351,972 | 0.21% |
| hAT-Charlie | 196,123 | 38,993,315 | 1.55% | 196,123 | 38,993,315 | 1.55% | 0 | 0 | 0.00% |
| TcMar-Tigger | 97,595 | 33,101,189 | 1.31% | 92,017 | 32,059,351 | 1.27% | 5,580 | 1,041,985 | 0.04% |
| Unclassified | 20,311 | 4,020,945 | 0.16% | 5,162 | 971,339 | 0.04% | 15,080 | 3,085,774 | 0.12% |
| **Total Interspersed Repeat Content** | 3,185,807 | **1,084,120,867** | **43.02%** | 3,067,607 | 1,062,270,285 | 42.15% | 116,929 | 21,806,094 | 0.87% |
| Small RNA | | 1,678,548 | 0.07% | | NA | NA | | NA | NA |
| Satellites | | 7,543,749 | 0.30% | | NA | NA | | NA | NA |
| Simple Repeats | | 26,560,076 | 1.05% | | NA | NA | | NA | NA |
| Low Complexity | | 5,132,506 | 0.20% | | NA | NA | | NA | NA |
| **Total Repeat Content** | | **1,124,865,661** | **44.63%** | | NA | NA | | NA | NA |

[1]SINE: short interspersed nuclear element; LINE: long interspersed nuclear element; LTR: long terminal repeat retrotransposons.

[2]Only the interspersed repeats are presented as we didn't analyse low complexity and simple repeats in both runs.

**Table S6** Comparison of evidence used in each iteration of Maker.

| | **Pass 1** | **Pass 2** | **Pass 3** |
|---|---|---|---|
| Transcript evidence | PASAv1, TrinDNv1 | PASAv1, TrinDNv1 | NRv1 |
| Homology: Proteomes | *Cebus, Saimiri, Callithrix, Aotus, Homo* | *Cebus, Saimiri, Callithrix, Aotus, Homo* | *Cebus, Saimiri, Callithrix, Homo* |
| Homology: Mammalian SwissProt/UniProtKB | Y | Y | N |
| Aligned evidence (est/prot2genome) | Y | N | N |
| *ab initio* HMM | BUSCO | Augustus v1 | Augustus v2 |
| Training models | AED<0.25; >50 aa | AED<0.1; >70 aa | NA |
| Augustus info. | v1; with PASAv1 | v2; with NRv1 | NA |

**Table S7** Input sequences (OrthoMCL), group (ortholog, alignment), and species set counts per species.

| Species ID | Genus | Species | # input seqs. | # orthologs | # initial 1-to-1 orthologs (of 12,160) | # final alignments (of 9,216) | Missing code | # final species sets (of 207) |
|---|---|---|---|---|---|---|---|---|
| Sape | *Sapajus* | *apella* | 25,279 | 16,164 | 8,669 | 7,134 | _1 | 140 |
| Cimi | *Cebus* | *imitator* | 20,309 | 18,824 | 10,229 | 9,092 | _2 | 177 |
| Sbol | *Saimiri* | *boliviensis* | 19,372 | 18,030 | 9,922 | 9,003 | _3 | 167 |
| Cjac | *Callithrix* | *jacchus* | 19,669 | 18,525 | 10,111 | 8,921 | _4 | 135 |
| Mmul | *Macaca* | *mulatta* | 21,047 | 18,742 | 9,965 | 8,179 | _5 | 135 |
| Ptro | *Pan* | *troglodytes* | 23,513 | 21,148 | 10,848 | 9,041 | _6 | 150 |
| Hsap | *Homo* | *sapiens* | 20,982 | 20,371 | 9,601 | 8,426 | _7 | 139 |
| Csyr | *Carlito* | *syrichta* | 18,387 | 16,644 | 8,948 | 7,734 | _8 | 101 |
| Mmur | *Microcebus* | *murinus* | 18,885 | 17,676 | 9,510 | 8,645 | _9 | 118 |
| Mmus | *Mus* | *musculus* | 22,794 | 19,753 | 9,861 | 8,645 | _10 | 122 |

Tables S8 and S9 are external excel files

**Table S10** Counts of groups, models, and significant results for BM and BSM tests with PAML.

| | **Robust capuchin (H1)** | **Gracile capuchin (H2)** | **Ancestral Cebinae (H3)** | **Across-Cebinae (H3a)** | **Ancestral Cebidae (H4)** | **Squirrel monkey (H5)** | **Of/total** | **Average** |
|---|---|---|---|---|---|---|---|---|
| **Total groups** | **7,010** | **7,010** | **6,978** | **9,003** | **8,740** | **9,003** | **9,216** | **7,957** |
| **Groups with all (10) species** | 4,636 | 4,636 | 4,636 | 4,636 | 4,636 | 4,636 | 4,636 | 4,636 |
| **Groups with 9 species** | 1,695 | 1,695 | 1,695 | 2,756 | 2,701 | 2,756 | 2,819 | 2,216 |
| **Groups with 5 to 8 species** | 679 | 679 | 647 | 1,611 | 1,403 | 1,611 | 1,761 | 1,105 |
| **Total model tests: BM & BSM** | 14,020 | 14,020 | 13,956 | 9,003 | 17,480 | 18,006 | 86,485 | |
| **Total analyses: Start values per null (1) + alt. (3) model** | 56,080 | 56,080 | 55,824 | 36,012 | 69,920 | 72,024 | 345,940 | |
| **Accelerated models: BM** | **292** | **248** | **302** | **552** | **278** | **435** | **2,107** | **351** |
| **Significant models: BSM** | **80** | **75** | **122** | **NA** | **104** | **186** | **567** | **113** |
| **Model overlap: BM & BSM** | 17 | 18 | 30 | NA | 26 | 34 | 125 | 25 |
| **% Accelerated models: BM** | 4.17% | 3.54% | 4.33% | 6.13% | 3.18% | 4.83% | | 4.36% |
| **% Significant models: BSM** | 1.14% | 1.07% | 1.75% | NA | 1.19% | 2.07% | | 1.44% |
| **FDR Accelerated models: BM** | 3 | 0 | 4 | 98 | 5 | 19 | 129 | 22 |
| **FDR Significant models: BSM** | 0 | 3 | 2 | NA | 5 | 5 | 15 | 3 |

Tables S11 to S22 are external excel files

**Table S23** Counts of enriched annotation categories and GO clusters per gene set enrichment analysis with key to detailed results tables.

| | **H1** | | **H2** | | **H3** | | **H3a** | **H4** | | **H5** | | **Avg.** | |
|---|---|---|---|---|---|---|---|---|---|---|---|---|---|
| | **BM** | **BSM** | **BM** | **BSM** | **BM** | **BSM** | **BM** | **BM** | **BSM** | **BM** | **BSM** | **BM** | **BSM** |
| Num. of genes | 292 | 80 | 248 | 75 | 302 | 122 | 552 | 278 | 104 | 435 | 186 | 351 | 113 |
| GO clusters | 7 | 2 | 2 | 0 | 4 | 1 | 5 | 13 | 3 | 4 | 9 | 6 | 3 |
| **Enriched Annotations** | | | | | | | | | | | | | |
| BP GO terms[1] | 57 | 27 | 45 | 0 | 36 | 17 | 68 | 100 | 32 | 31 | 80 | 56 | 31 |
| MF GO terms[1] | 6 | 0 | 5 | 1 | 6 | 0 | 10 | 14 | 3 | 5 | 13 | 8 | 3 |
| CC GO terms[1] | 9 | 1 | 8 | 3 | 17 | 3 | 9 | 33 | 13 | 34 | 2 | 18 | 4 |
| Total GO terms | 72 | 28 | 58 | 4 | 59 | 20 | 87 | 147 | 48 | 70 | 95 | 82 | 39 |
| UP keyword | 9 | 1 | 9 | 6 | 6 | 1 | 11 | 20 | 10 | 5 | 7 | 10 | 5 |
| Disease | 4 | 44 | 1 | 0 | 3 | 2 | 3 | 16 | 1 | 7 | 18 | 6 | 13 |
| KEGG pathway | 1 | 0 | 0 | 0 | 3 | 0 | 2 | 3 | 1 | 3 | 1 | 2 | 0 |
| Reactome pathway | 0 | 0 | 0 | 0 | 3 | 5 | 8 | 3 | 4 | 6 | 2 | 3 | 2 |
| Total (all categories) | 86 | 73 | 68 | 10 | 74 | 28 | 111 | 189 | 64 | 91 | 123 | 103 | 60 |
| **Results Tables ID** | | | | | | | | | | | | | |
| All enriched terms | S24 | S25 | S28 | S29 | S31 | S32 | S35 | S37 | S38 | S41 | S42 | NA | NA |
| GO clusters | S26 | S27 | S30 | None | S33 | S34 | S36 | S39 | S40 | S43 | S44 | NA | NA |

[1] BP: biological process; MF: molecular function; CC: cellular component

Tables S24 to S44 are external excel files

Table legends for the 36 supplementary tables found as excel files:

**Table S4** Comparison of counts, total bases, genome alignment, and BUSCO results for the seven RNAseq transcript assemblies.
**Table S8** Group information including assigned gene symbol, description, and Entrez ID, species set, lineages tested in PAML, and Ensembl (or Sape) ID for the sequence in the alignment.
**Table S9** Counts of groups (alignments) and species sets per lineage and species, along with other species set information including lineages analysed, species included, and tree setup.
**Table S11** Full list of groups analysed for H1 (robust capuchins) including gene information and BM/BSM result.
**Table S12** Full list of groups analysed for H2 (gracile capuchins) including gene information and BM/BSM result.
**Table S13** Full list of groups analysed for H3 (ancestral Cebinae) including gene information and BM/BSM result.
**Table S14** Full list of groups analysed for H3a (across-Cebinae) including gene information and BM result.
**Table S15** Full list of groups analysed for H4 (ancestral Cebidae) including gene information and BM/BSM result.
**Table S16** Full list of groups analysed for H5 (squirrel monkeys) including gene information and BM/BSM result.
**Table S17** List of groups significant for H1 (robust capuchins) for the BM or BSM analyses including gene information and PAML results.
**Table S18** List of groups significant for H2 (gracile capuchins) for the BM or BSM analyses including gene information and PAML results.
**Table S19** List of groups significant for H3 (ancestral Cebinae) for the BM or BSM analyses including gene information and PAML results.
**Table S20** List of groups significant for H3a (across-Cebinae) for the BM analyses including gene information and PAML results.
**Table S21** List of groups significant for H4 (ancestral Cebidae) for the BM or BSM analyses including gene information and PAML results.
**Table S22** List of groups significant for H5 (squirrel monkeys) for the BM or BSM analyses including gene information and PAML results.
**Table S24** List of DAVID results for all enriched annotation categories for the H1 (robust capuchin) BM gene set including term description and ID, gene hits, and enrichment statistics.
**Table S25** List of DAVID results for all enriched annotation categories for the H1 (robust capuchin) BSM gene set including term description and ID, gene hits, and enrichment statistics.
**Table S26** List of DAVID GO clusters for the H1 (robust capuchin) BM gene set including cluster enrichment score (ES), term description and ID, gene hits, and enrichment statistics.
**Table S27** List of DAVID GO clusters for the H1 (robust capuchin) BSM gene set including cluster enrichment score (ES), term description and ID, gene hits, and enrichment statistics.
**Table S28** List of DAVID results for all enriched annotation categories for the H2 (gracile capuchin) BM gene set including term description and ID, gene hits, and enrichment statistics.
**Table S29** List of DAVID results for all enriched annotation categories for the H2 (gracile capuchin) BSM gene set including term description and ID, gene hits, and enrichment statistics.

**Table S30** List of DAVID GO clusters for the H2 (gracile capuchin) BM gene set including cluster enrichment score (ES), term description and ID, gene hits, and enrichment statistics.
**Table S31** List of DAVID results for all enriched annotation categories for the H3 (ancestral Cebinae) BM gene set including term description and ID, gene hits, and enrichment statistics.
**Table S32** List of DAVID results for all enriched annotation categories for the H3 (ancestral Cebinae) BSM gene set including term description and ID, gene hits, and enrichment statistics.
**Table S33** List of DAVID GO clusters for the H3 (ancestral Cebinae) BM gene set including cluster enrichment score (ES), term description and ID, gene hits, and enrichment statistics.
**Table S34** List of DAVID GO clusters for the H3 (ancestral Cebinae) BSM gene set including cluster enrichment score (ES), term description and ID, gene hits, and enrichment statistics.
**Table S35** List of DAVID results for all enriched annotation categories for the H3a (across-Cebinae) BM gene set including term description and ID, gene hits, and enrichment statistics.
**Table S36** List of DAVID GO clusters for the H3a (across-Cebinae) BM gene set including cluster enrichment score (ES), term description and ID, gene hits, and enrichment statistics.
**Table S37** List of DAVID results for all enriched annotation categories for the H4 (ancestral Cebidae) BM gene set including term description and ID, gene hits, and enrichment statistics.
**Table S38** List of DAVID results for all enriched annotation categories for the H4 (ancestral Cebidae) BSM gene set including term description and ID, gene hits, and enrichment statistics.
**Table S39** List of DAVID GO clusters for the H4 (ancestral Cebidae) BM gene set including cluster enrichment score (ES), term description and ID, gene hits, and enrichment statistics.
**Table S40** List of DAVID GO clusters for the H4 (ancestral Cebidae) BSM gene set including cluster enrichment score (ES), term description and ID, gene hits, and enrichment statistics.
**Table S41** List of DAVID results for all enriched annotation categories for the H5 (squirrel monkeys) BM gene set including term description and ID, gene hits, and enrichment statistics.
**Table S42** List of DAVID results for all enriched annotation categories for the H5 (squirrel monkeys) BSM gene set including term description and ID, gene hits, and enrichment statistics.
**Table S43** List of DAVID GO clusters for the H5 (squirrel monkeys) BM gene set including cluster enrichment score (ES), term description and ID, gene hits, and enrichment statistics.
**Table S44** List of DAVID GO clusters for the H5 (squirrel monkeys) BSM gene set including cluster enrichment score (ES), term description and ID, gene hits, and enrichment statistics.

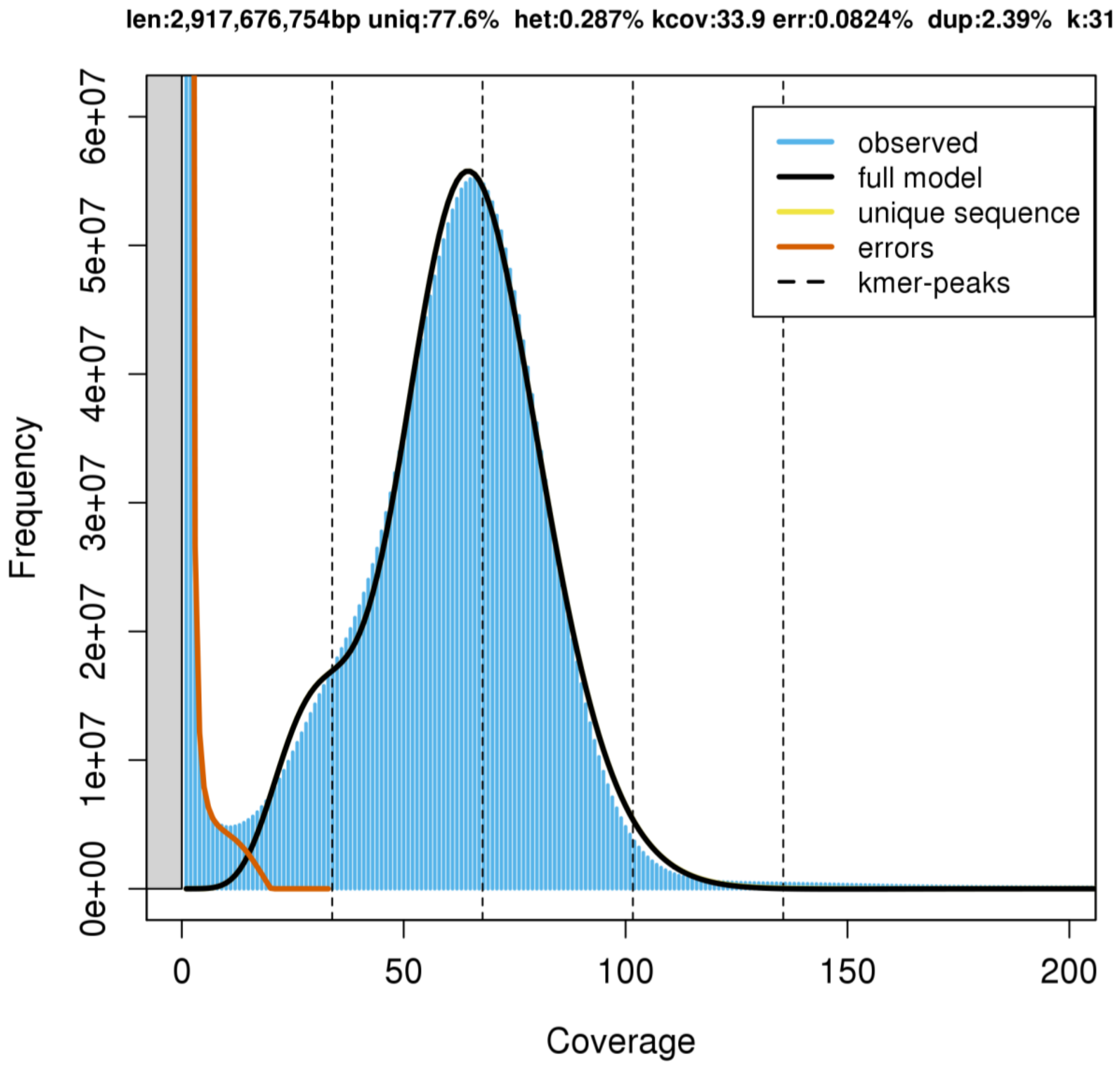


**Figure S1** Results of genome size estimation with GenomeScope showing estimated total length (len), percentage unique content (uniq), overall rate of heterozygosity (het), read error rate (err), and average rate of read duplications (dup).

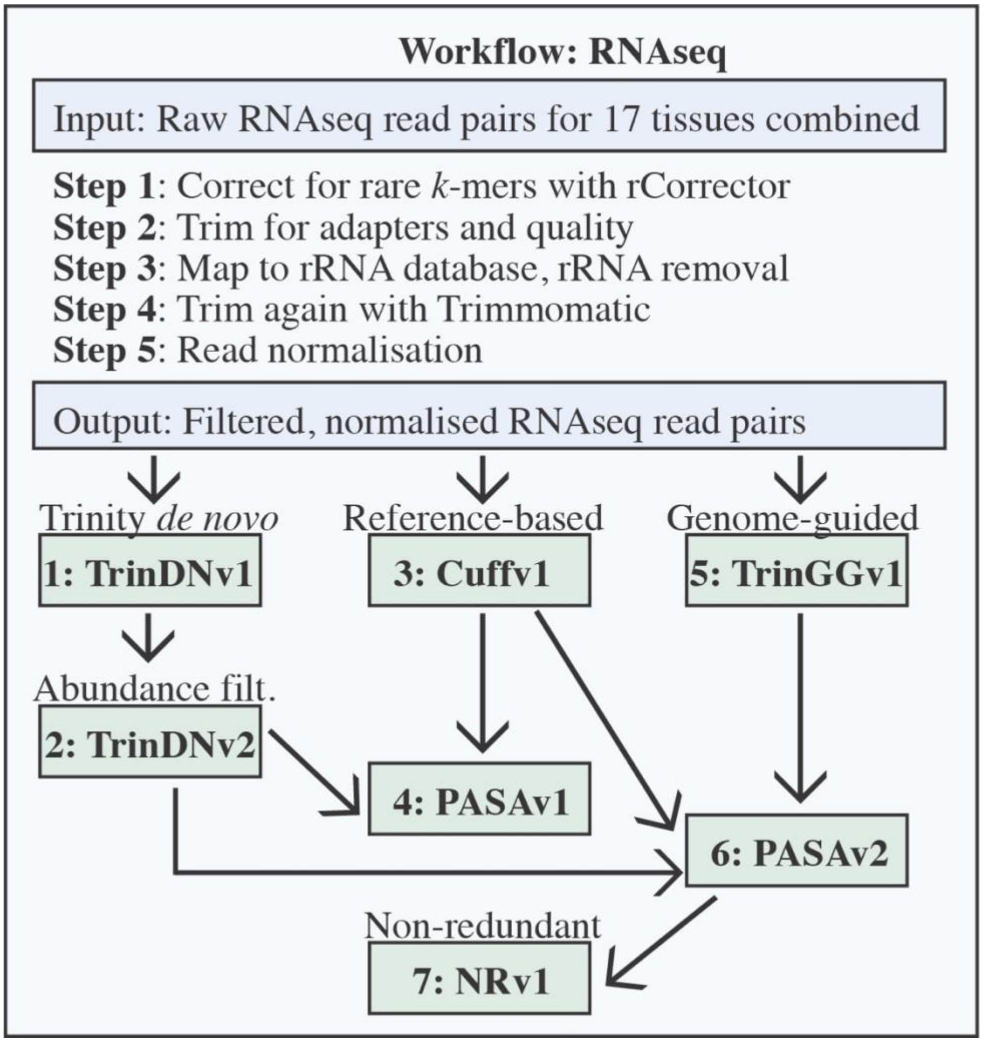


**Figure S2** Workflow summary graphic for the RNAseq filtering and assembly steps. Arrows indicate input for the various assemblies.

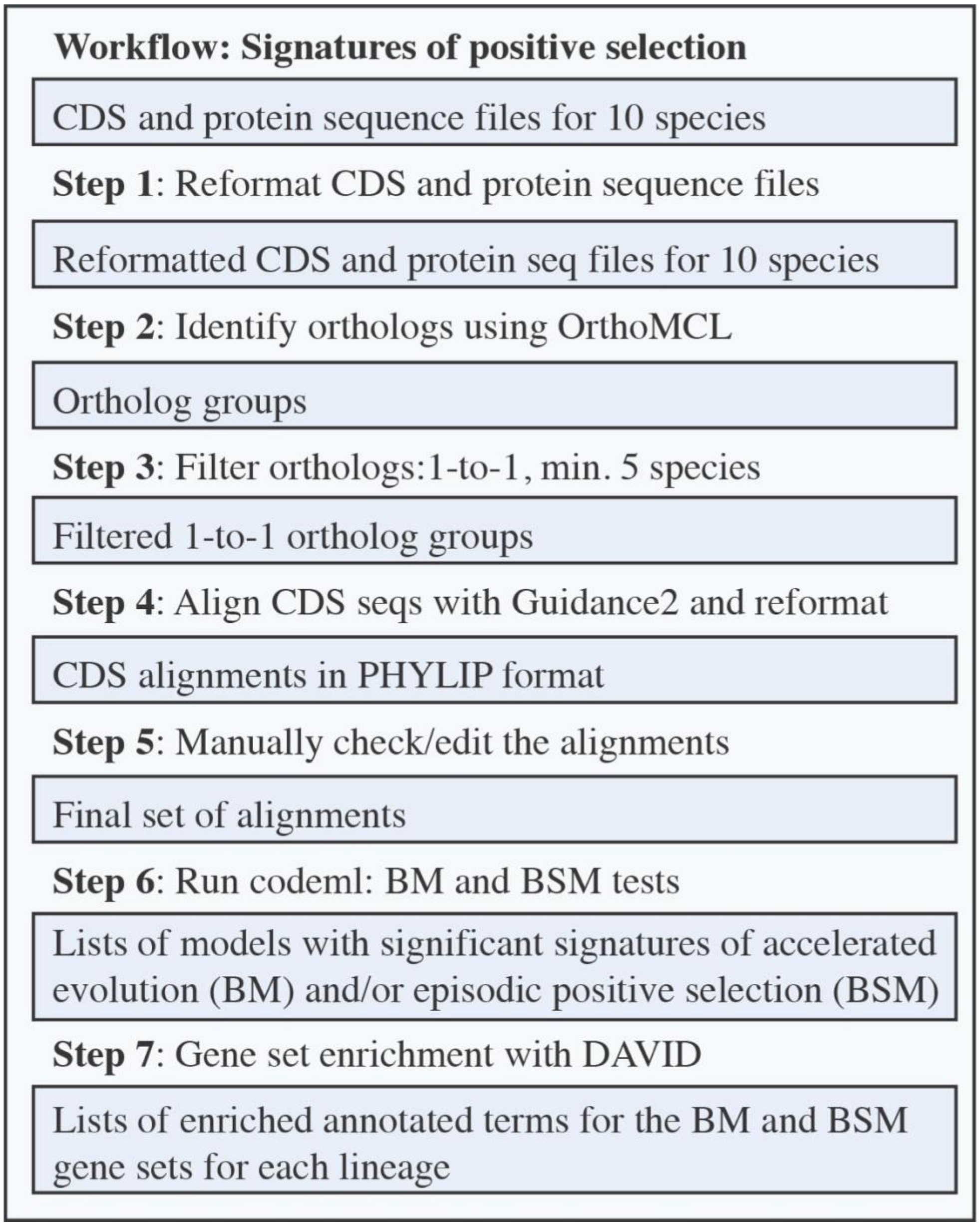


**Figure S3** A workflow summary graphic showing the input/output for the ortholog identification and alignment, codeml, and gene set enrichment analysis steps.
